# Coupling During Collective Cell Migration is Controlled by a Vinculin Mechanochemical Switch

**DOI:** 10.1101/2023.01.13.523997

**Authors:** T. Curtis Shoyer, Evan M. Gates, Jolene I. Cabe, Daniel E. Conway, Brenton D. Hoffman

## Abstract

Collective cell migration (CCM) plays important roles in development, physiological, and pathological processes. A key feature of CCM is the dynamic mechanical coupling between cells, which enables both long-range coordination and local rearrangements. This coupling requires the ability of cell adhesions to adapt to forces. Recent efforts have identified key proteins and implicated cellular-scale mechanical properties, but how key proteins give rise to these larger-scale mechanical processes is unclear. Using force-sensitive biosensors, cell migration assays, and molecular clutch models, we sought a molecular understanding of adhesion strengthening that could bridge this gap. We found that the mechanical linker protein vinculin bears substantial loads at AJs, FAs, and in the cytoplasm during epithelial sheet migration, and we identified a switch-like residue on vinculin that regulates its conformation and loading at the AJs during CCM. In vinculin KO-rescue, this switch jointly controlled the speed and coupling length-scale of CCM, which suggested changes in adhesion-based friction. To test this, we developed molecularly detailed friction clutch models of the FA and AJ. They show that open, loaded vinculin increases friction in adhesive structures, with larger affects observed in AJs. Thus, this work elucidates how load-bearing linker proteins can be regulated to alter mechanical properties of cells and enable rapid tuning of mechanical coupling in CCM.

## INTRODUCTION

The coordinated movements of groups of cells, termed collective cell migration (CCM), play important roles in many development, physiological and pathological processes, including tissue morphogenesis, wound healing, and the progression of cancer^1^. CCM is distinguished from single cell migration by the presence of adhesive contacts between cells. The types of cell-cell adhesion, and the associated coupling across many cells, are often used to define the various modes of CCM, which range from weakly-coupled neural crest cells undergoing streaming migration to the strongly-coupled epithelial cells undergoing sheet migration^2-4^. The differences between systems are thought to be due to expression of distinct sets of cell-adhesion receptors, such as the cadherin-switch associated with full and partial epithelial-mesenchymal transitions^3,5^. In contrast, the molecular-scale processes enabling rapid tuning of coupling within a given migration type are not as well-understood. This tuning is particularly important in the case of epithelial sheet migration, where the rapid alteration in coupling enables both the long-scale organization of large groups of cells while also permitting local cellular rearrangements required for efficient migration and avoidance of obstacles^6^.

Recent advances in the understanding of CCM have been driven by both screening-based approaches and mechanistic studies, which have identified key roles for many adhesive, scaffolding, and force-generating proteins, as well as physical models focusing on key cellular-scale mechanical properties, such as cell friction, polarity, and force-generation^7-9^. However, how these key proteins give rise to larger-scale mechanical processes is unclear. Interestingly, the process of adhesion strengthening, where force-application results in the strengthen of adhesion structures through the stabilization of key linkages and/or the recruitment of more linkages, has been implicated in both modeling and screening efforts.

We sought to determine if a molecular-scale, physical understanding of adhesion strengthening could elucidate the connections between key molecular players, cell-scale mechanical properties, and the regulation of epithelial cell coupling during CCM. To do so, we focused on the mechanical linker protein vinculin, as it is shown to be involved in CCM-associated processes, such as embryogenesis^10,11^. Furthermore, vinculin is also a key mediator of adhesion strengthening in two distinct ways. First, in response to force application, vinculin is recruited to the structures that link cells to the surrounding extracellular matrix (EMC), termed focal adhesions (FAs), as well as the structures that mediate linkages between neighboring cells, termed adherens junctions (AJs)^12-15^. Additionally, vinculin is amongst the strongest known catch-slip bonds, which exhibit increased binding lifetime in response to applied loads before eventually failing^16^.

Consistent with a role as a mediator of coupling during CCM, we find that vinculin bears substantial load at AJs, FAs, and throughout the cytoplasm during epithelial sheet migration. Furthermore, we identify a key residue, S1033, whose mutation affects the ability of vinculin to transition between inactive, unloadable and active, loadable states within the AJs and cytoplasm. Rescue of vinculn KO cells with WT, phosphomimetic (1033D), or unphosphorylatable (1033A) vinculin results in the modulation of cell speed and tuning of coupling, as measured by the length scale of correlated motion during CCM. Notably, these results are consistent with recent mechanical models of CCM, where variation in adhesion-based friction leads to covarying changes in cell speed and coordination. To assess the relationship between vinculin activation, vinculin load, and friction in adhesion structures, we created molecularly detailed frictional clutch models that relate force-sensitive binding dynamics of key components of AJs and FAs to the friction at each structure. Analyses reveal that increases in vinculin activation and load lead to increased friction, and these effects are stronger at AJs than FAs. Thus, this work reveals a novel regulatory switch that regulates the mechanical functions of vinculin to alter cell adhesion-based friction to enable the rapid tuning of coupling during CCM.

## RESULTS

### Vinculin is Loaded and Conformationally Open at the Edge of Collectively Migrating Cells

Key aspects of vinculin function are determined by the mechanical loads its experiences and its conformation^17^. Previous work in single cells has shown that load bearing and conformation regulation are separable^13^. Therefore, we sought to probe vinculin load and conformation during CCM. To do so, we developed a simple system of collective cell migration where the characteristics of vinculin loading and conformation could be readily observed. As has been done previously, we created radially expanding cell sheets using a cell droplet-based assay with Madin–Darby canine kidney (MDCK) epithelial cells (Extended Data Fig. 1a-c). As multiple MDCK strains have commonly been used in studies of epithelial dynamics^18^, we assessed both MDCK II and Parental MDCK cells. We verified that sheet expansion was primarily driven by migration, as reduction of cell proliferation with Actinomycin D caused no changes in dynamics (Extended Data Fig. 1d-e). To assess vinculin loading and conformation, we expressed either a FRET-based vinculin tension sensor (VinTS) or vinculin conformation sensor (VinCS) in each cell line^13,19^. All constructs produced stable proteins with the expected molecular weights (Extended Data Fig. 1f) and localized as expected to FAs and AJs in both cell types (Extended Data Fig. 1i, Fig. 1, and Extended Data Fig. 3). Over-expression of VinTS or VinCS did not alter migration dynamics or FA morphology in either cell line (Extended Data Fig. 1g-h). To interpret VinTS in this system, we verified that the cytosolic tension sensor module (TSMod) reported FRET efficiencies (∼0.29) consistent with no mechanical loading^20^ (Extended Data Fig. 2a-b). Similarly, to interpret VinCS, we established a reference for the closed state by measuring the FRET efficiency of VinCS in single cells non-specifically adhered to poly-L-lysine surfaces (Extended Data Fig 2c-d), a condition in which vinculin is predominantly cytosolic and unloaded^13^. Together, these data demonstrated that this system was sufficient for probing vinculin loading and conformation during collective cell migration.

**Fig. 1.**
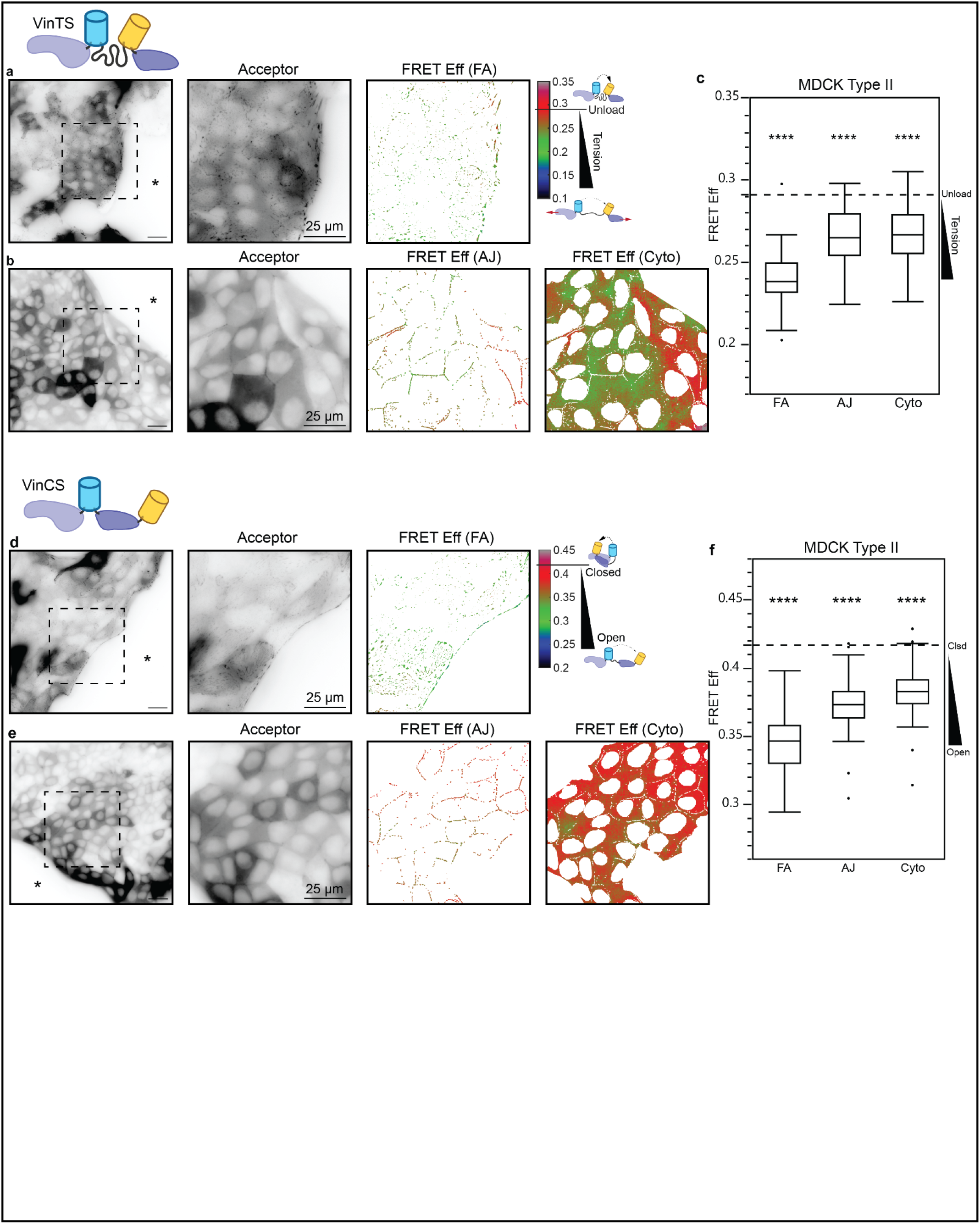
Vinculin is loaded and conformationally open at the edge of collectively migrating cells. (a) Representative image field of VinTS at the edge of migrating MDCK II cell monolayers in the basal plane with acceptor channel indicating sensor localization followed by zoom-in views of acceptor channel and FRET efficiency in the FA mask for the indicated region. Asterisk indicates free space adjacent to monolayer edge. (b) Representative image field of VinTS in the apical plane with acceptor channel followed by zoom-in views of acceptor channel and FRET efficiency in AJ and cytoplasm masks for the indicated region. (c) Box-whisker plot showing FRET efficiency for VinTS at FAs, AJs, and cytoplasm (n=43, 34, and 34 image fields respectively over at least 3 independent experiments). Differences between groups were detected using the Steel-Dwass test (****p < 0.0001). P-values shown are for comparisons to VinTS-I997A in MDCK II cells at the same structure (Extended Data Fig. 4c); p values for all comparisons can be found in Supplemental Note 2 Table S1. (d) Representative image field of VinCS at the edge of migrating MDCK II cell monolayers in the basal plane with acceptor channel indicating sensor localization followed by zoom-in views of acceptor channel and FRET efficiency in the FA mask for the indicated region. (e) Representative image field of VinCS in the apical plane with acceptor channel followed by zoom-in views of acceptor channel and FRET efficiency in AJ and cytoplasm masks for the indicated region. (f) Box-whisker plot showing FRET efficiency for VinCS at FAs, AJs, and cytoplasm (n=61, 51, and 52 image fields respectively over at least 3 independent experiments). Differences between groups were detected using the Steel-Dwass test (****p < 0.0001). P-values shown are for comparisons to the VinCS reference condition (Extended Data Fig. 2c); p values for all comparisons can be found in Supplemental Note 2 Table S2.

In confluent cells, vinculin exchanges between three sub-cellular compartments, the FAs, AJs, and cytoplasm, in a force-sensitive manner^21,22^. To probe the loads experienced by vinculin in these three compartments during CCM, we imaged VinTS-expressing MDCK cells at the leading edge of expanding epithelial sheets in the basal or apical plane. We employed standard image segmentation techniques to separate signals from the adhesion structures and cytoplasm in both focal planes. As the cytoplasmic signals were similar, we focused on the apical plane due to higher signal to noise in this compartment. Vinculin experiences the largest loads in the FAs, and lower, but substantial loading was observed in AJs and the cytoplasm in MDCK II (Fig. 1a-c) as well as Parental MDCK cells (Extended Data Fig. 3a-c). To determine if vinculin loading was dependent on interactions with F-actin, we used VinTS-I997A. This point mutation strongly disrupts actin binding while maintaining the ability of vinculin to undergo conformational regulation, and VinTS-I997A has been shown to not bear detectable loads in the FAs of single cells^23-25^. This resulted is the reduction of vinculin load in all compartments in both cell types (Extended Data Fig. 4), establishing that forces are transmitted through F-actin to vinculin in all compartments during CCM. Furthermore, we used STED imaging to confirm the existence of a cytoplasmic actin network in both MDCK cell types (Extended Data Fig. 5a-b). We also observed comparable loading of VinTS at the AJs and in the cytoplasm using confocal microscopy (Extended Data Fig. 5c-e), demonstrating that the VinTS signal at the AJs and in the cytoplasm were not due to out-of-plane optical effects from other sub-cellular structures.

As the exchange of vinculin between compartments is associated with conformation changes, we also probed vinculin conformation in the FAs, AJs, and cytoplasm during CCM using MDCK cells stably expressing VinCS and the sheet expansion assay. Vinculin was the most open in FAs, and smaller, but substantial, portions of vinculin was open in both the AJs and in the cytoplasm (Fig. 1d-f). FRET efficiency was highest in the cytoplasm but still significantly less than the closed reference, demonstrating the existence of a cytoplasmic population of open vinculin, which further supported a role for vinculin mediating mechanical connectivity in the cytoplasmic actin network. We found a similar trend in Parental MDCK cells (Extended Data Fig. 3d-f).

Taken together, the VinTS and VinCS data demonstrate that vinculin is loaded and conformationally open in the FAs, AJs, and the cytoplasm, but to varying degrees in each compartment. This suggests that vinculin facilitates differential mechanical connectivity of key load-bearing sub-cellular structures within and between cells during CCM.

### Vinculin S1033 mediates a regulatory switch that affects Vinculin Load and Conformation in Collectively Migrating Edges

Previous work has shown that the phosphorylation state of vinculin affects its mechanical functions^17^. Therefore, we sought to determine if vinculin phosphorylation contributed to distinct loading and conformation observed in the various sub-cellular compartments. Trends for vinculin loading at FA, AJ, and cytoplasm were similar for the two MDCK variants, so we performed these experiments with one variant, Parental MDCK. First, we verified that paraformaldehyde fixation did not affect the FRET efficiency of VinTS (Extended Data Fig. 5f), consistent with previous reports^20^. Use of VinCS in the same condition required a normalization approach, which yielded similar levels to live monolayers at all sub-cellular compartments (Extended Data Fig. 5g-h). Together, these data demonstrated that this system was sufficient for screening the effect of inhibitors and mutations on vinculin loading and conformation.

We first focused on how Src and Abl mediated phosphorylation of vinculin affected VinTS loading during CCM^21,26^. Inhibition of Src or Abl did not affect VinTS FRET efficiency at FAs, AJs, or the cytoplasm (Extended Data Fig. 6a-e). As phosphorylation at Y822 by Abl has been shown to affect vinculin mechanical function at the AJs of confluent epithelial cells, we also investigated the non-phosphorylatable point mutant (VinTS-Y822F)^21^. Consistent with inhibitor studies, VinTS and VinTS-Y882F exhibited identical localization and loading in collectively migrating Parental MDCK cells (Extended Data Fig. 6f-h).

Vinculin is also phosphorylated at S1033, and expression of non-phosphorylatable (S1033A) and phosphomimetic (S1033D) mutations affects the stiffness and traction force generation of fibroblasts^27^. To assess the effects of these mutations on vinculin loading during CCM, we incorporated these mutations in VinTS, creating VinTS-1033A and VinTS-1033D, stably expressed these sensors in Parental MDCKs, and performed sheet expansion assays. During CCM, both variants localized to FAs, AJs, and the cytoplasm. VinTS and the non-phosphorylatable VinTS-S1033A exhibit similar loading in all compartments (Fig. 2a-b,d and Extended Data Fig. 7a-b,d). In contrast, the phosphomimic VinTS-S1033D exhibited drastically increased FRET efficiency at the AJs and in the cytoplasm compared to VinTS (Fig. 2c-d), consistent with an apparent loss of loading. In FAs, VinTS-S1033D reported a partial loss of loading, suggesting a less-dominant regulatory role for 1033 phosphorylation in this compartment (Extended Data Fig. 7g-h).

**Fig. 2.**
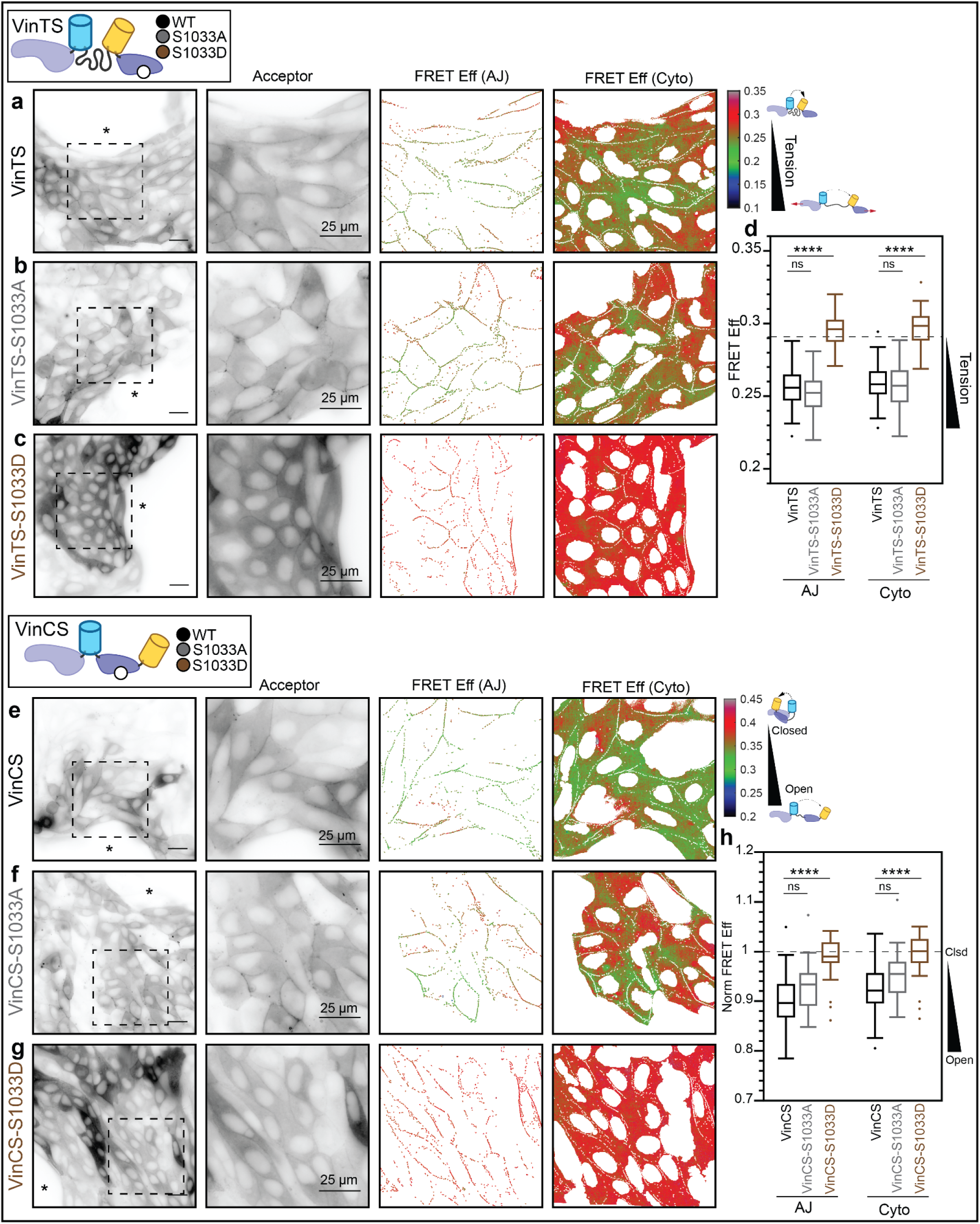
Vinculin S1033 mediates a regulatory switch that affects vinculin load and conformation at the edge of collectively migrating cells. (a-c) Representative image fields of VinTS, VinTS-S1033A, or VinTS-S1033D at the edge of migrating MDCK Parental cell monolayers in the apical plane with acceptor channel indicating sensor localization followed by zoom-in views of acceptor channel and FRET efficiency in AJ and cytoplasm masks for the indicated region. Asterisk indicates free space adjacent to monolayer edge. (d) Box-whisker plot showing FRET efficiency for VinTS, VinTS-S1033A, and VinTS-S1033D in AJs (n=61, 55, and 48 image fields respectively over at least 3 independent experiments) and cytoplasm (n=60, 55, and 48 image fields respectively over at least 3 independent experiments). (e-g) Representative image fields of VinCS, VinCS-S1033A, or VinCS-S1033D at the edge of MDCK Parental cell monolayers in the apical plane with acceptor channel indicating sensor localization followed by zoom-in views of acceptor channel and FRET efficiency in AJ and cytoplasm masks for the indicated region. (h) Box-whisker plot showing normalized FRET efficiency for VinCS, VinCS-S1033A, and VinCS-S1033D in AJs (n=58, 26, and 37 image fields respectively over at least 3 independent experiments) and cytoplasm (n=58, 26, and 37 image fields respectively over at least 3 independent experiments). Differences between groups were detected using the Steel-Dwass test (****p < 0.0001, ns not significant); p values for all comparisons can be found in Supplemental Note 2, Tables S4-5.

Vinculin phosphorylation is a potent regulator of vinculin conformation^17^. To determine if mutation of S1033 affects the conformation of vinculin, we created VinCS variants containing S1033A or S1033D, stably expressed them in Parental MDCK cells, and performed the sheet expansion assay. The VinCS and non-phosphorylatable VinCS-S1033A exhibited identical localization and conformation in all compartments conformation (Fig. 2e-f,h and Extended Data Fig. 7e-f,h). In contrast, in AJs and the cytoplasm, the phosphomimetic mutant (S1033D) exhibited drastically higher FRET, consistent with complete closing of vinculin (Fig. 2g-h). In FAs, VinCS-S1033D reported a reduction in the amount of open vinculin, consistent with a less-dominant regulatory role for 1033 phosphorylation in this compartment (Extended Data Fig. 7g-h).

Taken together, these data describe a regulatory switch for vinculin, where phosphorylation at S1033 biases vinculin towards a closed, unloaded state. Furthermore, this switch appears dominant at AJs and within the cytoplasm, but only partially reduces load and the amount of open vinculin in FAs. Thus, the switch mediates tuning of mechanical connectivity within and between cells, although with different strengths.

### Vinculin Regulatory Switch Affects Speed and Correlation Length of Collective Cell Migration

Next, we sought to determine the effects of this regulatory switch for vinculin on CCM. To do so, we first created a CRISPR-KO vinculin MDCK II cell line (Extended Data Fig. 8a) and rescued these cells with Vinculin-mVenus (VinV), Vinculin-mVenus-S1033A (VinV-S1033A), or Vinculin-mVenus-S1033D (VinV-S1033D). All constructs produced stable proteins with the expected molecular weights (Extended Data Fig. 8b). Furthermore, these variants localized to FAs, AJs, and cytoplasm as expected (Extended Data Fig. 8c-e) and exhibited similar trends in FA morphology as function of distance from the leading edge as (Extended Data Fig. 8f-g) as endogenous vinculin (Extended Data Fig. 1). Expression of VinV, VinV-S1033A, or VinV-1033D did not affect actin structures at the leading edge of monolayers (Extended Data Fig. 9a-f), and there was small (<∼25%) or non-significant differences in the abundance of E-cadherin (Extended Data Fig. 9g-k), alpha-Catenin (Extended Data Fig. 9l-p), or extended alpha-Catenin (Extended Data Fig. 9r-u) at the AJs across the four cell types. Together, these data demonstrated that this system was sufficient for testing the effects of the regulatory switch for vinculin on CCM.

To characterize CCM dynamics, we observed the migration of monolayers in a previously described barrier assay (Extended Data Fig. 10a-b) and measured velocity fields in the monolayer by optical flow constraint^28,29^ (Fig. 3a-d). To quantify effects on CCM, we used two well-studied kinematic parameters for migrating monolayers: the speed (average velocity magnitude) and the correlation length of deviations in the lateral velocity component, which is a previously described measure of mechanical coupling^30^. Rescue of MDCK II vinculin CRISPR-KO cells, with VinV reduced both the average speed (Fig. 3e), consistent with previous findings that vinculin knockdown increased speed^8^, and correlation length (Fig. 3f and Extended Data Fig. 10d). CCM of cells rescued with VinV-S1033A was comparable to those rescued with VinV, while rescue with VinV-S1033D was comparable to KO cells (Fig. 3f, Extended Data Fig. 10e). Together, these data show that the regulatory switch controlling vinculin loading and conformation determines the speed and correlation length of collectively migrating MDCK cells.

**Fig. 3.**
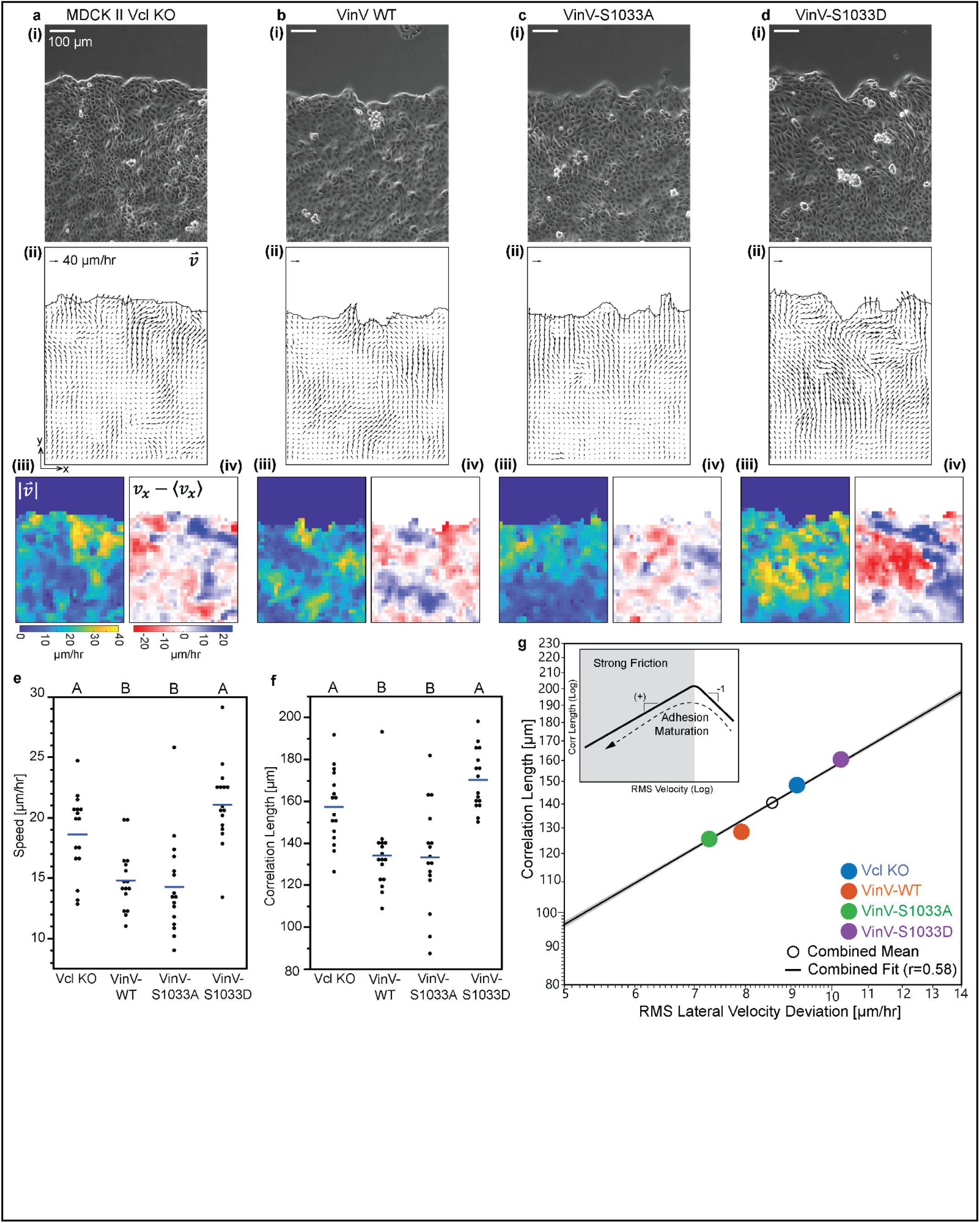
Vinculin Regulatory Switch Affects Speed and Correlation Length of Collective Cell Migration. (a-d) Representative image fields of MDCK II Vcl KO, VinV, VinV-S1033A, or VinV-S1033D cell monolayers in the barrier migration assay showing (i) phase contrast image, (ii) velocity field, (iii) velocity magnitude, and (iv) lateral velocity deviation. (e-f) Plots showing mean and all data points for velocity magnitude and correlation length for lateral velocity deviations (n=16 monolayers for each cell line over 6 independent experiments). Differences between groups were detected using Tukey’s HSD test. Levels not connected by the same letter are significantly different. (g) Log-log plot of correlation length versus RMS lateral velocity deviations for individual image fields and timepoints with mean for each cell line, combined mean, and fit of the combined data to log_10_(Y)=m*log_10_(X)+b with 95% confidence interval shaded. See Extended Data Fig 10 for data plotted separately by cell line.

### Effect of Vinculin Regulatory Switch in Models of Molecular Friction at the FA and AJ

We next leveraged recent modelling work to interpret the observed changes in CCM dynamics in terms of alterations in mechanical variables^31^. In the context of these models, a hallmark of strong adhesion-based friction in CCM is a positive relationship between correlation length and speed (Fig. 3g inset). The migration exhibited by CRISPR-KO vinculin and all the rescued MDCK II cells were found to be in this regime, both between experimental conditions (Fig. 3g) and within a given experimental conditions (Extended Data Fig. 10f). Furthermore, the lack of vinculin or the inability of vinculin to become open and loaded was consistent with a reduction in adhesion-based friction, suggesting that loaded vinculin enhances adhesion-based friction.

To probe the relationship between force-activated binding dynamics and adhesion-based friction, we used frictional clutch models (Supplementary Note 1), which predict the resistive force due to the sliding of two surfaces relative to each other at a particular speed as function of the number and properties of adhesive linkages between these surfaces^32^. To begin, we investigated previously developed models that contained linkages with a single simple bond type, such as an ideal bond that does not respond to force or a slip bond that weakens with force (Supplementary Note 1 Fig S2). To validate our implementation of a frictional clutch, we simulated clutches comprised of linkages with ideal or slip bonds, and we observed agreement with previous predictions of the relationship between frictional force and speed^32^. To gain intuition about the molecular determinants of friction, we also created a frictional clutch based on catch-slip bonds, and then compared the mean engagement lifetime and the effective friction coefficient, a standard parameter describing the frictional resistance between sliding surfaces (cell-ECM or cell-cell) in models of CCM for each scenario^33^. Notably, the qualitative shape of the friction coefficient-speed curve related to the individual linkage dynamics, being independent of speed for ideal bonds, monotonic decreasing for slip bonds, or biphasic for catch-slip bonds, indicating the underlying molecular-scale dynamics are indicative of the larger-scale mechanics of the frictional clutch.

To probe the effects of complex connectivity and potential regulatability of load-bearing linkages within adhesion structures, we developed multi-component linkages for use in the frictional clutch models. These multi-component linkages were based on integrin:talin:F-actin in FAs or E-cadherin:β-catenin:α-catenin:F-actin in AJs, which could be reinforced through the incorporation of vinculin. The ability these multi-component linkages to maintain connectivity under mechanical load was based on the force-dependent bond kinetics determined previously for key interfaces (Supplementary Note 1 Fig. S1). As these parameters were obtained from single molecule experiments characterizing the interfaces separately, we first assessed their suitability for use in combination to model multi-component linkages at the FA and AJ. The engagement lifetime of both linkages increased initially with loading rate and then decreased, indicating that the multi-component linkages possessed catch-slip characteristics, which were stronger for the integrin-based than cadherin-based linkage (Supplementary Note 1 Fig. S3). Furthermore, the reinforcement of the F-actin interface with vinculin did not change the overall functional form, but instead increased the lifetime of both FA and AJ linkages across a wide range of loading rates, as expected for a mechanical stabilizer. We note that these behaviors of the multi-component linkages are not readily predictable from the force-sensitive dynamics of single components, as there is no single dominant interface (Supplementary Note 1 Fig. S1).

Next, we determined the qualitative relationships between force-activated bond dynamics and larger-scale mechanics in FAs and AJs using frictional clutch models containing multi-component integrin- or cadherin-based linkages (Fig. 4a-b and Supplementary Note 1 Fig. S4-5). We represented the action of the S1033-based vinculin regulatory switch by modeling vinculin in two states. As mutation of S1033 affected vinculin load and conformation, but not localization to FAs/AJs, vinculin is conceptualized as either (1) closed and unloadable in FA/AJ (potentially bound to PIP2 or another unloaded component), or (2) open and loadable in FA/AJ (potentially bound to an exposed cryptic binding site in talin/α-catenin and F-actin) according to its binding kinetics. In the absence of vinculin, the mean linkage engagement lifetime varied biphasic with velocity (Fig. 4c,f), resembling the stronger/weaker catch-slip behaviors we had found for individual integrin-/cadherin-linkages (Supplementary Note 1 Fig. S2). Furthermore, the effective friction coefficient-velocity relationships were also biphasic. Thus, as was observed in the simpler frictional clutch models, the multi-component linkage dynamics were predictive of the qualitative shape of the friction coefficient-speed relationships (Fig. 4d,g). Furthermore, over the range of speeds associated with epithelial sheet migration (1-30 um/hr), increasing the fraction of loadable vinculin did not drastically change the functional forms of these, but did increase both the engagement lifetime and effective friction coefficient, consistent with a mechanical stabilizer. To assess the ability of the vinculin switch to tune friction, we assessed the effect of finer variations in the fraction of loadable vinculin at an intermediate speed in the range for CCM experimentally probed here (10 um/hr) (Fig.4e,d). In both FAs and AJs, the friction coefficient was tuned linearly by the amount of loadable vinculin, although vinculin’s effect was overall higher in AJs (∼4-fold increase) than in FAs (∼2-fold increase) (Fig.4e, h). Similarly, the ensemble vinculin tension scaled linearly with the amount of loadable vinculin, and the prediction of higher vinculin tension at FAs versus AJs was consistent with our experimental observations of lower VinTS FRET efficiencies (high tensions) in FAs compared to AJs.

**Fig. 4.**
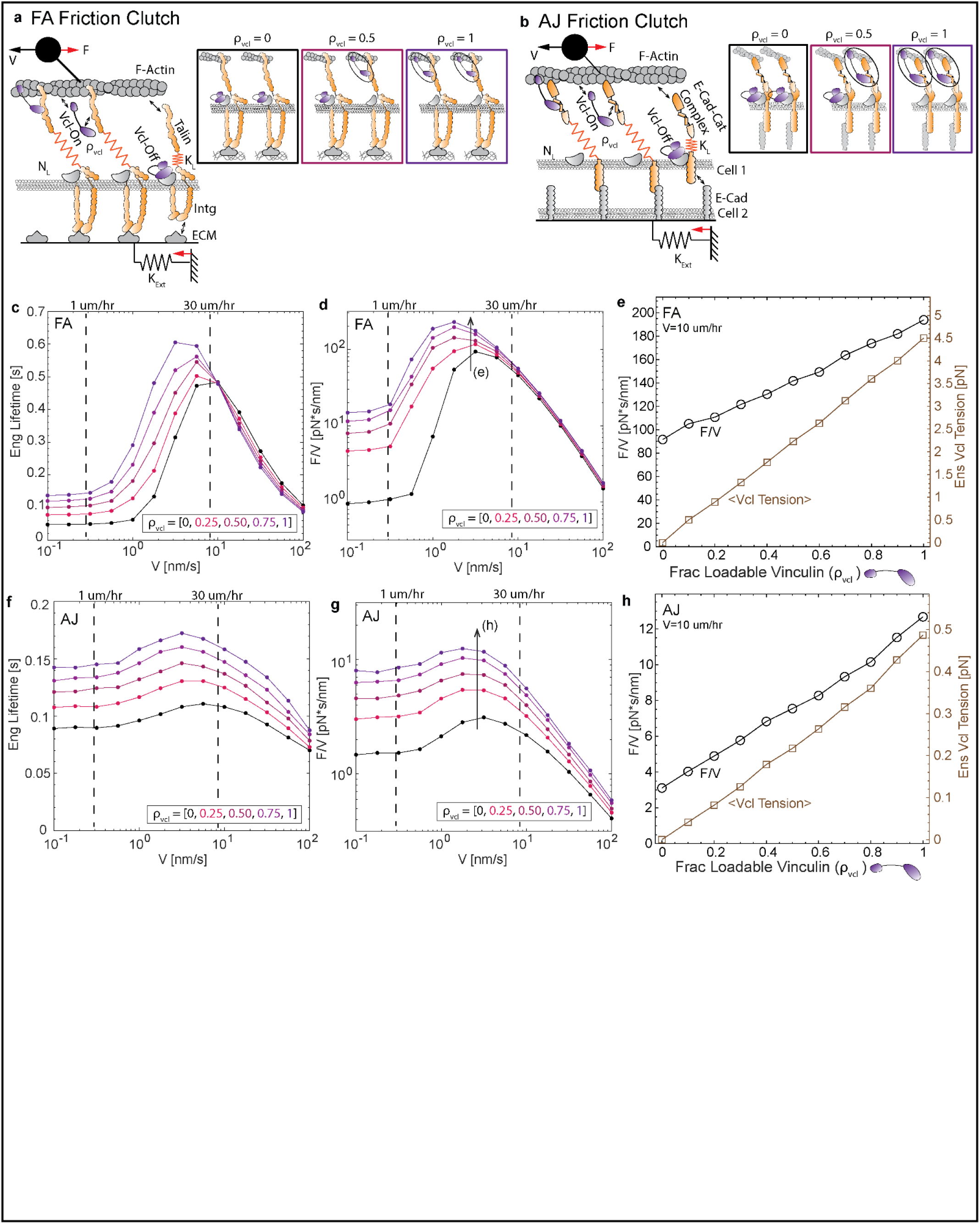
Effect of Vinculin Regulatory Switch in Models of Molecular Friction at the FA and AJ. (a-b) Schematics of FA and AJ friction clutch models. Linkage schematics depict different values for the fraction of linkages with loadable vinculin (*ρ*_*Vcl*_). (c-d) For FA friction clutch model, plots of mean linkage engagement lifetime and mean effective friction coefficient (*F/v*) versus speed (*v*) for 5 values of the fraction of loadable vinculin (*ρ*_*Vcl*_). (e) Plot of mean effective friction coefficient (left y-axis) and mean ensemble vinculin molecular tension (right y-axis) versus fraction of loadable vinculin for an intermediate speed (10 um/hr). (f-h) Analogous plots for the AJ friction clutch model. See Supplementary Note 1 for more information about the friction clutch models.

Together, these results suggest that regulation of vinculin mechanical reinforcement increases frictional drag strongly at the cell-cell interface, and to a lesser extent at cell-ECM interface, to affect the speed and coordination of collectively migrating epithelial sheets.

## DISCUSSION

Taken together, this work reveals a regulatory switch that determines the ability of vinculin to mediate mechanical connectivity within sub-cellular structures as well as the coupling between collectively migrating cells. The switch functions by toggling vinculin within AJs and the cytoplasm between closed, unloaded and open, loadable states. Reinforceable frictional clutch-based models based on force-sensitive binding dynamics of key components of AJs and FAs, show that the presence of open, loadable vinculin increases friction in adhesive structures, with larger affects observed in AJs. Previously, developed mechanical models of CCM predict that increases in adhesion-based friction drives a concomitant reduction in the speed and coupling length scale of collectively migrating cells, exactly as is observed when vinculin is locked into an open, loadable state through expression of non-phosphorylatable vinculin S1033A. Thus, this work elucidates how load-bearing proteins can be regulated to alter mechanical properties of cells to enable rapid tuning of mechanical coupling in CCM.

Several important open questions emerge from this work. First, vinculin is subject to a variety of other post-translational modifications^17^. For instance, previous work in confluent epithelial cells has shown that Abl-mediated phosphorylation of vinculin at Y822 was required for its localization to AJs^21^. This mechanism did not appear to be dominant in collectively migrating cells. Also, the S1033-based regulatory switch appeared dominant at AJs and with the cytoplasm, but had a smaller effect within FAs. These data suggest that an important question for future research is determining the kinases and phosphatases regulating vinculin in these diverse contexts, as well as their level of specificity for different compartments. Secondly, adhesion strengthening can occur through the recruitment of new linkages or the reinforcement of existing linkages. Previous work in single cells has focused on the recruitment of new linkages, especially integrins^34^. This work demonstrates the existence, tunability, and consequences of the reinforcement-based mechanism. Therefore, another important question is how these strengthening mechanisms interact and may be regulated in a coordinated or independent fashion.

Our work demonstrates a regulatory switch for vinculin that enables control of friction and modulation of cell coupling during migration. However, there are a plethora of mechanical linker proteins that localize to AJs and FAs to reinforce these structures and affect CCM^6,35-38^. Interestingly, we note that in our models of adhesion-based friction, we found that the vinculin reinforcement controlled the magnitude of linkage dynamics and frictional forces but had little effect on the functional form of these properties with respect to speed. This suggests independent control over the receptor specificity (achieved by E-cad or Integrin), functional form (achieved by the primary mechanical linker, E-cadherin:catenin complex or Intg:talin), and magnitude (achieved by secondary mechanical linker, vinculin) of force-dependent adhesion dynamics and mechanical force output of adhesion structures. Additionally, vinculin’s effect was speed-dependent, which was tied to the force-sensitivity of its actin bond. Thus, an attractive hypothesis is that the large number of linker proteins could enable precise and multi-factorial regulation of cell force transmission and dynamics in diverse processes, and that different force-sensitive dynamics could be optimized for certain processes based on the associated speeds at the given interface, whether controlled by relative cell motion or by actin dynamics. The framework developed here integrating biosensors to probe the state and molecular loads on a linker protein, mathematical models connecting force-sensitive bond dynamics of the linker protein to adhesion level force transmission, and tests of the function of the linker in collective migration provide a new means for determining the relative importance of different mechanical linker proteins in specific contexts, which will likely impact a variety of future studies in mechanobiology.

## Methods

### Generation of DNA constructs

Construction of pcDNA3.1-TSMod, pcDNA3.1-VinTS, pcDNA3.1-VinV, pcDNA3.1-VinCS, and pcDNA3.1-VinTS-I997A have been described previously ^13,24^. PCR mutagenesis was used to generate DNA constructs for the mutant of VinTS that is deficient in Y822 phosphorylation (pcDNA3.1-VinTS-Y822F), mutants of VinV, VinTS, and VinCS that are deficient in S1033 phosphorylation (pcDNA3.1-VinV-S1033A, pcDNA3.1-VinTS-S1033A, and pcDNA3.1-VinCS-S1033A), and mutants of VinV, VinTS, and VinCS that mimic phosphorylated S1033 (pcDNA3.1-VinV-S1033D, pcDNA3.1-VinTS-S1033D, and pcDNA3.1-VinCS-S1033D). For the Y822F mutant, forward primer 5’-TTGGATTCTGGATTCAGGATTCTGGG-3’, reverse primer 5’-CCCAGAATCCTGAATCCAGAATCCAA-3’, and template DNA pcDNA3.1-VinTS were used. For the S1033A mutants, forward primer 5’-AACCTCATGCAGGCTGTGAAGGAAACT-3’, reverse primer 5’-CTGGGCGTTATGAACCAACATCTCAG-3’, and template DNA pcDNA3.1-VinV, pcDNA3.1-VinTS, or pcDNA3.1-VinCS were used. For the S1033D mutants, forward primer 5’-AACCTCATGCAGGATGTGAAGGAAACT-3’, reverse primer 5’-CTGGGCGTTATGAACCAACATCTCAG-3’, and template pcDNA3.1-VinV, DNA pcDNA3.1-VinTS, or pcDNA3.1-VinCS were used. To create plasmids for lentiviral expression of these constructs, pcDNA3.1 plasmids were digested with NruI/XbaI and ligated into pRRL vector that had been digested with EcoRV/XbaI. All newly generated constructs were verified by DNA sequencing (Genewiz).

### Cell Culture and Expression of DNA constructs

MDCK Parental cells (ATCC® CCL-34™, obtained from Duke University Health System’s Cell Culture Facility) and MDCK II cells (generous gift from Dr. Adam Kwiatkowski, University of Pittsburgh) were maintained in a humidified 5% CO2 atmosphere at 37°C in DMEM-LG (D6046; Sigma Aldrich) supplemented with 10% fetal bovine serum (HyClone), 1% antibiotic/antimycotic (Gibco), and 1 g/L sodium bicarbonate (Gibco). MDCK cell type was confirmed by probing expression for Claudin-2 (expressed by MDCK II cells but not MDCK Parental cells, data not shown).

CRISPR/Cas9-mediated knockout of vinculin in MDCK II cells was performed using a previously described guide RNA^39^. Knockout of vinculin was confirmed by western blot analysis.

HEK293-T cells, used for viral production, were maintained in DMEM-HG (D5796; Sigma Aldrich) supplemented with 10% fetal bovine serum (HyClone) and 1% antibiotic/antimycotic (Gibco). For viral transduction, the second generation viral packaging plasmids psPax2 (Plasmid #12260) and pMD2.G (Plasmid #12259) were purchased from Addgene. To generate viral particles containing the DNA for a desired construct, the corresponding pRRL-based construct, psPax2, and pMD2G plasmids were transfected into HEK293-T cells using Lipofectamine 2000 (Invitrogen) according to the manufacturer’s protocol. After 4 hours, the transfection mixture was exchanged for complete growth media. After an additional 72 hours, media containing viral particles was harvested and stored at -80°C. One day prior to viral transduction, MDCK cells were plated in 6-well dishes at a density of approximately 100,000 cells per dish. Cells were transduced with 500 µL of viral particle containing growth media supplemented with 2 µg/mL Polybrene (Sigma Aldrich) to enhance viral uptake. After three passages, transduced cells were sorted via fluorescence activated cell sorting (FACS) based on the intensity of the fluorescent signal of the construct. For expression of FRET sensors in Parental MDCK and MDCK II cells, expression levels were selected that yielded sufficient signal-to-noise in FRET measurements and did not affect cell migration or FA morphology. For rescue of MDCK II Vcl KO cells with VinV constructs, expression levels were selected that localized to FAs, AJs, and cytoplasm as expected.

### Droplet-based Migration Assay

To create cell adherent surfaces appropriate for imaging, glass bottom dishes (World Precision Instruments) or no. 1.5 glass coverslips mounted in reusable metal dishes (Bioptechs, Butler, PA) were incubated with 10 µg/ml fibronectin (Fisher Scientific) in PBS at room temperature for 1 hour, rinsed once with PBS, and allowed to dry prior to cell seeding. Following a previously published protocol to create collectively migrating cell islands^40^, approximately 5×10^3^ cells suspended in 5 μL growth medium were plated as a droplet on the fibronectin-coated glass bottom cell culture dishes. Cells adhered for 30 minutes at 37°C and 5% CO_2_, and then the dish was filled with 2 mL growth medium. Cells were incubated for 72 hours at 37°C and 5% CO_2_ to enable formation of a mechanically integrated and collectively migrating cellular layer. In the droplet assay, all experiments were conducted at 72 hr post-seeding, except for area expansion, which was quantified between 48 and 72 hr post-seeding.

To quantify migration in the droplet assay, cell islands were imaged at 48 and 72hrs using phase microscopy with a 10x objective (UPlan FLN/NA0.3 10x Objective, Olympus) on an Olympus inverted fluorescent microscope (Olympus IX83, Tokyo, Japan) equipped with a sCMOS ORCA-Flash4.0 V2 camera (Hamamatsu Photonics, Hamamatsu, Japan). After establishing Köhler illumination, a fixed grid of images was acquired using Metamorph acquisition software (Olympus). For each sample, images were stitched together using the ImageJ Grid/Collection stitching plugin. Island size was then manually measured in ImageJ. Briefly, background was subtracted using the Subtract Background tool. The rolling ball radius was set to 75 pixels, and the resultant image was converted to a binary mask using the associated built-in function. To determine the island’s expansion over 24hrs, the change in area from 48 to 72hrs was normalized by the initial island size at 48 hrs.

To assess the effect of proliferation in the droplet assay, monolayer expansion was compared in the absence or presence of the proliferation inhibitor Actinomycin D. Immediately following the imaging at 48 hrs post-seeding, cells were treated with Actinomycin D (Sigma, Product SBR00013) at a concentration of 2 ng/mL for a duration of 8 hr. To assess levels of cell proliferation, Click-iT™ Plus EdU Alexa Fluor™ 647 Imaging Kit (Fisher Scientific) was used. At 64hrs post-seeding, cells were treated with 10µM EdU for 8 hours before fixation. Detection of EdU was performed per the manufacturer’s protocol.

### Barrier-based Migration Assay

To prepare the surface, 12-well glass bottom plates (Cellvis) were incubated with 10 µg/ml fibronectin (Fisher Scientific) in PBS at room temperature for 1 hour, rinsed once with PBS, and allowed to dry prior to cell seeding. Barrier molds (iBidi) were positioned and adhered to the 12-well glass bottom plate using a custom alignment tool. A 70uL suspension of cells was seeded at a density of 500 cells/µL into one chamber in a barrier mold. Cells grew for 14.5 hours, forming a confluent monolayer inside the barrier. Then, the barrier was lifted, at which point the cells were able to migrate into free space. After barrier removal, cells were washed immediately with PBS once and then provided media. In the barrier assay, migration kinematics and actin organization were assessed.

To quantify migration in the barrier assay, timelapse multifield imaging of migration in the barrier assay was performed using phase contrast microscopy on a Zeiss Axio Observer Z1 microscope outfitted with a Pecon XL S1 incubator regulating temperature (37°C), CO_2_ concentration (5%), and humidity. The following objective was used: 10x/0.30 Plan-NeoFluar Ph1, (440331-9902) WD: 5.2mm. Movement of the sample (motorized XY stage), image acquisition (Photometrics CoolSNAP HQ2 CCD camera), and software-based autofocus were computer-controlled using Metamorph (Olympus) software. Imaging was started approximately 3 hours post-barrier lift and conducted for a duration of approximately 6 hours. The delay between two successive images of the same field was 10 minutes. For each monolayer, a minimum of 4 fields of view located along the longer free edge of the rectangular monolayer were monitored.

MATLAB (Mathworks) was used for all image analysis. Velocity fields were computed from the timelapse images using a previous implementation of the Optical Flow Constraint method from Vig, et al. ^29^.

Velocities were computed on a square grid 32 px (20.8 µm) apart at all positions inside the monolayer in the field of view. To validate velocity field computation, we simulated the motion of artificial particles subjected to the computed velocity field and overlaid the positions onto the original timelapse movie, as previously described^29^. To quantify migration kinematics, speed and spatial correlation length were computed for each field of view and then averaged to obtain a single value for each monolayer. Speed was defined as the magnitude of the velocity vector averaged over grid points located less than or equal to 500 µm from the leading edge, as given below:

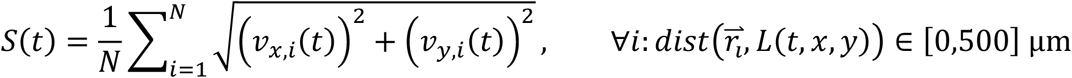

where 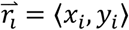 is the position of grid point *i*, 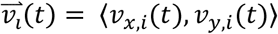 is the velocity of grid point *i* at timepoint *t, L*(*t, x, y*) is the curve representing the leading edge at timepoint *t*, and *dist*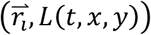 is the minimum distance between the grid position 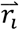 and the leading edge curve *L*(*t, x, y*). The fixed coordinate system is defined such that the y-direction is normal to the free edge created by the barrier mold, and the x-direction is parallel to it. For each field of view, a time-averaged leading edge speed was then obtained. Furthermore, as a measure of spatial correlation in the velocity field, we used the correlation length for lateral velocity deviations, as previously described^30^. The lateral velocity deviation for grid point *i* at timepoint *t, u*_*i*_(*t*), is defined as the x component of the velocity at the grid location minus the average of x velocity components over all grid locations in the monolayer in the field of view, given by: *u*_*i*_(*t*) = *v*_*x,i*_(*t*) *−* ⟨*v*_*x*_(*t*)⟩. The normalized spatial correlation coefficient as a function of radial distance *r* at time *t, C*(*r, t*), was then computed using the following equation:

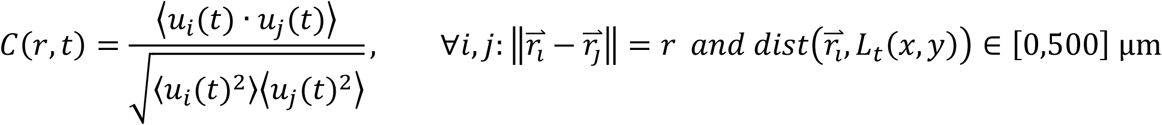

Computation was performed over all grid points *i* located less than or equal to 500 µm from the leading edge, which includes most of the monolayer but keeps a constant computation window for all fields of view, and radial distances were binned in 10 µm bins. For each field of view, a single correlation length was determined by plotting the time-averaged normalized correlation coefficient, *C*(*r*) = ⟨*C*(*r, t*)⟩, versus distance *r* and determining the smallest value for *r* such that the correlation function decays below a threshold of 0.1. To assess relationships between speed and correlation length, the root-mean-square lateral velocity deviation and correlation length in lateral velocity deviation from individual timepoints were used.

### Western Blot Analysis

Cells were washed one in ice-cold PBS buffer and lysed in ice-cold lysis buffer [10% Glycerol, 2 mM EDTA, 250 mM NaCl, 50mM HEPES, 0.5% NP-40, protease inhibitor cocktail (Sigma)]. Cell lysates were centrifuged for 10 minutes at 13000 RPM and 4°C. Supernatants were separated and pellets of cell debris were discarded. Afterwards, 2x Laemmli sample buffer (Bio-Rad Laboratories) was added to the lysate for a 1:1 dilution and the sample was boiled at 100 °C for 5 minutes. Samples were then loaded into Mini-PROTEAN® TGX™ Precast Gels (4-20%, Biorad) and ran at 100 V for 70 minutes before being transferred to a PVDF membrane (Bio-Rad Laboratories) via wet-transfer. Membranes were blocked with 5% dry milk in TBST [10 mM Tris-HCl, 100 mM NaCl, 0.1% Tween 20] for 1 hour at room temperature and then incubated with primary antibodies per the dilution listed overnight at 4 °C. Afterwards, the membrane was rinsed 3 times in TBST and incubated with the appropriate enzyme conjugated secondary antibody (Life Technologies), depending on the animal species, for 1 hour at room temperature. Membranes were then rinsed again 3 times in TBST and then developed using Supersignal West Pico Chemiluminescent Substrate (Thermo Fisher Scientific). The signal was detected either on X-ray film (Kodak) or by imaging (ChemiDoc Imaging System, Bio-Rad Laboratories). Primary and secondary antibodies used for Western Blots are provided in Table 1.

**Table 1:** Primary and Secondary Antibodies for Western Blot Analysis.

| Type | Antibody | Species | Clonality | Manufacturer | Product | Dilution |
| --- | --- | --- | --- | --- | --- | --- |
| Primary | Vinculin | Mouse | monoclonal | Sigma | V91314 | 1:8000 |
| Primary | GAPDH | Rabbit | polyclonal | Santa Cruz | sc25778 | 1:4000 |
| Primary | GFP | Rabbit | polyclonal | Abcam | ab6556 | 1:5000 |
| Secondary | Anti-mouse IgG (H+L), Cross- | Goat | polyclonal | Thermo Fisher | G21040 | 1:3000 – 1:5000 |
|  | Adsorbed<br>Secondary<br>Antibody, HRP |  |  |  |  |  |
| Secondary | Anti-rabbit IgG<br>(H+L) , Cross-Adsorbed<br>Secondary<br>Antibody, HRP | Goat | polyclonal | Thermo Fisher | G21234 | 1:3000 –<br>1:5000 |

### Fixation & Immunofluorescent Staining

For fixation or immunofluorescent labeling, cells were washed once with PBS (containing Ca^2+^ and Mg^2+^), fixed with 4% methanol-free (EM grade) paraformaldehyde (Electron Microscopy Sciences, Hatfield, PA) for 10 minutes and then rinsed with PBS. For immunofluorescent labeling, cells were treated with 0.1% Triton-X for 15 min and then rinsed with PBS. Fresh 2% bovine serum albumin (BSA, Sigma Aldrich) in PBS was used as blocking buffer for 30 min. Primary antibody was applied for 60 min and then rinsed three times with PBS. Cells were again blocked for 30 min. Secondary antibody was applied for 60 min. Cells were then rinsed three times with PBS and imaged in PBS. Primary antibodies and the dilutions used for immunofluorescent labeling are provided in Table 2. Secondary antibodies raised against the appropriate primary species and conjugated with dyes, including Alexa Fluor 488, 594, and 647, were purchased from Thermo Fisher and used at a dilution of 1:500. To label actin, cells were treated with Alexa Fluor 488-, 594-, or 647-conjugated phalloidin (Invitrogen) at a 1:100 dilution during the secondary antibody step. To label nuclei, after fixation and permeabilization, cells were rinsed in PBS and stained with a 1:5000 dilution of 4’, 6-diamidino-2-phenylindole, dihydrochloride (DAPI, Life Technologies, D1306) in PBS for 5 min at room temperature then rinsed twice in PBS.

**Table 2:** Primary Antibodies for Immunofluorescent Staining.

| Antibody | Species | Clonality | Manufacturer | Product | Dilution |
| --- | --- | --- | --- | --- | --- |
| vinculin | Mouse | monoclonal | Sigma | V91314 | 1:500 |
| E-cadherin | Rat | monoclonal | Sigma | U3254 | 1:500 |
| α-catenin | Rabbit | polyclonal | Cell Signaling | 3236 | 1:200 |
| α-catenin<br>extended<br>conformation-sensitive<br>antibody (α18) | Rat | monoclonal | Nagafuchi Lab* | N/A | 1:1000 |

\*Generous gift of Dr. Akira Nagafuchi (Nara Medical University)

### Imaging of FRET-based Sensors and Immunofluorescence

An Olympus inverted fluorescent microscope (Olympus IX83, Tokyo, Japan) was used to image samples. Images were acquired at 60x magnification (UPlanSApo 60X/NA1.35 Objective, Olympus) and illuminated by a LambdaLS equipped with a 300W ozone-free xenon bulb (Sutter Instrument, Novato, CA). The images were captured using a sCMOS ORCA-Flash4.0 V2 camera (Hamamatsu Photonics, Hamamatsu, Japan). The FRET images were acquired using a custom filter set comprised of an mTFP1 excitation filter (ET450/30x; Chroma Technology Corp, Bellows Falls, VT), a mTFP1 emission filter (FF02-485/20-25, Semrock, Rochester, NY), Venus excitation filter (ET514/10x; Chroma Technology Corp), Venus emission filter (FF01-571/72; Semrock) and dichroic mirror (T450/514rpc; Chroma Technology Corp). For sensitized emission FRET microscopy, three images are acquired to calculate FRET efficiency^41^.

These include imaging the acceptor (Venus excitation, Venus emission), FRET (mTFP1 excitation, Venus emission), and donor (mTFP1 excitation, mTFP1 emission). Exposure times for imaging of Venus, Teal-Venus FRET, and Teal were 1000ms, 1500ms, and 1500ms, respectively. To avoid photobleaching, only one image was taken per cellular region, either at the basal (FA’s) or apical (AJ’s) focal plane. For immunofluorescent imaging, we utilized the DA/FI/TR/Cy5-4X4 M-C Brightline Sedat filter set (Semrock) and the associated dichroic mirror (FF410/504/582/669-Di01). The motorized filter wheels (Lambda 10-3; Sutter Instrument), automated stage (H117EIX3; Prior Scientific, Rockland, MA), and image acquisition were controlled through MetaMorph Advanced software (Olympus). For live cell imaging, growth media was replaced with live cell visualization media (Sapphire North America, Ann Arbor, MI, MC102), supplemented with 10% FBS and 1 g/L sodium bicarbonate, 30 minutes before imaging. A constant temperature was maintained across the sample using an objective heater (Bioptechs, Butler, PA 150819-13) in conjunction with a stage and lid heater (Bioptechs Stable Z System 403-1926). A humidified CO_2_ perfusion system (Bioptechs 130708) was used to maintain a stable pH. All components were brought to equilibrium prior to imaging.

### Calculation of FRET Efficiency from Sensitized Emission

FRET was detected through measurement of sensitized emission^42^ and calculated using custom written code in MATLAB (Mathworks)^41^. All analyses were conducted on a pixel-by-pixel basis. Prior to FRET calculations, all images were first corrected for dark current, uneven illumination, background intensity, and three-dimensional offsets caused by chromatic aberrations and minute hardware misalignments (registration) as previously described ^20^. Spectral bleed-through coefficients were determined through FRET-imaging of cells expressing only donor or only acceptor FP. The donor bleed-through coefficient (dbt) was calculated for mTFP1 as:

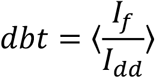

where *I*_*f*_ is the intensity in the FRET-channel, *I*_*dd*_ is the intensity in the donor-channel, and data were binned by donor-channel intensity. Similarly, the acceptor bleed-through coefficient (abt) was calculated for Venus (A206K) as:

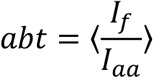

where *I*_*aa*_ is the intensity in the acceptor-channel, and data were binned by acceptor-channel intensity. For the mTFP1-Venus (A206K) FP pair on our microscope setup, the cross-talk between donor and acceptor channels (signal from donor in acceptor channel and vice-versa) was determined to be negligeable. To correct for spectral bleedthrough in experimental data, pixel-by-pixel FRET corrections were performed according to the equation:

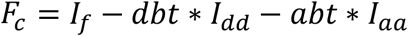

where *F*_*c*_ is the corrected FRET image, *I*_*f*_ is the intensity in the FRET-channel, *I*_*dd*_ is the intensity in the donor-channel, and *I*_*aa*_ is the intensity in the acceptor-channel. After bleed-through correction, FRET efficiency was calculated. Through imaging donor-acceptor fusion constructs of differing, but constant, FRET efficiencies, it is possible to calculate two proportionality constants that enable the calculation of FRET efficiencies for any single-chain biosensor ^42^. These proportionality constants are G:

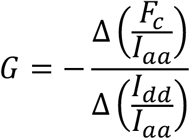

where Δ indicates the change between two donor-acceptor fusion proteins, and k:

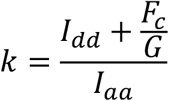

Using published methods^20^, the calibration factors were experimentally determined for mTFP1 and Venus (A206K). With these two proportionality constants, it is possible to calculate both FRET efficiency (E):

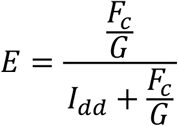

and the relative concentration of donor and acceptor fluorescent proteins [D]/[A] (or DPA) in a sample:

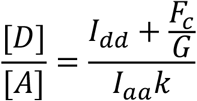

The calibration constants G and k were monitored over the course of this work to control for changes in lamp and filter performance.

Segmentation routines were used to quantitate FRET efficiencies within FA’s (for images of the basal focal plane) or AJ’s and Cytosol (for images of the apical focal plane). All operations were conducted on the acceptor channel, which is independent of FRET and proportional to the concentration of VinTS or VinCS. Segmentation of FAs was done as previously reported using a water-based algorithm ^43^. Briefly, for AJ segmentation, edge detection on the acceptor image was conducted using a dispersive phase stretch transform^44^. The resultant edge detection was then high-pass filtered and a user-defined intensity threshold was used to eliminate background signal and isolate the AJs. For basal and apical images, binary masks containing all FA’s or all AJ’s, respectively, were applied to FRET efficiency images. Cytosolic signal was examined in all apical images by inverting the AJ mask and then removing nuclear regions from the cytosolic mask via local normalization followed by morphological processing. After segmentation, closed boundaries were manually drawn by the user based on the unmasked acceptor channel image to include regions with appropriate and uniform expression of the sensor. Information outside manual boundaries was discarded. For each image, mean acceptor intensity, FRET efficiency, and donor-per-acceptor ratio were characterized.

To ensure the quality of FRET data, a multi-scale filtering approach was used. In comparing samples, the same filtering approach was applied to each population of data points. First, at the pixel level, regions detected as having DPA outside of the 0.5-2.0 range, were discarded. Then, at the image level, images with less than 2000 px in the analysis region were discarded. Additionally images were discarded if >33% of the pixels were removed for out-of-bounds DPA. Overall, the pixel-level filtering process removed less than 5% of pixels on average (across all constructs and all subcellular structures), and the image-level filtering process removed less than 3% of images.

### Kinase Inhibitor Treatments

Inhibitor treatments were performed on cells in the droplet assay 72 hr post-seeding. To inhibit Src kinase, cells were treated with 10 μM PP2 (Abcam ab120308) for 1 hour. To inhibit Abl kinase, cells were treated with 50 μM Imatinib (Sigma SML1027) for 1 hour. After treatment, cells were washed, fixed, and imaged as previously described.

### Quantification of Actin Organization at the Edge of Migrating Monolayers

Actin organization was assessed using phalloidin labeling of actin (see Fixation & Immunofluorescent Staining) in the droplet assay 72 hr post-seeding. Lamellipodia and actin belts (of continuous contour length greater than 75µm) were identified manually.

### Quantification of Abundance of Proteins at AJs in Migrating Monolayers

The abundance of proteins at the AJ were assessed using immunofluorescent labeling (see Fixation & Immunofluorescent Staining) in the droplet assay 72 hr post-seeding. As with FRET imaging, all images were corrected for dark current, uneven illumination, background intensity, and three-dimensional offsets caused by chromatic aberrations and minute hardware misalignments (registration). Then, binary masks of the AJs were generated via high-pass filtering of the immunofluorescent channel using custom MATLAB software and applied to images to obtain average intensities for each image. To account for day-to-day variability in immunofluorescent staining, the immunofluorescent signals were each normalized by day.

### Quantitation of Vinculin FA Morphology

To quantify vinculin focal adhesion (FA) morphology, images of immunofluorescent-labeled vinculin or the accepter channel of VinCS or VinTS were used. Focal adhesion segmentation was performed as described above using the signal of immunofluorescent-labeled vinculin or the acceptor channel of VinCS or VinTS. For each FA, the distance from its center to the nearest point on a manually drawn leading edge was determined. The orientation of the FA was defined as the angle between the major axis of the ellipse fit to the FA and the normal direction of the leading edge at the nearest point to the FA. As such, 0° indicates the FA is parallel to the migration direction, and 90° indicates the FA is perpendicular to the migration direction. FA’s were binned on distance from leading edge, and FA area and orientation were plotted as functions of distance from leading edge.

### Estimation of VinCS Closed FRET Efficiency and Normalization of VinCS FRET Data

To obtain a reference value for the fully closed FRET Eff of VinCS, VinCS-expressing pMDCK cells were sparsely seeded on poly-L-Lysine coated surfaces, where they non-specifically adhered, as previously done^13^. In detail, glass bottom dishes were coated with Poly-L-Lysine (Sigma P4832-50ML) using the Millipore Sigma Poly-L-Lysine Cell Attachment Protocol. pMDCK cells stably expressing VinCS were sparsely seeded on the pL coated dishes with standard media and allowed to adhere for 30 minutes, after which they were immediately fixed or imaged. FRET Eff was analyzed on a cell basis, using a manual cell mask and minimum acceptor intensity threshold (BSA>1000), identical to the analysis approach for VinCS in the cytosol of MDCK monolayers. Cells that were highly spread, as determined by bright field images, possessed cell-substrate adhesions, as determined by VinCS localization, or had too low VinCS expression (>50% of pixels had BSA values below the BSA threshold) were excluded. The mean FRET Eff from the VinCS on poly-L-Lysine experiments was used to normalize the FRET Eff values for VinCS in cell monolayers.

### Confocal Imaging and FRET Analysis

For confocal imaging, samples were imaged with an Andor XD Revolution Spinning Disk Confocal, which consists of an Olympus IX81 inverted microscope equipped with a Yokogawa CsuX-1 spinning disk (5000rpm) controlled with Metamorph Software. This microscope is maintained by the Duke Light Microscope Core Facility. Images were acquired at 100x magnification (UPlanSApo 100X/NA1.4 Objective, Olympus) using an Andor EMCCD Camera (Ixon3 897 512 EMCCD). The FRET images were acquired using a filter set comprised of a mTFP1 emission filter (483/32), Venus emission filter (542/27) and dichroic mirror (CYR; 445/515/561). For FRET microscopy, three images were acquired. These images included the acceptor (515nm 50mW diode excitation, Venus emission), FRET (445nm 40mW diode excitation, Venus emission), and donor (445nm 40mW diode excitation, mTFP1 emission). Images were acquired without gain and a 75% laser power. Exposure times for Venus, FRET, and Teal were 1000ms, 1500ms, and 1500ms, respectively. Images were post-processed to correct for dark current and aligned using a custom MATLAB script. Ratiometric FRET images were determined by dividing the FRET image by its respective mTFP1 image. Segmentation of cells and adhesions was conducted on the acceptor channel using a custom MATLAB script and user-defined masking.

### Stimulated Emission Depletion Microscopy (STED) Imaging of Actin

Cells were fixed, permeabilized and blocked as described previously. For STED imaging, cells were labeled with Alexa Fluor 488 phalloidin (Invitrogen) at a concentration of 1:25. Following immunofluorescent staining, PBS was removed from the sample, and ProLong™ Diamond Antifade Mountant (Invitrogen, P36965) was applied per manufacturer’s instructions. Mountant set for 24hrs before imaging. Samples were imaged using a Leica STED Confocal, which consists of an inverted Leica DMi8 Platform with motorized scanning stage and controlled by LAS X. Images were acquired at 93x (HC PL APO 93X/1.30 GLYC motCORR, Leica). Alexa Fluor 488 was excited with a tunable white light laser and simultaneously depleted with a 660nm laser. Sample emission was collected using a high-sensitivity, gated GaAsP HyD detector. Hyugens deconvolution, linked to Leica’s LAS X software, was implemented to deconvolve image stacks.

### Statistics

Statistical analyses were performed using JMP Pro (SAS, Cary, NC) software. Comparisons of data with equal variances, as determined with Levene’s test, were analyzed with an ANOVA and, if necessary, Tukey’s Honest Significant Difference (HSD) tests. Datasets with unequal variances were analyzed with a non-parametric Welch’s ANOVA and, if necessary, the Steel-Dwass multiple comparisons test. A *p* value of *p* < 0.05 was considered statistically significant. In figures, a single asterisk (*), double asterisk (**), triple asterisk (***), and quadruple asterisk (****) indicate p-values less than 0.05, 0.01, 0.001, and 0.0001 respectively, and ns indicates a p-value greater than or equal to 0.05. Where used, standard box plots were created using JMP Pro, where the bottom and top of the box indicate the first and third quartiles, respectively, the middle line indicates the median, the whiskers extend to the outermost data points below the first quartile and above the third quartile that are within 1.5 times the interquartile range, and data outside the whiskers are indicated as points.

### Computational Friction Clutch Models of FA and AJ

Details on the computational friction clutch models of the FA and AJ and their implementation are provided in the Supplementary Note 1. MATLAB code used to simulate the model can be made available on request to the corresponding author.

### Code availability

Computer code used in this study can be made available on request to the corresponding author.

### Data availability

All data supporting the findings of the study are available from the authors on reasonable request.

## Acknowledgements and Funding

We thank Dr. Adam Kwiatkowski (University of Pittsburgh) for providing MDCK II cells used in this study and Dr. Akira Nagafuchi (Nara Medical University) for providing the α-catenin conformation-sensitive antibody (α18). This research was supported by the National Institute of Health (1R01GM121739) and the National Science Foundation (GRFP DGE 1644868).

## Author Contributions

B.D.H. conceived the project and obtained funding. T.C.S., E.M.G., J.I.C., and D.E.C. created key reagents and/or cell lines. T.C.S. and E.M.G. designed and conducted experiments. T.C.S. and E.M.G. performed data analyses. T.C.S. designed, implemented, and analyzed mathematical models. T.C.S. and B.D.H. wrote and edited the paper.

## Additional Information

**Extended Data Figures** are included following main figures.

**Supplementary Information** Supplementary Notes I and II are included in separate documents.

## FIGURE LEGENDS

**Extended Data Fig. 1.**
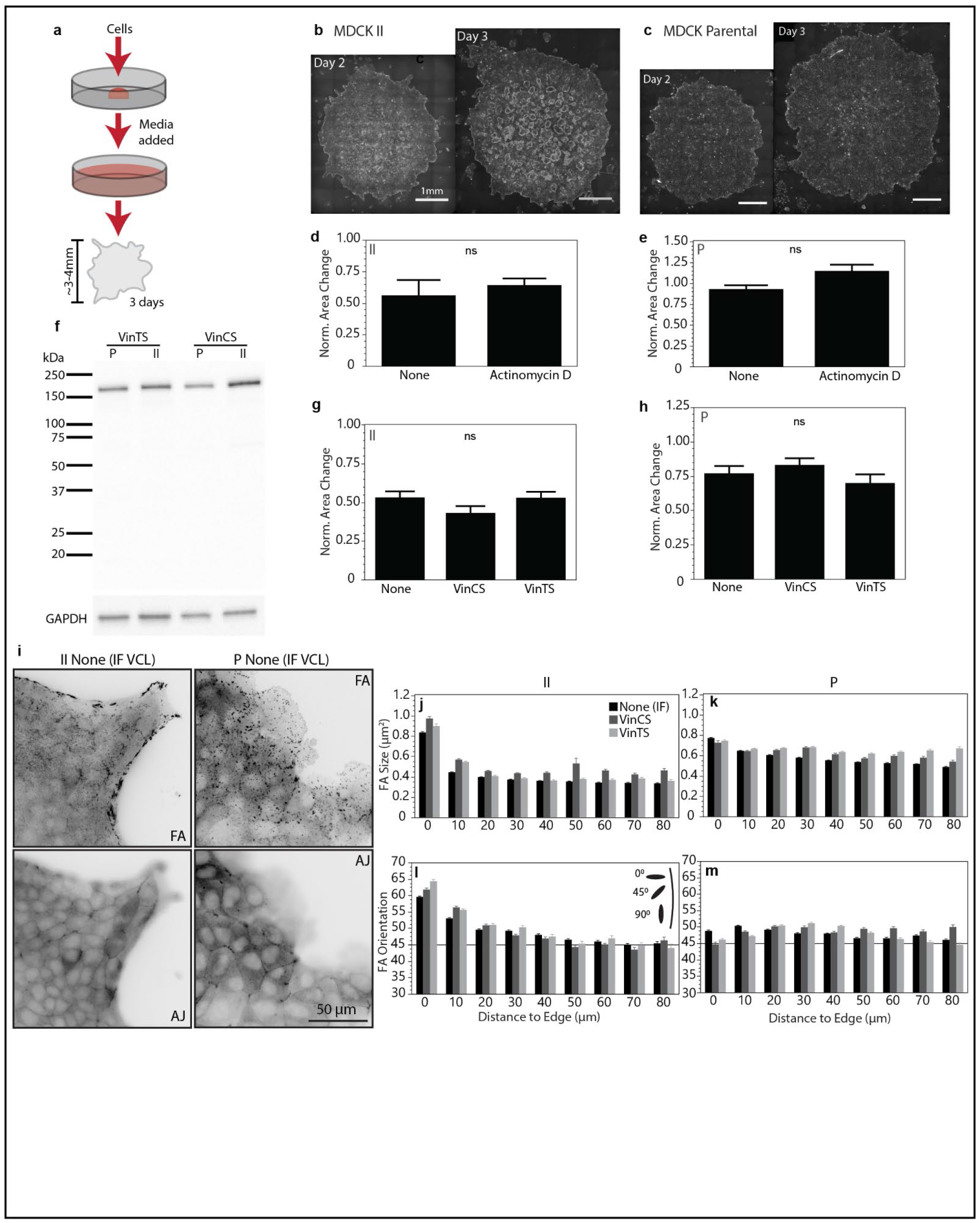
Controls for droplet island assay and expression of VinTS and VinCS in MDCK cells. (a) Schematic depiction of droplet island assay. (b-c) Representative phase contrast images of migrating MDCK II or MDCK Parental islands at 2 and 3 days after droplet seeding. (d) Bar plot (mean +/- SEM) of normalized area change of droplet island assay between days 2 and 3 with or without treatment with Actinomycin D for MDCK II cells (n=3 or 4 islands, respectively, over at least 3 independent experiments). (e) Same for MDCK Parental cells (n=3 or 4 islands, respectively, over at least 3 independent experiments). (f) Western blot with GFP primary antibody showing that VinTS and VinCS are produced as stable proteins with the expected molecular weight in both MDCK II and MDCK Parental cells. (g) Bar plot (mean +/- SEM) of normalized area change of droplet island assay between days 2 and 3 for MDCK II cells expressing no sensor (“None”), VinCS, or VinTS cells (n=12, 4, or 7 islands, respectively, over at least 3 independent experiments). (h) Same for MDCK Parental cells (n=11, 5, or 4 islands, respectively, over at least 3 independent experiments). (i) Representative images of vinculin immunolabeling at the edge of migrating MDCK II or MDCK Parental monolayers in the basal (FAs) or apical (AJs) plane. (j-m) Plots of FA size or FA orientation versus distance from edge for MDCK II or MDCK Parental cells expressing no sensor (“None”), VinCS, or VinTS. Differences between pairs in (d-e) were tested for using t-test and differences between groups in (g-h) were tested for using ANOVA (ns: not significant).

**Extended Data Fig. 2.**
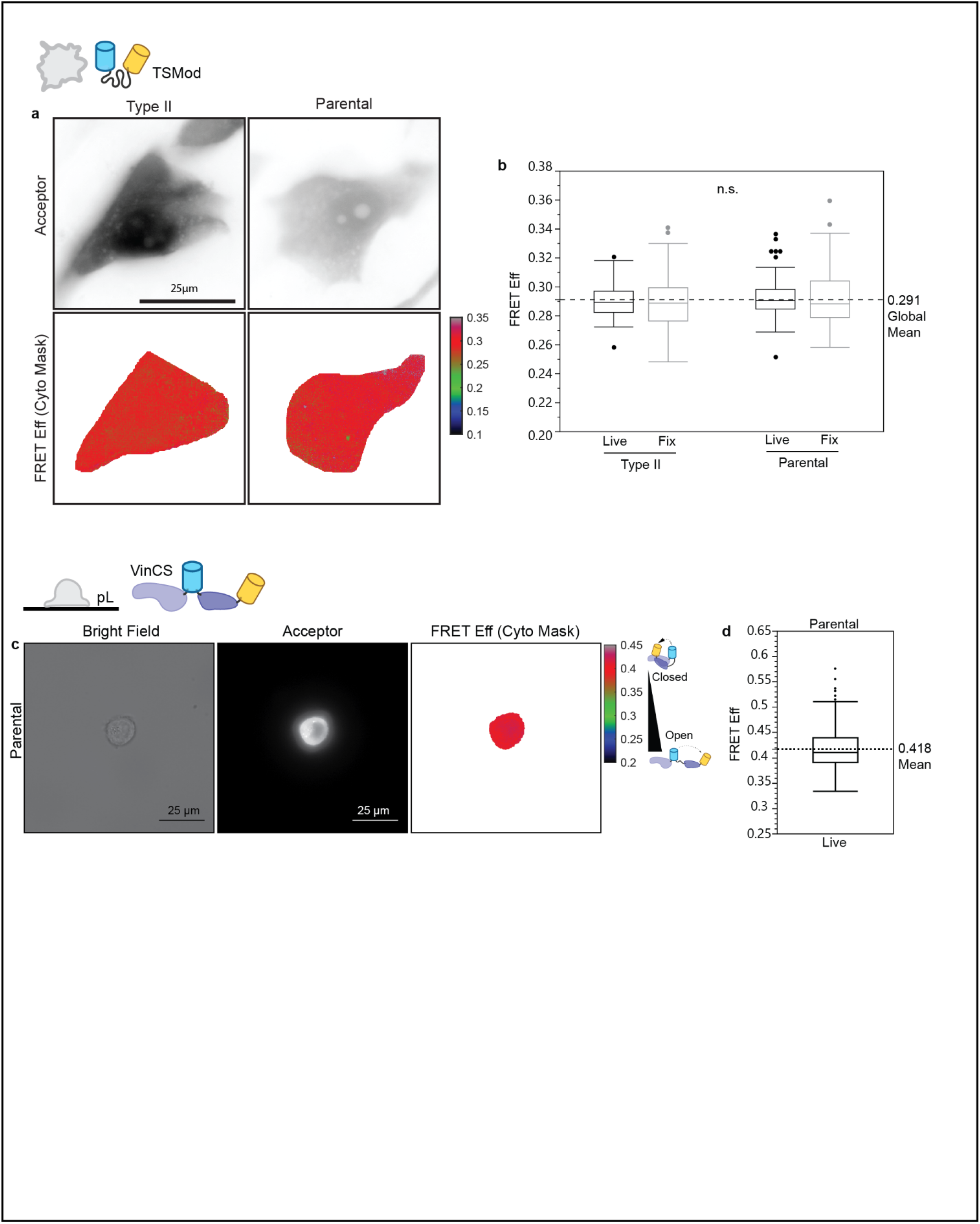
Reference Conditions for VinTS (TSMod) and VinCS (pL). (a) Representative acceptor and masked FRET efficiency images of TSMod in the cytoplasm of cells at the edge of migrating MDCK II or MDCK Parental monolayers. (b) Box-whisker plot showing FRET efficiency for TSMod in MDCK II cells in live or fixed condition or MDCK Parental cells in live or fixed condition (n=117, 102, 119, and 120 cells, respectively, over 4 independent experiments). Differences between groups were tested for using a non-parametric Welch’s ANOVA (ns: not significant). (c) Representative bright field, acceptor, and masked FRET efficiency images of VinCS in the cytoplasm of a MDCK Parental cell adhered to poly-L-lysine surface in the live condition. (d) Box plot shows FRET efficiency for VinCS in the cytoplasm of single MDCK Parental cells adhered to poly-L-lysine surfaces in the live condition, with mean indicated by the dashed line (n=244 cells over 3 independent experiments).

**Extended Data Fig. 3.**
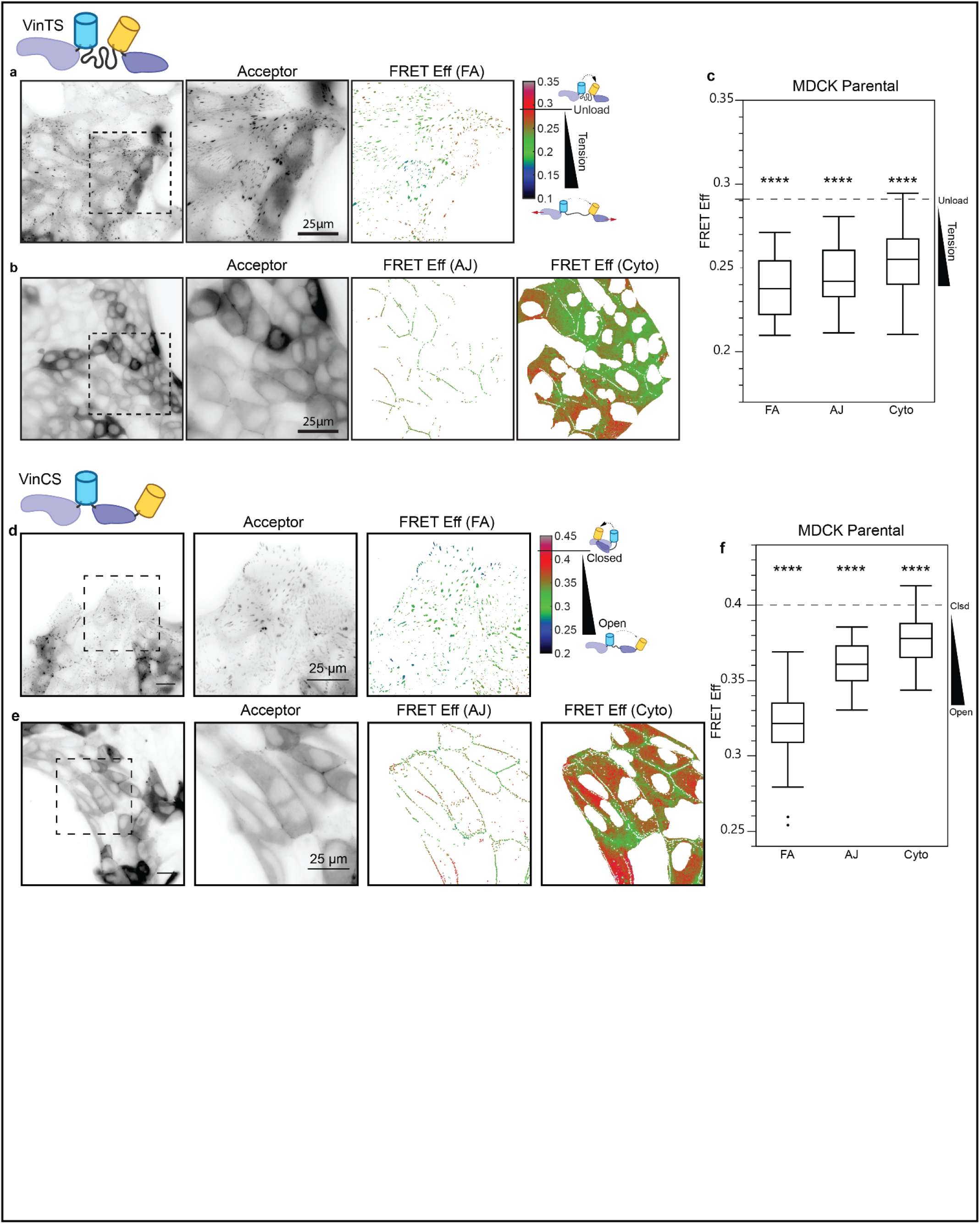
VinTS and VinCS at the edge of collectively migrating MDCK Parental cells. (a) Representative image field of VinTS at the edge of migrating MDCK Parental cell monolayers in the basal plane with acceptor channel indicating sensor localization followed by zoom-in views of acceptor channel and FRET efficiency in the FA mask for the indicated region. (b) Representative image field of VinTS in the apical plane with acceptor channel followed by zoom-in views of acceptor channel and FRET efficiency in AJ and cytoplasm masks for the indicated region. (c) Box-whisker plot showing FRET efficiency for VinTS at FAs, AJs, and cytoplasm (n=67, 48, and 48 image fields respectively over at least 3 independent experiments). Differences between groups were detected using the Steel-Dwass test (****p < 0.0001). P-values shown are for comparisons to VinTS-I997A in MDCK Parental cells at the same structure (Extended Data Fig. 4f); p values for all comparisons can be found in Supplemental Note 2 Table S3. (d) Representative image field of VinCS at the edge of migrating MDCK Parental cell monolayers in the basal plane with acceptor channel indicating sensor localization followed by zoom-in views of acceptor channel and FRET efficiency in the FA mask for the indicated region. (e) Representative image field of VinCS in the apical plane with acceptor channel followed by zoom-in views of acceptor channel and FRET efficiency in AJ and cytoplasm masks for the indicated region. (f) Box-whisker plot showing FRET efficiency for VinCS at FAs, AJs, and cytoplasm (n=103, 51, and 53 image fields respectively over at least 3 independent experiments). Differences between groups were detected using the Steel-Dwass test (****p < 0.0001). P-values shown are for comparisons to the VinCS reference condition (Extended Data Fig. 2c); p values for all comparisons can be found in Supplemental Note 2 Table S2.

**Extended Data Fig. 4.**
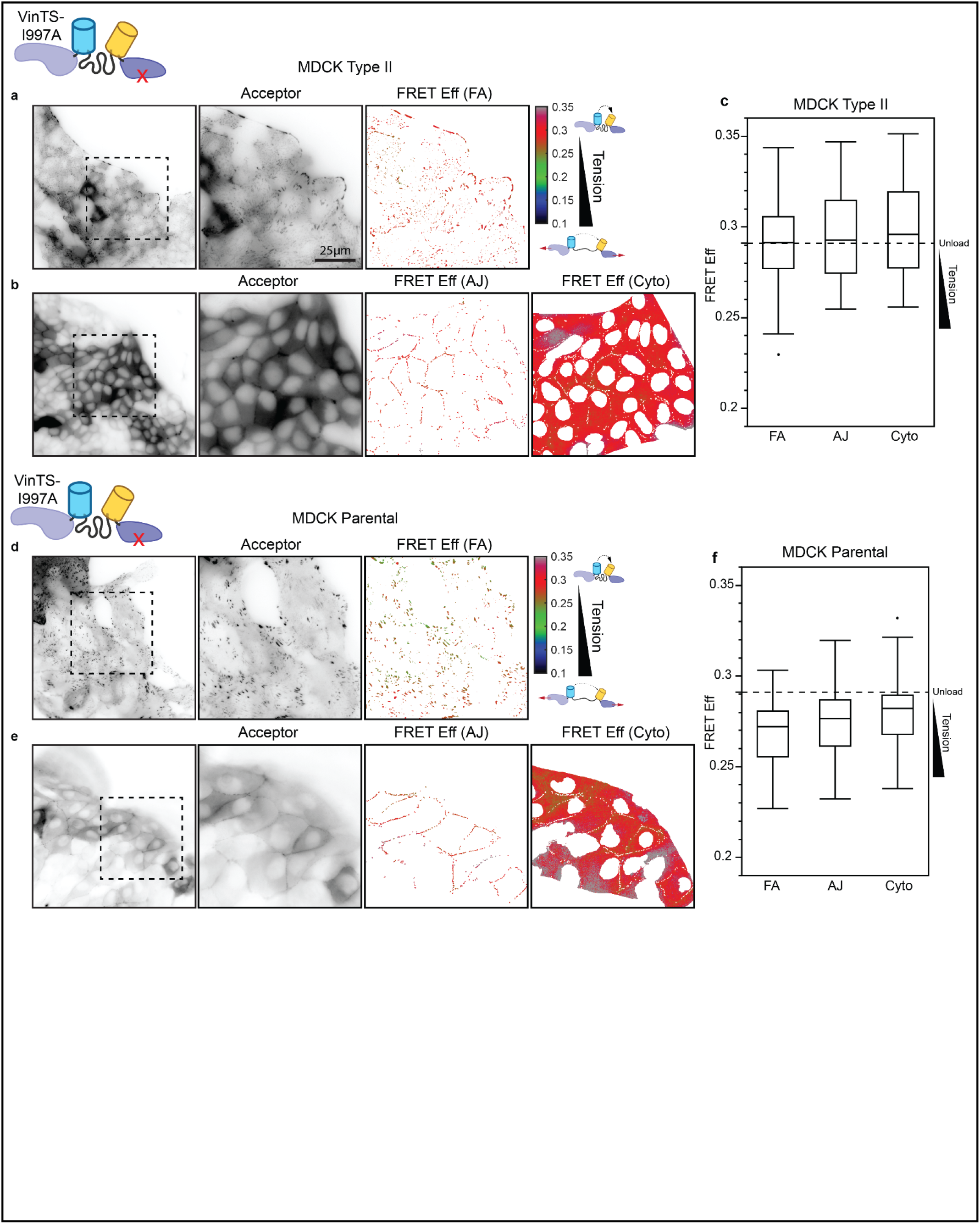
VinTS-I997A at the edge of collectively migrating MDCK II and MDCK Parental cells. (a) Representative image field of VinTS-I997A at the edge of migrating MDCK II cell monolayers in the basal plane with acceptor channel indicating sensor localization followed by zoom-in views of acceptor channel and FRET efficiency in the FA mask for the indicated region. (b) Representative image field of VinTS-I997A in the apical plane with acceptor channel followed by zoom-in views of acceptor channel and FRET efficiency in AJ and cytoplasm masks for the indicated region. (c) Box-whisker plot showing FRET efficiency for VinTS-I997A at FAs, AJs, and cytoplasm (n=52, 31, and 31 image fields respectively over at least 3 independent experiments). (d-f) Analogous representative images and plot for VinTS-I997A in MDCK Parental cells (n=36, 24, and 25 image fields for FAs, AJs, and cytoplasm, respectively, over at least 3 independent experiments).

**Extended Data Fig. 5.**
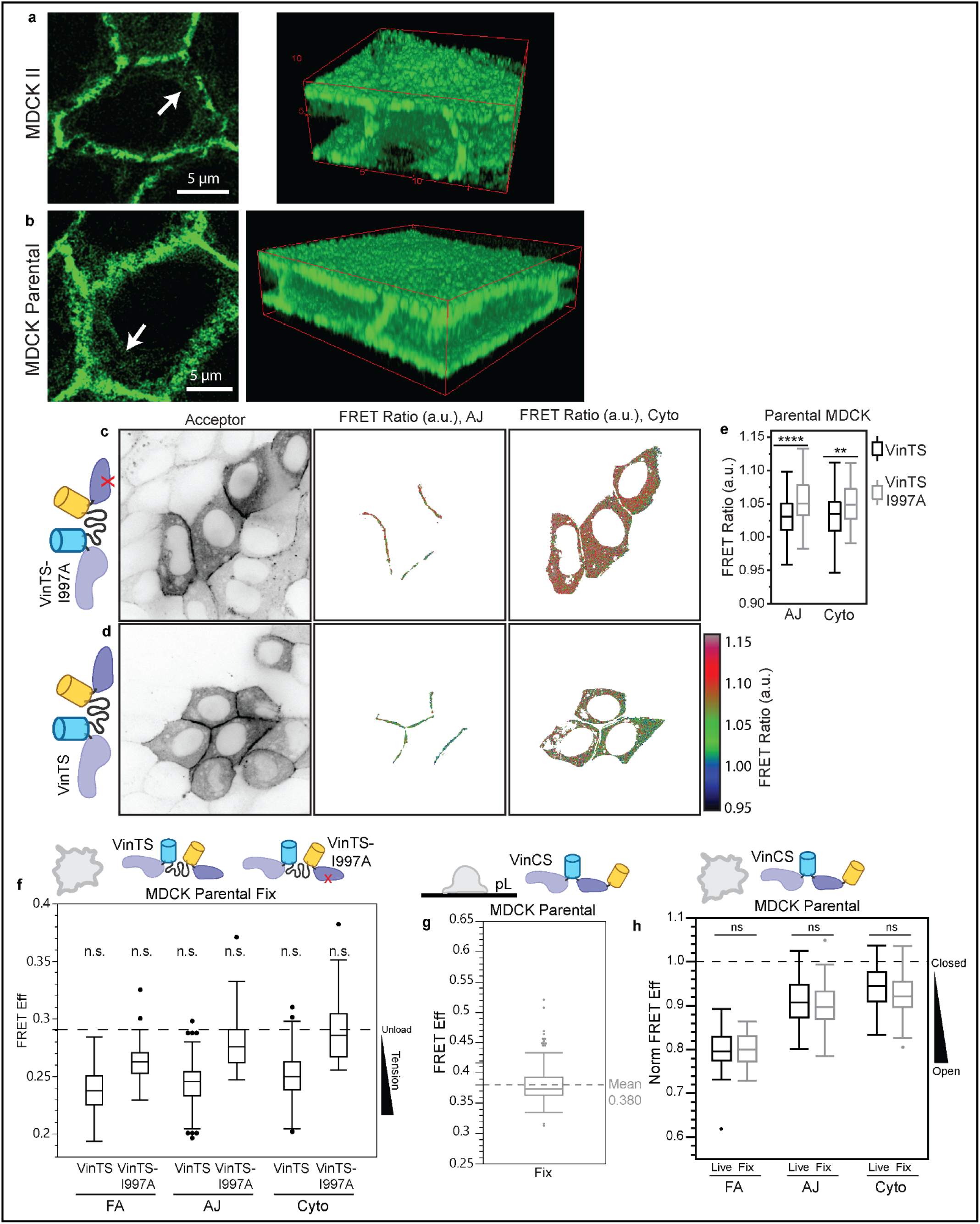
Super-resolution imaging of actin, confocal imaging of VinTS, controls for the fixation of VinTS, and normalization for the fixation of VinCS. (a-b) Stimulated emission depletion (STED) super-resolution imaging of phalloidin-labeled actin in collectively migrating MDCK II and MDCK Parental cells. White arrows indicate regions exhibiting a diffuse, cytoplasmic actin network. (c-d) Representative image fields for confocal imaging of VinTS or VinTS-I997A at the edge of migrating MDCK Parental cell monolayers in the apical plane with acceptor channel indicating sensor localization followed by FRET ratio in AJ and Cytoplasm masks. (e) Box plot showing FRET ratio in AJs and Cytoplasm for confocal imaging of VinTS and VinTS-I997A (n = 207 and 160 junctions for VinTS and VinTS-I997A AJs, respectively, and 63 and 44 cells for VinTS and VinTS-I997A Cytoplasm, respectively, over 3 independent experiments). (f) Box plot showing FRET efficiency at the FAs, AJs, and cytoplasm for VinTS (n=152, 146, and 146 image fields respectively over at least 3 independent experiments) and VinTS-I997A (n=61, 76, and 76 image fields respectively over at least 3 independent experiments) at the edge of migrating MDCK Parental cell monolayers. Differences between groups were detected using the Steel-Dwass test (ns: not significant). P-values shown are for comparisons to the respective construct at the respective structure in MDCK Parental cells in the live condition (Extended Data Fig. 3 and 4); p values for all comparisons can be found in Supplemental Note 2 Table S3. (g) Box plot showing FRET efficiency for VinCS in the cytoplasm of single MDCK Parental cells adhered to poly-L-lysine surfaces in the fixed condition, with mean indicated by the dashed line (n=164 cells over 3 independent experiments). (h) Box plot shows normalized FRET efficiency for VinTS at the edge of MDCK Parental cell monolayers at the FAs, AJs, and cytoplasm in the live condition (n=40, 23, and 23 respectively over 2 independent experiments) or fixed condition (n=67, 58, and 58 respectively over 6 independent experiments, repeated from Fig 2 to show comparison).

**Extended Data Fig. 6.**
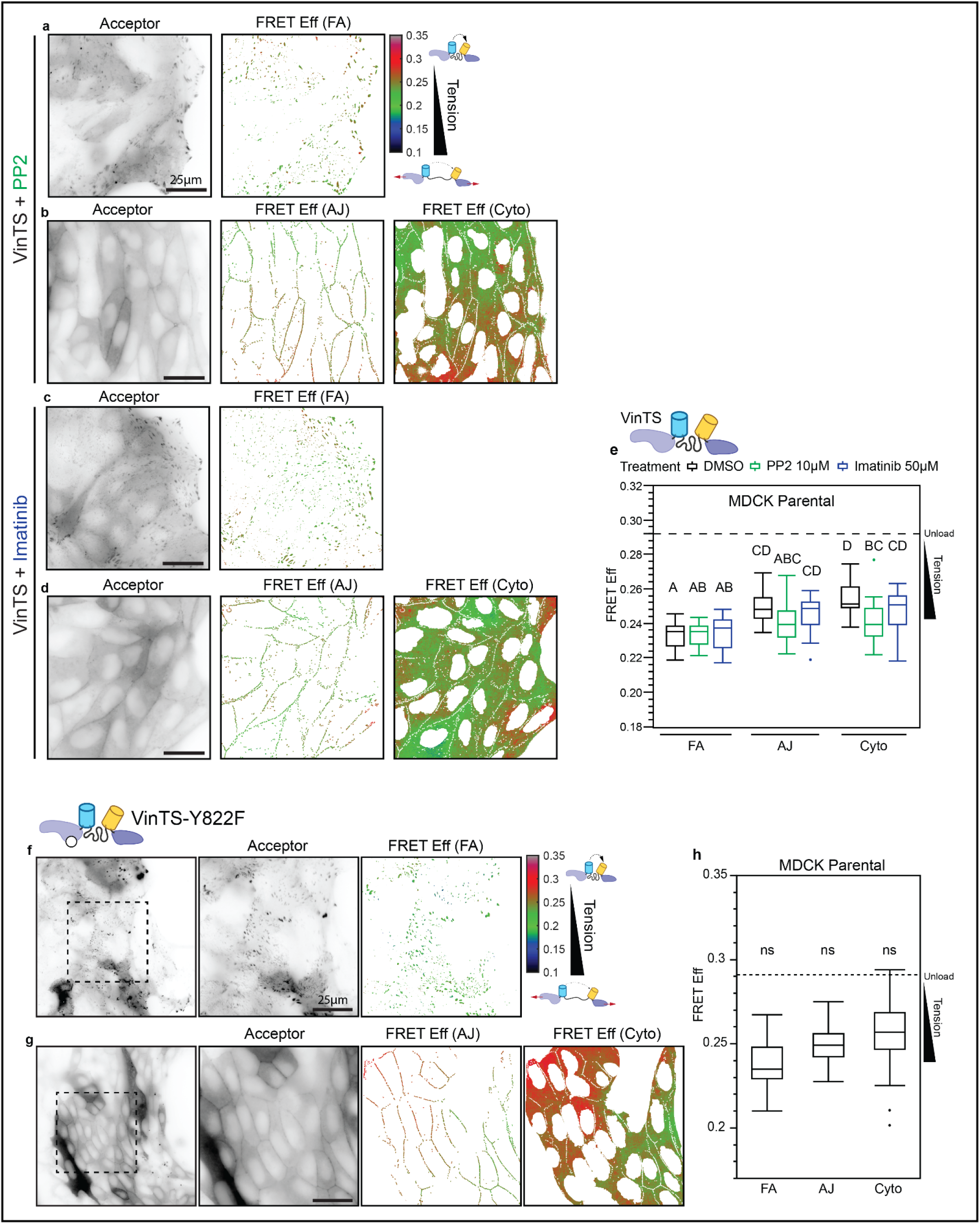
Effect of Src and Abl inhibition and Y822F point mutation on vinculin loading at the edge of collectively migrating cells. (a) Representative image field of VinTS in PP2-treated MDCK Parental cells in the basal plane with acceptor channel and FRET efficiency in the FA mask. (b) Representative image field of VinTS in PP2-treated MDCK Parental cells in the apical plane with acceptor channel and FRET efficiency in AJ and cytoplasm masks. (c-d) Analogous representative image fields for VinTS in Imatinib-treated MDCK Parental cells. (e) Box-whisker plot showing FRET efficiency of VinTS at FAs, AJs, and cytoplasm of untreated (n=25, 23, and 23 image fields respectively over 3 independent experiments), PP2-treated (n=25, 21, and 21 image fields respectively over 3 independent experiments), and Imatinib-treated (n=23, 26, and 26 image fields respectively over 3 independent experiments) cells. Differences between groups were detected using the Tukey HSD test. Levels not connected by the same letter are significantly different. (f) Representative image field of VinTS-Y822F at the edge of migrating MDCK Parental cell monolayers in the basal plane with acceptor channel indicating sensor localization followed by zoom-in views of acceptor channel and FRET efficiency in the FA mask for the indicated region. (g) Representative image field of VinTS-Y822F in the apical plane with acceptor channel followed by zoom-in views of acceptor channel and FRET efficiency in AJ and cytoplasm masks for the indicated region. (h) Box-whisker plot showing FRET efficiency of VinTS-Y822F at FAs, AJs, and cytoplasm (n=44, 42, and 58 image fields respectively over at least 3 independent experiments). Differences between groups were detected using the Steel-Dwass test (ns: not significant). P-values shown are for comparisons to VinTS in MDCK Parental cells at the same structure (Extended Data Fig. 5); p values for all comparisons can be found in Supplemental Note 2 Table S3.

**Extended Data Fig. 7.**
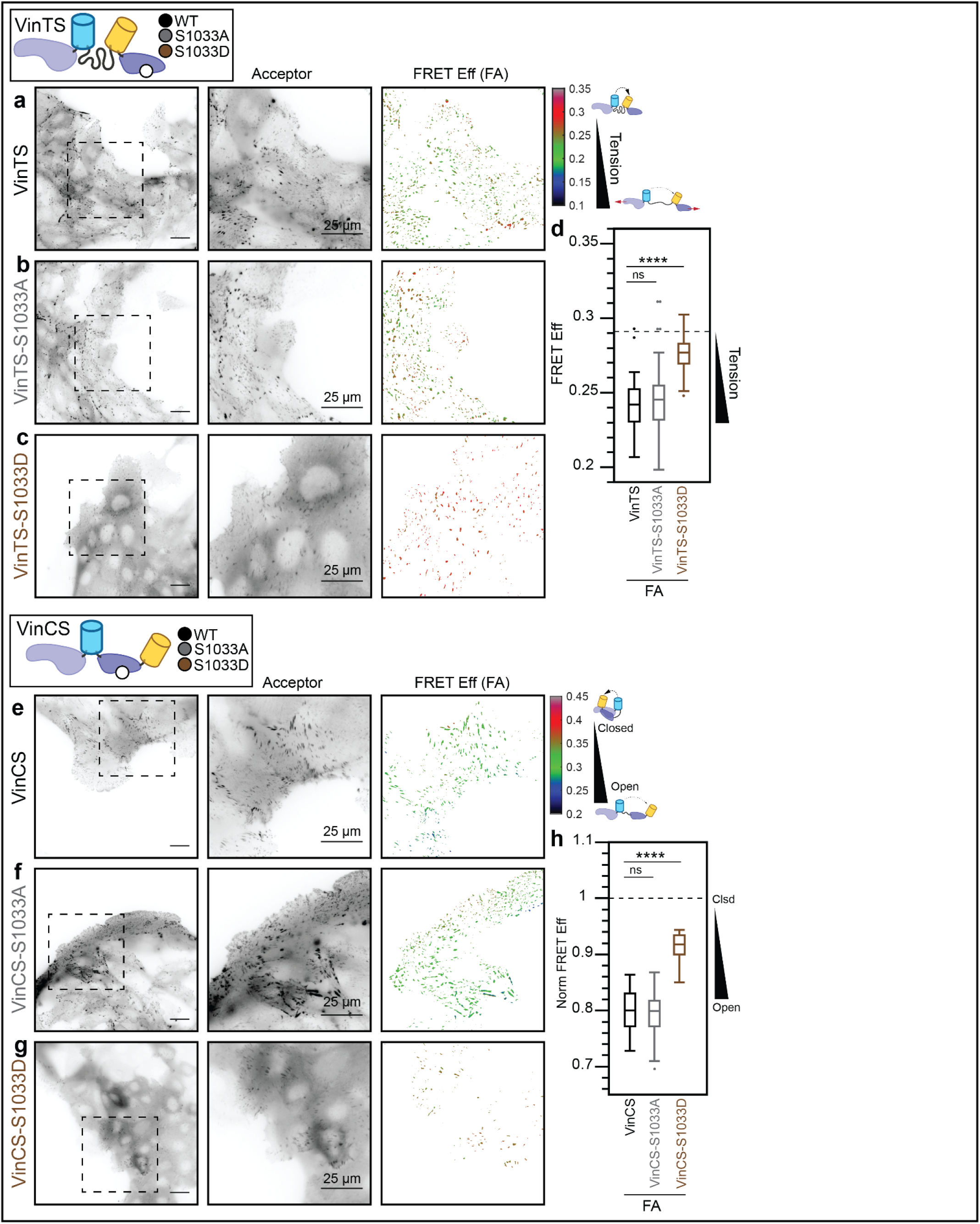
Effect of vinculin S1033 mutants on vinculin load and conformation in FAs at the leading edge of collectively migrating cells. (a-c) Representative image fields of VinTS, VinTS-S1033A, or VinTS-S1033D at the edge of MDCK Parental cell monolayers in the basal plane with acceptor channel indicating sensor localization followed by zoom-in views of acceptor channel and FRET efficiency in FA masks for the indicated region. (d) Box-whisker plot showing FRET efficiency for VinTS, VinTS-S1033A, and VinTS-S1033D in FAs (n=85, 55, and 49 image fields respectively over at least 3 independent experiments). (e-g) Representative image fields of VinCS, VinCS-S1033A, or VinCS-S1033D at the edge of MDCK Parental cell monolayers in the basal plane with acceptor channel indicating sensor localization followed by zoom-in views of acceptor channel and FRET efficiency in FA masks for the indicated region. (h) Box-whisker plot showing FRET efficiency for VinTS, VinTS-S1033A, and VinTS-S1033D in FAs (n=67, 35, and 30 image fields respectively over at least 3 independent experiments). Differences between groups were detected using the Steel-Dwass test (****: p < 0.0001, ns: not significant); p values for all comparisons can be found in Supplemental Note 2, Tables S4-5.

**Extended Data Fig. 8.**
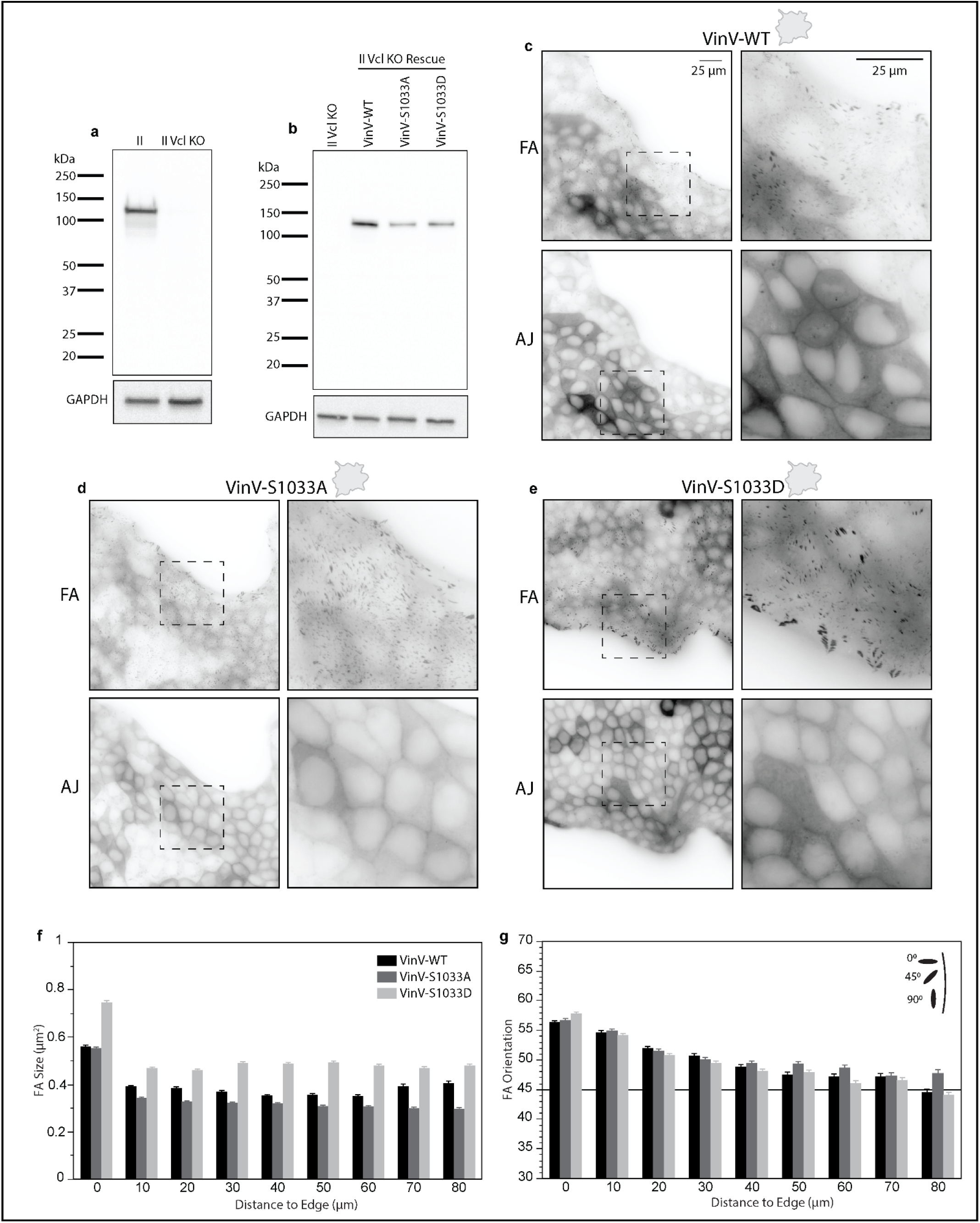
Rescue of Vcl KO MDCK II cells with Vinculin-mVenus and S1033 mutants. (a) Western blot with vinculin antibody confirming CRISPR/Cas9-mediated knockout of vinculin in MDCK II cells. (b) Western blot with GFP primary antibody showing production of stable proteins with the expected molecular weight for rescue of MDCK II Vcl KO cells with Vinculin-mVenus (VinV-WT or VinV), Vinculin-mVenus-S1033A (VinV-S1033A), or Vinculin-mVenus-S1033D (VinV-S1033D). (c-e) Representative image fields of VinV, VinV-S1033A, or VinV-S1033D at the edge of migrating MDCK II monolayers in the basal (FAs) and apical (AJs) plane, with zoom-in views. (f-g) Plots of FA size and FA orientation versus distance from edge for VinV, VinV-S1033A, or VinV-S1033D MDCK II monolayers.

**Extended Data Fig. 9.**
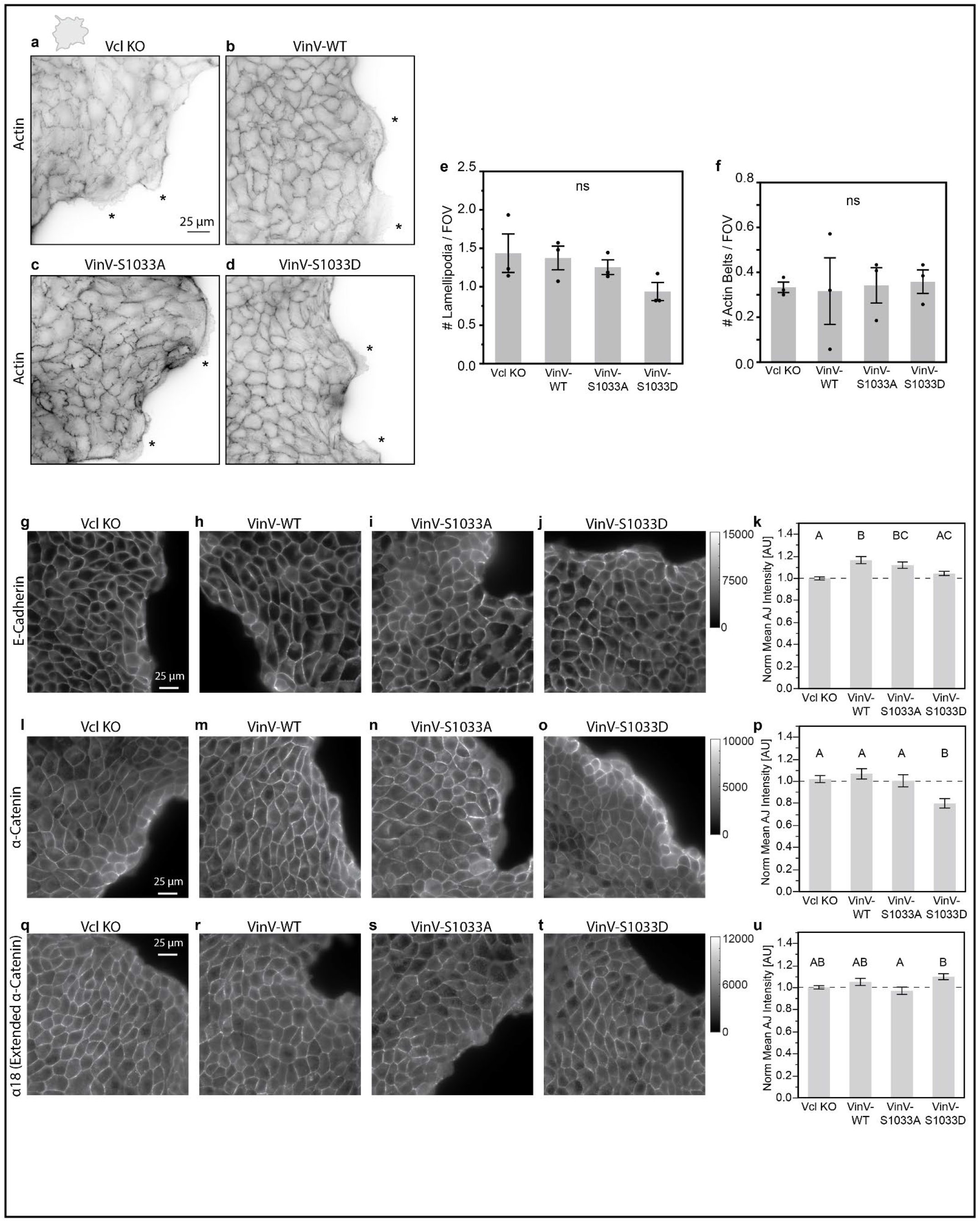
Expression of VinV, VinV-S1033A, or VinV-S1033D does not affect actin structures at the leading edge and has small or no effects on the abundance of E-cadherin or abundance of total or extended α-catenin at AJs. (a-d) Representative image fields of phalloidin labeling at the edge of migrating Vcl KO, VinV, VinV-S1033A, or VinV-S1033D MDCK II cell monolayers. Stars indicate manually identified lamellipodia. (e-f) Bar plots showing number of lamellipodia per image field or number of actin belts per image field for Vcl KO, VinV, VinV-S1033A, or VinV-S1033D MDCK II cells (n=3 droplet island assays per cell line over 3 independent experiments). (g-j) Representative image fields of E-cadherin immunolabeling at the edge of migrating Vcl KO, VinV, VinV-S1033A, or VinV-S1033D MDCK II cell monolayers. (k) Bar plot showing normalized mean E-cadherin stain intensity in AJ masks for Vcl KO, VinV, VinV-S1033A, and VinV-S1033D MDCK II cells (n=48, 44, 41, and 41 images over 3 independent experiments). (i-o) Representative image fields of α-catenin immunolabeling at the edge of migrating Vcl KO, VinV, VinV-S1033A, or VinV-S1033D MDCK II cell monolayers. (p) Bar plot showing normalized mean α-catenin stain intensity in AJ masks for Vcl KO, VinV, VinV-S1033A, and VinV-S1033D MDCK II cells (n=37, 39, 36, and 37 images over 3 independent experiments). (q-t) Representative image fields of α-catenin extended conformation-sensitive antibody (α18) immunolabeling at the edge of migrating Vcl KO, VinV, VinV-S1033A, or VinV-S1033D MDCK II cell monolayers. (u) Bar plot showing normalized mean α-catenin extended conformation-sensitive antibody (α18) stain intensity in AJ masks for Vcl KO, VinV, VinV-S1033A, and VinV-S1033D MDCK II cells (n=36, 37, 35, and 36 images over 3 independent experiments). Bar plots indicate mean +/- SEM. Differences between groups in (e-f) were tested for using ANOVA (ns: not significant). Differences between groups in (k), (p), and (u) were detected using the Steel-Dwass test. Levels not connected by the same letter are significantly different.

**Extended Data Fig. 10.**
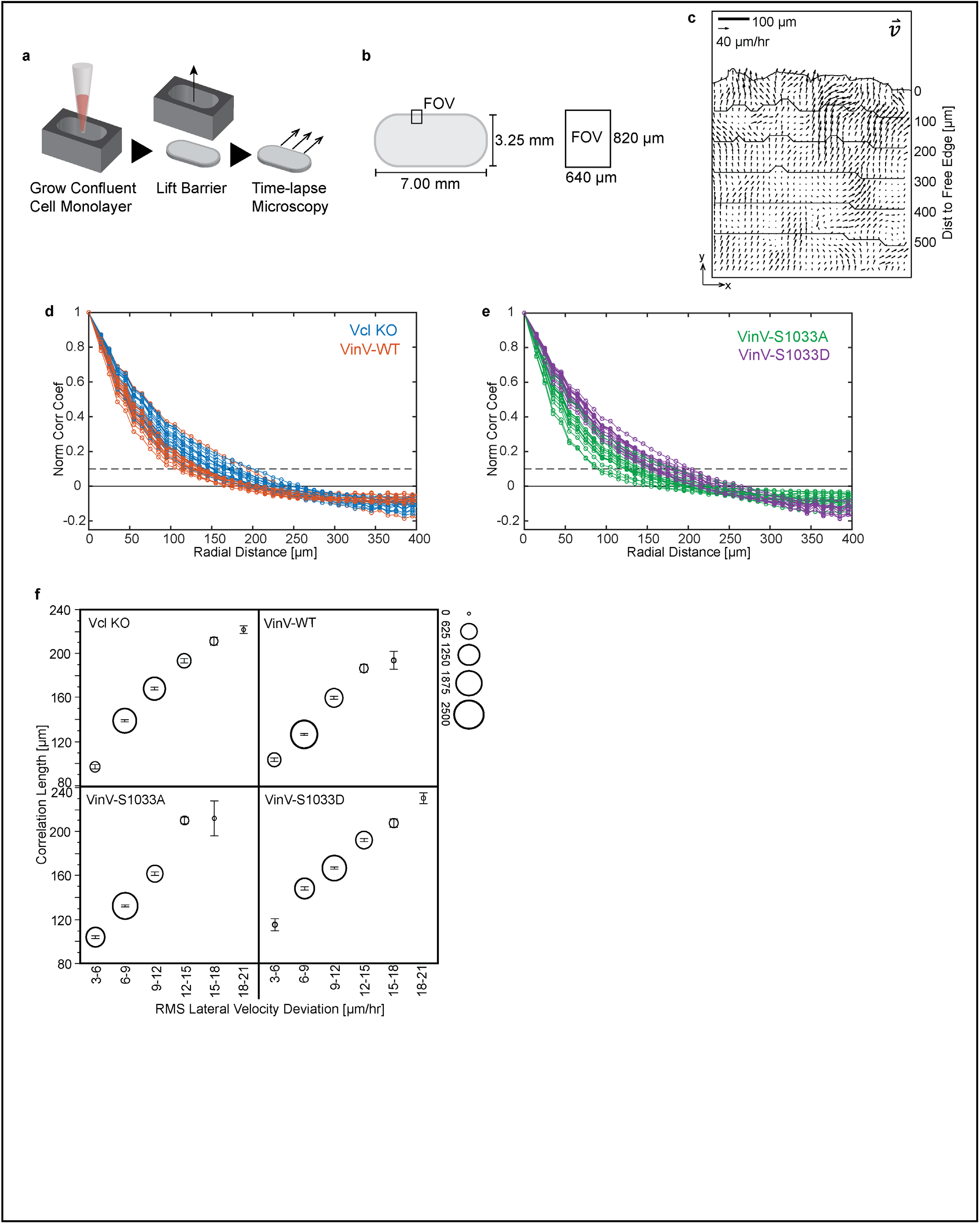
Quantification of migration in barrier assay. (a-b) Schematic of barrier migration assay. (c) Example velocity field. (d-e) Plots of time-averaged normalized spatial correlation coefficient versus radial distance for MDCK II Vcl KO, VinV, VinV-S1033A, or VinV-S1033D cells (n=16 monolayers for each cell line over 6 independent experiments) with dashed line indicating the threshold value for computing the correlation length. (f) Plots of correlation length vs binned RMS lateral velocity deviations for individual image fields and timepoints for MDCK II Vcl KO, VinV, VinV-S1033A, or VinV-S1033D cells combined (>1,500 timepoints for each cell line). Center of circle indicates mean, error bars indicate SEM, and size of circle indicates number of data points in bin. This figure contains additional representations of the same data shown in Fig 3.

