## Supplementary Note 1 for "Coupling During Collective Cell Migration is Controlled by a Vinculin Mechanochemical Switch"

### Supplementary Note 1: Clutch Models of Adhesion-based Friction at the FA and AJ

#### I. OVERVIEW

In this supplemental note, we develop molecularly specific, stochastic models of adhesion-based friction at FAs/AJs based on multi-component integrin-/cadherin-based linkages with force-sensitive bond parameters from single molecule experiments. We use the models to probe the relationship between force-activated binding dynamics and adhesion-based friction existing between adjacent cells and with the ECM. The model formulation and implementation are described in Section II. To validate implementation of the friction clutch model and to gain intuition about molecular determinants of friction, we first used a friction clutch model containing generic single-component linkages (Section III-A). As biological linkages in the FAs and AJs are composed of multiple proteins connected through multiple force-sensitive binding interfaces and subject to potential regulation, we developed multi-component integrin/cadherin linkages to use in the frictional clutch models (Section III-B). To assess regulators of cell-ECM/cell-cell friction, we incorporated the multi-component integrin/cadherin linkages into the friction clutch model and assessed relationships between molecular properties of linkages and friction at the FA/AJ, including the effect of increasing the fraction of linkages with loadable vinculin (Section III-C/D). We then verified the robustness of the vinculin-based reinforcement mechanism across a wide range of model parameters (Section III-E). Key assumptions and limitations of the model are discussed in Section IV. Major conclusions are summarized in Section V. Overall, the models indicated that increasing the fraction of loadable vinculin increased the lifetime of cadherin-/integrin-mediated linkages under load and thereby increased the friction coefficient at FAs and AJs in a tunable manner across a range of speeds corresponding to cell velocities during CCM. Increases in friction coefficient were associated with increases in ensemble vinculin molecular tension, and this effect of vinculin was robust to other model parameters. In the context of macroscopic models of CCM, this is consistent with the experimental effect of vinculin loading on CCM dynamics in our system.

#### II. MODEL CONCEPTUALIZATION AND FORMULATION

Cell migration requires the transmission of forces between cells and their environment. This process occurs at low Re number, meaning cell-generated forces that propel the cell forward are balanced by drag forces that resist motion. Propulsive forces are generated mainly by active processes in the cytoskeleton and are transmitted to the surrounding via specific adhesions. The predominant source of drag forces are the connections between the cell body/cytoskeleton and the external surroundings by specific adhesions. Drag arising from frictional forces at cell-ECM adhesions play a major role in single cell migration[1], and drag forces at both the cell-ECM and cell-cell interfaces play a role in collective cell migration[2,3]. Frictional forces at the cell-ECM and cell-cell interfaces are important components of physical models of CCM[4]. However, the molecular mechanisms that regulate these physical parameters remain poorly understood.

The transmission of forces between the actin cytoskeleton and the substrate or adjacent cell mediated by specialized adhesion structures, termed focal adhesions (FAs) or adherens junctions (AJ) respectively. FAs mechanically couple the actin cytoskeleton to the ECM through integrin transmembrane proteins and various cytosolic adapter proteins. AJs mechanically couple the actin cytoskeletons of adjacent cells through cadherin transmembrane proteins and a set of adapter proteins.

To understand molecular regulators of friction at the FA (cell-ECM) and AJ (cell-cell), and investigate the role of vinculin mechanical loading, we sought to use the frictional clutch framework, which predicts the resistive force due to the sliding of two surfaces relative to each other at a particular speed as function of the number and properties of adhesive linkages between these surfaces[5]. Similar frameworks based on adhesive bond dynamics have also been applied to the cell-ECM and cell-cell drag coefficients in macroscopic models of collective cell migration[2]. In FAs/AJs, integrin-/cadherin-based linkages that contain force-sensitive bonds at multiple interfaces, to F-actin internally and the ECM (Integrin:FN) or adjacent cell (E-Cad:E-Cad) externally[6]. Additionally, both FAs and AJs undergo adhesion reinforcement in response to applied forces, which involves the reinforcement of connections to the actin cytoskeleton via mechanical linker proteins like vinculin. We hypothesized that vinculin, due to its strong catch bond with F-actin[7], could affect the bond lifetime of molecular linkages at FAs and/or AJs and thereby friction. The predominant load-bearing linkage at the FA is Integrin:Talin:F-actin and at the AJ is the minimal cadherin-catenin-F-actin complex (Cadherin:β-Catenin:α-Catenin:F-actin), both of which can be reinforced with an additional mechanical connection to actin via the adapter protein vinculin[6]. Given their importance in force transmission across the FA and AJ, significant work has been conducted to characterize the force-sensitivity of bonds in these multi-protein linkages. In the integrin-based linkage, the force-sensitivities of various integrin heterodimers and ligand pairs, including the Integrin-α5β1:FN[8] and Integrin-αVβ3:FN[9] bonds, as well as the Talin:F-actin[10] and Vinculin:F-actin[7] bonds have been characterized at the single molecule level. In the cadherin-based linkage, the force-sensitive bond kinetics of both the E-cadherin trans-dimer[11] and the α-Catenin:F-actin bond[12] have been elucidated using single molecule techniques.

Using force-sensitive bond kinetics from the single molecule literature, we developed models of mechanical connections between the actin cytoskeleton and external surfaces mediated by focal adhesion and adherens junction that account for the dynamics of molecular linkages at two key interfaces, allowing us to assess what limits force transmission at the molecular scale and the effect of mechanical reinforcement via an adapter protein, here vinculin.

**A. Mechanics of Friction Clutch Models**

As a starting point, we utilized an existing framework for modeling the sliding friction due to adhesive bonds between an actin filament moving with an imposed velocity over a fixed surface[5]. The model considers a 1D interface connected by molecular linkages that behave like Hookean springs and dynamically bind/unbind. The linkages are stretched by the relative motion at the interface and exert a restoring force proportional to their extension. In the FA model, integrin-based molecular linkages connect the actin network inside the cell to ECM ligands on the substrate (Fig S1a). These linkages bind/unbind at two interfaces, the Integrin:FN bond and the Talin:F-actin bond (Section IIC), and can be reinforced with vinculin (Section IIE). Similarly, in the AJ model, E-cadherin-based molecular linkages connect the actin network of one cell to cadherins on the surface of the adjacent cell (Fig S1d). These linkages bind/unbind at two interfaces, the E-cadherin trans-dimer and the α-Catenin:F-actin bond (detailed in Section IID) , and can be reinforced with vinculin (Section IIF). When molecular linkages are bound at both interfaces, they are engaged and transmit forces from the actin cytoskeleton inside the cell to the external substrate (FA) or adjacent cell (AJ). Engaged linkages are represented as set of parallel springs with spring constant $K_{Link}$ that are together in series with an external spring with spring constant $K_{Ext}$, which represents the substrate (FA) or the adjacent cell (AJ). Engaged linkages are extended and loaded by a constant velocity, $V_0$, corresponding to the motion between the actin cytoskeleton (cell body) with respect to the substrate (FA) or the adjacent cell (AJ). At each time step, the total extension of the ith engaged linkage, $x_i$, is updated according to the following equation:

$$x_i(t + \Delta t) = x_i(t) + V_0 \Delta t \tag{1}$$

The total force transmitted across the linkages, $F_{TOT}$, is found by imposing a force balance across the ensemble of $N_{Eng}$ engaged linkage springs that are in series with the external spring.

$$F_{TOT} = \sum_{i=1}^{N_{Eng}} F_{Link,i} = F_{Ext} \quad (2)$$

The external force is related to the extension of the external spring, $x_{Ext}$, by the external spring constant:

$$F_{Ext} = x_{Ext} K_{Ext} \quad (3)$$

The force across an engaged linkage is given by the product of the linkage spring constant and extension of the linkage, $x_{Link,i}$, which is determined by subtracting the extension of the external spring, $x_{Ext}$, from the total extension of the engaged linkage with respect to the resting position of the motor-clutch system, $x_i$ ($x_i = x_{Link,i} + x_{Ext}$).

$$F_{Link,i} = K_{Link} x_{Link,i} = K_{Link}(x_i - x_{Ext}) \quad (4)$$

From these relationships, the total force can be solved for in terms of the spring constants, number of engaged linkages, and the total extension of the engaged linkages.

$$F_{TOT} = \frac{K_{Ext} K_{Link} \sum_{i=1}^{N_{Eng}} x_i}{K_{Ext} + K_{Link} N_{Eng}} \quad (5)$$

Note that this expression for total force is the same relationship used in previous motor-clutch models[13,14], and that the framework used here is equivalent to that of previous motor-clutch models, except that the velocity is constant, corresponding to relative cell motion, instead of force-dependent as for myosin motor-driven actin flow. From the above Equations 2-5, the extension of the external spring, $x_{Ext}$, extension of each linkage, $x_{Link,i}$, and force across each linkage, $F_{Link,i}$, are determined. Parameters for the FA and AJ friction clutch models are given in Table 1.

**Table 1: Parameters for FA and AJ Friction Clutch Models**

| Parameter Name | Parameter Symbol | FA Friction Clutch | AJ Friction Clutch | Rationale |
|---|---|---|---|---|
| Velocity | $V_0$ | Base*: 2.778 nm/s (10 um/hr)<br>Sweep: 0.1-100 nm/s (0.36-360 um/hr) | Base: 2.778 nm/s (10 um/hr)<br>Sweep: 0.1-100 nm/s (0.36-360 um/hr) | FA and AJ**: Base value inside range of MDCK monolayer velocities observed in this work and elsewhere[15], ~1-30 um/hr or ~0.28-8.33 nm/s. Sweep range extends below and above this range. |
| Linkage Spring Constant | $K_{Link}$ | 5 pN/nm | 5 pN/nm | FA and AJ: Within range of effective stiffness for many mechanical proteins[16]. Also, matches value used in previous motor-clutch model of FA[17] and is similar to the experimentally determined value for PCDH15[18]. |
| Number of Linkages | $N_{Link}$ | 50 | 50 | FA and AJ: Matches previous motor-clutch models[13,17]. Correspond to number of integrins or cadherins in adhesive clusters of radius ~100 nm at reported densities of integrin[19] and cadherin[20] molecules in clusters. |
| Binding Rate Constants | FA: | 2/s<br>2/s | 2/s<br>2/s | FA and AJ: Matches previous motor-clutch model with multiple binding interfaces[21]. |

| | | | | |
|---|---|---|---|---|
| | $k_{01}^{Intg:FN}$<br>$k_{01}^{Tal:Factin}$<br>$k_{01}^{Vcl:Factin}$<br>$k_{01}^{Vcl:Tal}$<br>AJ:<br>$k_{01}^{Cad:Cad}$<br>$k_{01}^{aCat:Factin}$<br>$k_{01}^{Vcl:Factin}$<br>$k_{01}^{Vcl:aCat}$ | 2/s<br>1/s | 2/s<br>1/s | Resulting engagement rate for combined linkage is similar to linkage binding rate constant in previous motor-clutch models of FA[13,17]. |
| Unbinding Rate Constants | | Force-Dependent Unbinding Rate Models per Table 2 | Force-Dependent Unbinding Rate Models per Table 3 | Rationale given in Table 2 and Table 3. |
| Fraction of linkages with loadable vinculin | $\rho_{Vcl}$ | Base, Non-reinforced: 0<br>Base, Fully reinforced: 1<br>Sweep: [0, 1] | Base, Non-reinforced: 0<br>Base, Fully reinforced: 1<br>Sweep: [0, 1] | FA and AJ: Varied across full range to assess effect of vinculin mechanical reinforcement. |
| External Spring Constant [pN/nm] | $K_{Ext}$ | $10^6$ pN/nm | 3.16 pN/nm | FA: Set arbitrarily high to match experimental condition in this paper (glass). AJ: Corresponds to estimates of the stiffness of cell monolayers[22,23] (~20-33 kPa). See Section II-J for conversion from elastic modulus to spring constant. |

*For parameters that were swept, the base value and sweep value range are given.
**Labels indicate whether rationales apply to both FA and AJ ("FA and AJ"), only FA ("FA"), or only AJ ("AJ").
***Sensitivity analyses were conducted on all major parameters over +/- 1 decade (see Section III-E).

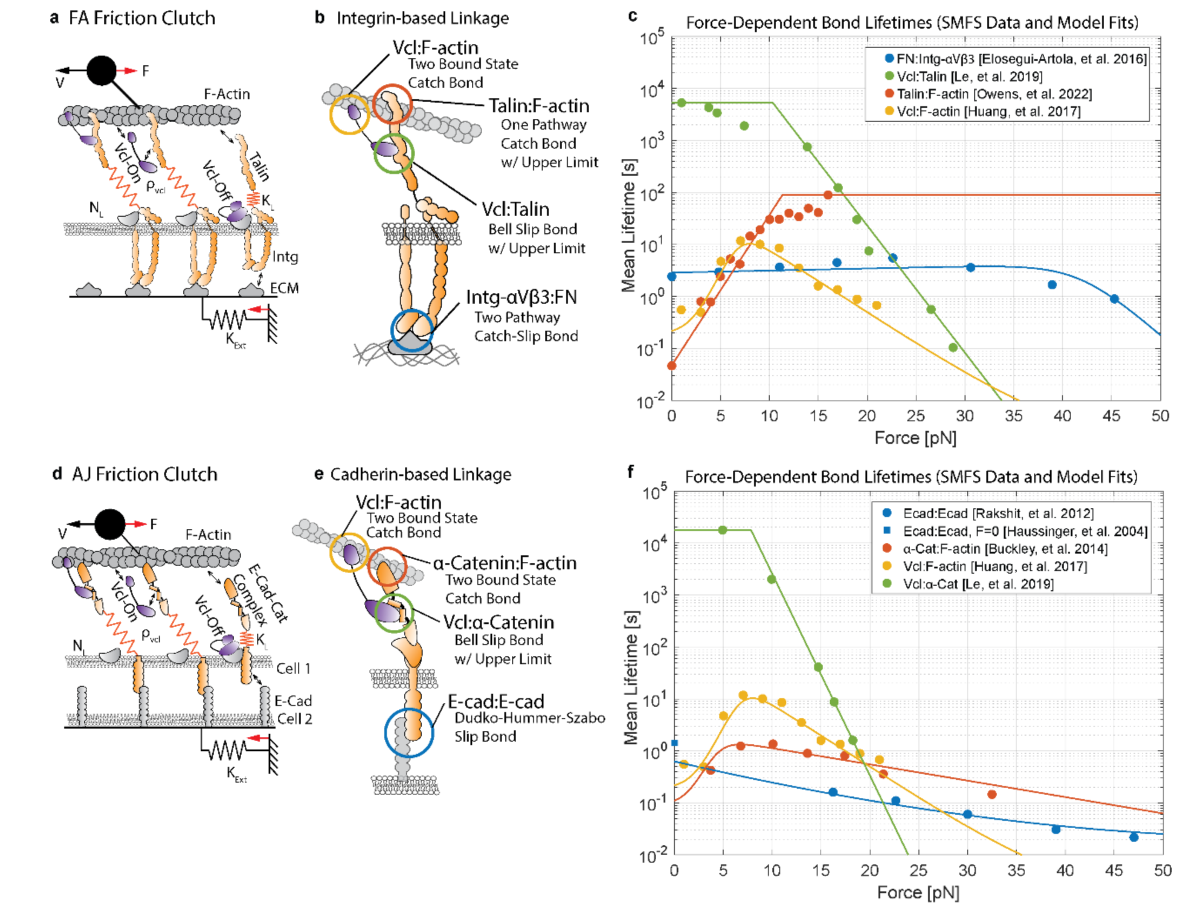


**Fig. S1 Model Formulation for FA and AJ Friction Clutch**. (a) Schematic of FA friction clutch model. (b) Schematic of multi-component integrin-based linkage in FA model indicating binding/unbinding interfaces with the type of force-dependent bond model used for the unbinding rate constants. (c) Plot of mean bond lifetime versus force for each bond in the integrin-based linkage. Data points are reproduced from the indicated experimental studies[7,9,10,24] and lines represent the respective model fit (see Table 2). (d) Schematic of AJ friction clutch model. (e) Schematic of multi-component cadherin-based linkage in AJ model indicating binding/unbinding interfaces with the type of force-dependent bond model used for the unbinding rate constants. (f) Plot of mean bond lifetime versus force for each bond in the cadherin-based linkage. Data points are reproduced from the indicated experimental studies[7,11,12,24,25] and lines represent the respective model fit (see Table 3).

**B. Binding/Unbinding Dynamics in Integrin-based Linkage without Vinculin Reinforcement**

In the FA model, integrin-based molecular linkages dynamically bind/unbind at two interfaces, the Integrin:FN bond (referred to as the integrin interface) and the Talin:F-actin bond (referred to as the F-actin interface (Fig S1c). We chose to model binding/unbinding at these two interfaces, but not between talin and integrin, because there are limited measurements of the force-sensitivity of Talin:Integrin bonds and the fact that Talin is known to be proteolyzed by calpain to facilitate FA release suggests that the Talin:Integrin bond is strong[26]. Also, the fact that vinculin increases the tension across Talin suggests that the Talin:F-actin bond is weaker than the Talin:Integrin bond[27].

The Intg:FN bond is modeled as a single state, two-pathway catch-slip bond, as in previous FA motor-clutch models, and is based on experimental measurements of the bond between Integrin-αVβ3 and FN[9]. Integrin-αVβ3 was used because Integrin-αV is required for mechanotransduction in collectively migrating MDCK cells[28]. However, we note that the FA model containing Integrin-based linkages with Intg:FN kinetics for Integrin-α5β1[8] behaved similarly (data not shown). The state variable for the Integrin:FN bond in the jth linkage is given by:

$$\theta_j^{Intg:FN} = \begin{cases} 0 & Unbound\ State \\ 1 & Bound\ State \end{cases} \tag{6}$$

In the Integrin:FN bond, binding (0→1) occurs with the with the force-independent rate constant $k_{01}^{Intg:FN}$ and unbinding (1→0) occurs with the force-dependent rate constant function $k_{10}^{Intg:FN}(F)$ defined in Table 2, where $F$ denotes the force across the bond (see Fig 1Se for plot of mean lifetime versus force).

The Talin:F-actin bond is modeled as a as a one state catch bond based on single molecule experiments with Talin ABS3 and F-actin[10], with a maximum lifetime. The state variable for the Talin:F-actin bond in the jth linkage is given by:

$$\theta_j^{Tal:Factin} = \begin{cases} 0 & Unbound\ State \\ 1 & Bound\ State \end{cases} \tag{7}$$

In the Talin:F-actin bond, binding (0→1) occurs with the with the force-independent rate constant $k_{01}^{Tal:Factin}$ and unbinding (1→0) occurs with the force-dependent rate constant function $k_{10}^{Tal:Factin}(F)$ defined in Table 2, where $F$ denotes the force across the bond.

To be engaged and transmit forces from the actin cytoskeleton to FN ligands on the substrate, a molecular linkage must be bound at both the integrin and the actin interfaces. This is indicated by the engagement state variable for the jth linkage, $\Theta_j$, which equals 1 when engaged and 0 when disengaged:

$$\Theta_j = \begin{cases} 1 & \theta_j^{Intg:FN} > 0 \land \theta_j^{Tal:Factin} > 0 \\ 0 & elsewhere \end{cases} \tag{8}$$

Disengaged linkages ($\Theta_j = 0$) experience do not transmit force. When linkages transition from disengaged to engaged ($\Theta_j = 0 \rightarrow 1$), the total linkage extension is set to $x_i = x_{Ext}$, as $x_{Link,i} = 0$ at the onset of engagement. Once engaged, the linkage is loaded and extended by the actomyosin network as one of the $N_{Eng}$ engaged linkages described previously. The force across a linkage is thus given by:

$$F_{Link,j} = \begin{cases} K_{Link} x_{Link,j} & \Theta_j = 1 \\ 0 & \Theta_j = 0 \end{cases} \tag{9}$$

where $x_{Link,j}$ , which is defined for engaged linkages, is equal to the total extension of the linkge with respect to the resting position of the motor-clutch system minus the extension component of the external spring: $x_{Link,i} = x_i - x_{Ext} = x_i - \frac{F_{TOT}}{K_{Ext}}$. Because the actin and integrin interfaces are in series, the Integrin:FN and Talin:F-actin bonds both experience the entire linkage force in the minimal integrin-based linkage:

$$F_{Link,j}^{Intg:FN} = F_{Link,j}^{Tal:Factin} = F_{Link,j} \tag{10}$$

These bond forces affect unbinding kinetics according to the force-dependent unbinding rate constants described above with parameters in Table 2. Lastly, when the bond at either interface breaks, the linkage returns to a disengaged state, in which it experiences no extension and bears no force. We note that only the linkage extensions and forces are tracked. Neither the location of linkages nor the sites on F-actin or the substrate (or adjacent cell surface for AJ) to which they bind are explicitly modeled. However, a physical interpretation of disengaged linkages is as follows. Linkages that unbind from the actin interface remain at the position of the ECM on the substrate (or the cadherin on the adjacent cell for AJ) to which they are bound and may rebind F-actin. Linkages that unbind from the integrin interface (or cadherin interface for AJ) remain at the position of their bond with F-actin and are free to bind other FN molecules on the substrate (or cadherin molecules on the adjacent cell for AJ).

**Table 2: Force-Dependent Unbinding Kinetics for Integrin-based Linkage**

| Bond | Model Type | Model Equation(s) | Parameter Values | Literature Reference |
|---|---|---|---|---|
| Talin: F-actin | Catch Bond with Upper Lifetime Limit<br>0 = Unbound<br>1 = Bound | $k_{10}(F) = \max\left\{k_0 \exp(Fx/k_BT), \frac{1}{\tau_{max}}\right\}$ | $k_0$ 21.0084 1/s<br>$x$ -2.7412 nm<br>$\tau_{max}$ 89.3450 s | SMFS data and model fit from Owen et al.[10], for Talin-ABS3 in the negative pull direction. Model fit is single exponential from Owen et al. with max lifetime corresponding to lifetime at max force probed. |
| Vinculin: F-actin | Two Bound State Catch Bond<br>0 = Unbound<br>1= Weak Bound<br>2 = Strong Bound | $k_{10}(F) = k_{10}^0 \exp(Fx_{10}/k_BT)$<br>$k_{20}(F) = k_{20}^0 \exp(Fx_{20}/k_BT)$<br>$k_{12}(F) = k_{12}^0 \exp(Fx_{12}/k_BT)$<br>$k_{21}(F) = k_{21}^0 \exp(Fx_{21}/k_BT)$ | $k_{10}^0$ 5.3 1/s<br>$x_{10}$ 0 nm<br>$k_{20}^0$ 5.5E-3 1/s<br>$x_{20}$ 1.2 nm<br>$k_{12}^0$ 6.1 1/s<br>$x_{12}$ 0.4 nm<br>$k_{21}^0$ 43 1/s<br>$x_{21}$ -3.4 nm | SMFS data and model fit from Huang et al.[7], for Vcl-T12 in the negative pull direction. |
| Vinculin: Talin | Bell Slip Bond with Upper Lifetime Limit<br>0 = Unbound<br>1 = Bound | $k_{10}(F) = \max\left\{k_0 \exp(Fx/k_BT), \frac{1}{\tau_{max}}\right\}$ | $k_0$ 5.57E-7 1/s<br>$x$ 2.3164 nm<br>$\tau_{max}$ 5311.09 s | SMFS data from Le et al.[24]. Data fit to Bell Model with max lifetime corresponding to lifetime at lowest force probed. |
| Intg:FN | Two-Pathway Catch-Slip Bond<br>0 = Unbound<br>1 = Bound | $k_{10}(F) = k_1^0 \exp(Fx_1/k_BT) + k_2^0 \exp(Fx_2/k_BT)$ | $k_1^0$ 0.3471 1/s<br>$x_1$ -0.03741 nm<br>$k_2^0$ 2.87E-8 1/s<br>$x_2$ 1.5669 nm | SMFS data for Integrin-αVβ3:FN bond from Elosegui-Artola et al.[9] (Fig 3A) fit to two-pathway catch-slip bond model[29]. |

**C. Binding/Unbinding Dynamics in Cadherin-based Linkage without Vinculin Reinforcement**

In the AJ model, molecular linkages dynamically bind/unbind at two interfaces, the E-cadherin trans-dimer (referred to as the cadherin interface) and the α-Catenin:F-actin bond (referred to as the F-actin interface) (Fig S1d). As such, the model framework is similar to that of the integrin-based linkage, but with different kinetic parameters. We chose to model binding/unbinding at these two interfaces, but not within the E-cadherin-catenin complex, because the affinities of interactions within the E-cadherin-catenin complex[30] ($K_d$ ~ 1 nM) are substantially higher than those of the E-cadherin-catenin complex for F-actin[31] ($K_d$~1 μM) and the E-cadherin trans-dimer[32] ($K_d$~100 μM). Additionally, the α-Catenin:β-Catenin complex has a higher mechanical stability than that of the E-cadherin trans-dimer or α-Catenin:F-actin bond, with a lifetime of tens to hundreds of seconds up to forces of 10 pN[33].

The E-cadherin:E-cadherin bond is modeled as a single state slip bond based on single molecule experiments with WT and K14E E-cadherin[11]. The state variable for the E-cadherin:E-cadherin bond in the jth linkage is given by:

$$\theta_j^{Cad:Cad} = \begin{cases} 0 & Unbound\ State \\ 1 & Bound\ State \end{cases} \tag{11}$$

In the E-cadherin:E-cadherin bond, binding (0→1) occurs with the with the force-independent rate constant $k_{01}^{Cad:Cad}$ and unbinding (1→0) occurs with the force-dependent rate constant function $k_{10}^{Cad:Cad}(F)$ defined in Table 3, where $F$ denotes the force across the bond (see Fig 1Sf for plot of mean lifetime versus force).

The α-Catenin:F-actin bond is modeled as a two-bound state catch-slip bond based on single molecule experiments with F-actin and the E-cadherin-catenin complex[12]. The state variable for the α-Catenin:F-actin bond in the jth linkage is given by:

$$\theta_j^{aCat:Factin} = \begin{cases} 0 & Unbound\ State \\ 1 & Weak\ Bound\ State \\ 2 & Strong\ Bound\ State \end{cases} \tag{12}$$

In the α-Catenin:F-actin bond, binding only occurs from the unbound to weak bound state (0→1) with the rate constant $k_{01}^{aCat:Factin}$, as done in a previous model of actin dynamics and substrate adhesion using two-state catch-slip bond[7]. Rate constants for unbinding from the weak state, $k_{10}^{aCat:Factin}(F)$, and strong state, $k_{20}^{aCat:Factin}(F)$, and interconversion between the weak and strong states, $k_{12}^{aCat:Factin}(F)$ and $k_{21}^{aCat:Factin}(F)$, are defined in Table 2.

To be engaged and transmit forces from the actin cytoskeleton to cadherins on the surface of the adjacent cell, a molecular linkage must be bound at both the cadherin and the actin interfaces. This is indicated by the engagement state variable for the jth linkage, $\Theta_j$, which equals 1 when engaged and 0 when disengaged:

$$\Theta_j = \begin{cases} 1 & \theta_j^{Cad:Cad} > 0 \wedge \theta_j^{aCat:Factin} > 0 \\ 0 & elsewhere \end{cases} \tag{13}$$

Disengaged linkages ($\Theta_j = 0$) experience do not transmit force. When linkages transition from disengaged to engaged ($\Theta_j = 0 \to 1$), the total linkage extension is set to $x_i = x_{Ext}$, as $x_{Link,i} = 0$ at the onset of engagement. Once engaged, the linkage is loaded and extended by the actomyosin network as one of the $N_{Eng}$ engaged linkages described previously. The force across a linkage is thus given by:

$$F_{Link,j} = \begin{cases} K_{Link} x_{Link,j} & \Theta_j = 1 \\ 0 & \Theta_j = 0 \end{cases} \tag{14}$$

where $x_{Link,j}$ , which is defined for engaged linkages, is equal to the total extension of the linkge with respect to the resting position of the motor-clutch system minus the extension component of the external spring: $x_{Link,i} = x_i - x_{Ext} = x_i - \frac{F_{TOT}}{K_{Ext}}$. Because the actin and cadherin interfaces are in series, the E-cadherin:E-cadherin and α-Catenin:F-actin bonds both experience the entire linkage force in the minimal cadherin-catenin linkage:

$$F_{Link,j}^{Cad:Cad} = F_{Link,j}^{aCat:Factin} = F_{Link,j} \tag{15}$$

These bond forces affect unbinding kinetics according to the force-dependent unbinding rate constants described above. As with the integrin-based linkage, when the bond at either interface breaks, the linkage returns to a disengaged state, in which it experiences no extension and bears no force.

**Table 3: Force-Dependent Unbinding Kinetics for Cadherin-based Linkage**

| Bond | Model Type | Model Equation(s) | Parameter Values | Literature Reference |
|---|---|---|---|---|
| α-Catenin: F-actin | Two Bound State Catch Bond<br>0 = Unbound<br>1= Weak Bound<br>2 = Strong Bound | $k_{10}(F) = k_{10}^0 \exp(Fx_{10}/k_BT)$<br>$k_{20}(F) = k_{20}^0 \exp(Fx_{20}/k_BT)$<br>$k_{12}(F) = k_{12}^0 \exp(Fx_{12}/k_BT)$<br>$k_{21}(F) = k_{21}^0 \exp(Fx_{21}/k_BT)$ | $k_{10}^0$ 11 1/s<br>$x_{10}$ 0 nm<br>$k_{20}^0$ 0.14 1/s<br>$x_{20}$ 0.4 nm<br>$k_{12}^0$ 3 1/s<br>$x_{12}$ 0.2 nm<br>$k_{21}^0$ 20 1/s<br>$x_{21}$ -4 nm | SMFS data and model fit from Buckley et al.[12]. |
| Vinculin: F-actin | Two Bound State Catch Bond<br>0 = Unbound<br>1= Weak Bound<br>2 = Strong Bound | $k_{10}(F) = k_{10}^0 \exp(Fx_{10}/k_BT)$<br>$k_{20}(F) = k_{20}^0 \exp(Fx_{20}/k_BT)$<br>$k_{12}(F) = k_{12}^0 \exp(Fx_{12}/k_BT)$<br>$k_{21}(F) = k_{21}^0 \exp(Fx_{21}/k_BT)$ | $k_{10}^0$ 5.3 1/s<br>$x_{10}$ 0 nm<br>$k_{20}^0$ 5.5E-3 1/s<br>$x_{20}$ 1.2 nm<br>$k_{12}^0$ 6.1 1/s<br>$x_{12}$ 0.4 nm<br>$k_{21}^0$ 43 1/s<br>$x_{21}$ -3.4 nm | SMFS data and model fit from Huang et al.[7], for Vcl-T12 in the negative pull direction. |
| Vinculin: α-Catenin | Bell Slip Bond with Upper Lifetime Limit<br>0 = Unbound<br>1 = Bound | $k_{10}(F) = \max\left\{k_0 \exp(Fx/k_BT), \frac{1}{\tau_{max}}\right\}$ | $k_0$ 5.136E-8 1/s<br>$x$ 3.6771 nm<br>$\tau_{max}$ 1.7691E4 s | SMFS data from Le et al.[24]. Data fit to Bell Model with max lifetime corresponding to lifetime at lowest force probed. |
| E-cad trans-dimer | Slip bond for cusp free energy surface<br>0 = Unbound<br>1 = Bound | $k_{10}(F) = k_0\left(1 - \frac{vFx^\#}{\Delta G^\#}\right)^{1/v-1} \exp\left(\frac{\Delta G^\#}{k_BT}\left[1 - \left(1 - \frac{vFx^\#}{\Delta G^\#}\right)^{1/v}\right]\right)$ | $v$ 0.5<br>$k_0$ 1/6.2 1/s<br>$x^\#$ 0.30 nm<br>$\Delta G^\#$ 25.1 $k_BT$ | SMFS data and global model fit for K14E and WT E-cadherin from Rakshit et al.[11]. Model originates from Dudko et al.[34]. Fit for K14E and WT E-cadherin used because strand swap-dimer is dominant form at equilibrium and the transition from X-dimer to SS-dimer occurs very rapidly. Note: Model fit at F=0 also agrees well with off-rate measurements in solution[25]. |

#### D. Vinculin Reinforcement of Integrin- and Cadherin-based Linkages

Vinculin is a mechanical linker protein that localizes to both FAs and AJs, where it mediates connections to the actin cytoskeleton involved in transmitting forces across these structures. Vinculin is also subject to a head-tail autoinhibitory interaction, existing in at least two states, open and closed[35].

Together, key aspects of vinculin function are determined by the mechanical loads its experiences and its conformation[35]. At FAs, vinculin is recruited to the membrane-proximal integrin compartment in a closed conformation via interactions with Paxillin or to the membrane via PIP2, and it moves upward to engage F-actin via a transition to an open conformation and interaction with Talin[36]. At AJs, vinculin also bridges a membrane-proximal cadherin-catenin compartment and a membrane distal F-actin compartment, and this depends on its transition to an open conformation and interactions with α-Catenin[37].

Therefore, to investigate the effect of vinculin on the transmission of forces across integrin- and cadherin-based molecular linkages, we modeled vinculin's ability to form a second mechanical connection to F-actin. Specifically, we modeled the behavior of vinculin inside mechanical linkages in two states: (1) closed and unable to bear loads in the FA/AJ (potentially bound to PIP2 or another unloaded component), or (2) open and able to bear loads in the FA/AJ (potentially bound to an exposed cryptic binding site in Talin/α-Catenin and F-actin). In these linkages, vinculin is free to bind a VBS on either Talin or α-Catenin with its head domain and F-actin with its tail domain, and thus its ability to bear loads is subject to this binding kinetics. In the context of the S1033-based vinculin regulatory switch, where mutation of S1033 affected vinculin load and conformation but not localization to FAs/AJs, the unloadable and loadable states correspond to the phosphorylated and unphosphorylated states of vinculin, respectively. As we experimentally observed spatial variations in vinculin conformation and loading at both FAs and AJs, we assessed the effect of varying the fraction of loadable vinculin, $\rho_{Vcl}$. Based this fraction, the number of linkages in the Reinforced and Non-Reinforced States are defined as follows:

$$N_{Link,Reinforced} = [\rho_{Vcl} N_{Link}] \tag{16}$$

$$N_{Link,Non-Reinforced} = N_{Link} - N_{Link,Reinforced} \tag{17}$$

where brackets denote rounding to the nearest integer.

**E. Binding/Unbinding Dynamics of Vinculin in Integrin-based Linkages**

Here, we describe the state variables and transition kinetics for vinculin in integrin-based linkages. Vinculin reinforces an integrin linkage by binding at its head to Talin (Vcl:Talin bond) and its tail to F-actin (Vcl:F-actin bond), and it reinforces a cadherin linkage by binding at its head to α-Catenin (Vcl:α-Catenin bond) and its tail to F-actin (Vcl:F-actin bond). As such, vinculin provides an additional connection to the F-actin interface when it is bound at both ends.

In both integrin and cadherin linkages, the Vcl:F-actin bond is modeled as a two-state catch-slip bond based on single molecule experiments with T12 vinculin (constitutively active) and F-actin[7]. The state variable for the Vcl:F-actin bond in the jth linkage is given by:

$$\theta_j^{Vcl:Factin} = \begin{cases} 0 & Unbound\ State \\ 1 & Weak\ Bound\ State \\ 2 & Strong\ Bound\ State \end{cases} \tag{18}$$

In the Vcl:F-actin bond, binding only occurs from the unbound to weak bound state (0→1) with the rate constant $k_{01}^{Vcl:Factin}$, as in a previous model of vinculin with the two-state catch-slip bond[7]. Rate constants for unbinding from the weak, $k_{10}^{Vcl:Factin}(F)$, and strong states, $k_{20}^{Vcl:Factin}(F)$, and interconversion between the weak and strong states, $k_{12}^{Vcl:Factin}(F)$ and $k_{21}^{Vcl:Factin}(F)$, are defined in Table 2.

In the integrin linkage, the Vcl:Talin bond is modeled as a one state slip bond based on single molecule experiments with vinculin head domain (Vh) and Talin VBS[24]. The state variable for the Vcl:Talin bond in the jth linkage is given by:

$$\theta_j^{Vcl:Tal} = \begin{cases} 0 & Unbound\ State \\ 1 & Bound\ State \end{cases} \tag{19}$$

In the Vcl:Talin bond, binding (0→1) occurs with the with the force-independent rate constant $k_{01}^{Vcl:Tal}$ and unbinding (1→0) occurs with the force-dependent rate constant function $k_{10}^{Vcl:Tal}(F)$ defined in Table 2, where $F$ denotes the force across the bond. When bound at both the Vcl:Tal and Vcl:F-actin bonds, vinculin provides an additional connection to the F-actin that is parallel to the Tal:F-actin bond. As such, the linkage can be engaged via bonds though Talin, Vinculin, or both, and all bonds to the F-actin interface must be broken for the linkage to become disengaged. As such, the general expression for the engagement state of the jth linkage becomes:

$$\Theta_j = \begin{cases} 1 & \theta_j^{Intg:FN} > 0 \wedge \left(\theta_j^{Tal:Factin} > 0 \vee \left(\theta_j^{Vcl:Factin} > 0 \wedge \theta_j^{Vcl:Tal} > 0\right)\right) \\ 0 & elsewhere \end{cases} \tag{20}$$

In the vinculin-reinforced linkage, the Talin:F-actin and Vcl:F-actin bonds form parallel connections to the same location on F-actin. As such, when forces are transmitted across both vinculin and Talin, they are shared equally between the Talin:F-actin and Vcl:F-actin bonds. We modeled equal load sharing between Talin and Vcl because it is not known how the forces inside the integrin-based linkages are distributed. Together, general expressions for the force across each bond in a linkage are defined below.

$$F_{Link,j}^{Intg:FN} = F_{Link,j} \tag{21}$$

$$F_{Link,j}^{Tal:Factin} = \begin{cases} F_{Link,j} & \theta_j^{Tal:Factin} > 0 \ \wedge \left(\theta_j^{Vcl:Factin} = 0 \vee \theta_j^{Vcl:Tal} = 0\right) \\ \dfrac{F_{Link,j}}{2} & \theta_j^{Tal:Factin} > 0 \ \wedge \left(\theta_j^{Vcl:Factin} > 0 \wedge \theta_j^{Vcl:Tal} > 0\right) \\ 0 & \theta_j^{Tal:Factin} = 0 \end{cases} \tag{22}$$

$$F_{Link,j}^{Vcl:Factin} = F_{Link,j}^{Vcl:Tal} = \begin{cases} F_{Link,j} & \left(\theta_j^{Vcl:Factin} > 0 \wedge \theta_j^{Vcl:Tal} > 0\right) \wedge \theta_j^{Tal:Factin} = 0 \\ \dfrac{F_{Link,j}}{2} & \left(\theta_j^{Vcl:Factin} > 0 \wedge \theta_j^{Vcl:Tal} > 0\right) \wedge \theta_j^{Tal:Factin} = 0 \\ 0 & \theta_j^{Vcl:Factin} = 0 \vee \theta_j^{Vcl:Tal} = 0 \end{cases} \tag{23}$$

As with the unreinforced integrin linkage, when all bonds at either interface break, the linkage returns to a disengaged state, in which it experiences no extension and bears no force. If vinculin head unbinds Talin, the linkage is free to recruit another vinculin molecule.

#### F. Binding/Unbinding Dynamics of Vinculin in Cadherin-based Linkages

Here, we describe the state variables and transition kinetics for vinculin in cadherin-based linkages. The behavior of vinculin in the cadherin linkage is modeled the same as in the integrin linkage, except with different kinetic parameters for the Vcl:α-Catenin bond. The Vcl:F-actin bond is modeled as described in the previous section, identical to in the integrin-based linkage.

The Vcl:α-Catenin bond is modeled as a one state slip bond based on single molecule experiments with vinculin head domain (Vh) and α-Catenin VBS[24]. The state variable for the Vcl:α-Catenin bond in the jth linkage is given by:

$$\theta_j^{Vcl:aCat} = \begin{cases} 0 & Unbound\ State \\ 1 & Bound\ State \end{cases} \quad (24)$$

In the Vcl:α-Catenin bond, binding (0→1) occurs with the with the force-independent rate constant $k_{01}^{Vcl:aCat}$ and unbinding (1→0) occurs with the force-dependent rate constant function $k_{10}^{Vcl:aCat}(F)$ defined in Table 3, where $F$ denotes the force across the bond.

Overall, when bound at both the Vcl:α-Catenin and Vcl:F-actin bonds, vinculin provides an additional connection to the F-actin that is parallel to the α-Catenin:F-actin bond. As such, linkage can be engaged via bonds though α-Catenin, Vinculin, or both, and all bonds to the F-actin interface must be broken for the linkage to become disengaged. As such, the general expression for the engagement state of the jth linkage becomes:

$$\Theta_j = \begin{cases} 1 & \theta_j^{Cad:Cad} > 0 \wedge \left(\theta_j^{aCat:Factin} > 0 \vee \left(\theta_j^{Vcl:Factin} > 0 \wedge \theta_j^{Vcl:aCat} > 0\right)\right) \\ 0 & elsewhere \end{cases} \quad (25)$$

In the vinculin-reinforced linkage, the α-Catenin:F-actin and Vcl:F-actin bonds form parallel connections to the same location on F-actin. As such, when forces are transmitted across both vinculin and α-Catenin, they are shared equally between the α-Catenin:F-actin and Vcl:F-actin bonds. We modeled equal load sharing between α-Catenin and Vcl because it is not known how the forces inside the cadherin-based linkages are distributed. Together, general expressions for the force across each bond in a linkage are defined below.

$$F_{Link,j}^{Cad:Cad} = F_{Link,j} \quad (26)$$

$$F_{Link,j}^{aCat:Factin} = \begin{cases} F_{Link,j} & \theta_j^{aCat:Factin} > 0 \ \wedge \left(\theta_j^{Vcl:Factin} = 0 \vee \theta_j^{Vcl:aCat} = 0\right) \\ \dfrac{F_{Link,j}}{2} & \theta_j^{aCat:Factin} > 0 \ \wedge \left(\theta_j^{Vcl:Factin} > 0 \wedge \theta_j^{Vcl:aCat} > 0\right) \\ 0 & \theta_j^{aCat:Factin} = 0 \end{cases} \quad (27)$$

$$F_{Link,j}^{Vcl:Factin} = F_{Link,j}^{Vcl:aCat} = \begin{cases} F_{Link,j} & \left(\theta_j^{Vcl:Factin} > 0 \wedge \theta_j^{Vcl:aCat} > 0\right) \ \wedge \theta_j^{aCat:Factin} = 0 \\ \dfrac{F_{Link,j}}{2} & \left(\theta_j^{Vcl:Factin} > 0 \wedge \theta_j^{Vcl:aCat} > 0\right) \ \wedge \theta_j^{aCat:Factin} = 0 \\ 0 & \theta_j^{Vcl:Factin} = 0 \vee \theta_j^{Vcl:aCat} = 0 \end{cases} \quad (28)$$

As with the unreinforced cadherin-catenin linkage, when all bonds at either interface break, the linkage returns to a disengaged state, in which it experiences no extension and bears no force. If vinculin head unbinds α-Catenin, the linkage is free to recruit another vinculin molecule. Lastly, we assumed that integrin- and cadherin-based linkages have a homogenous spring constant and that incorporation of vinculin into large integrin- or cadherin-based molecular linkages does not alter the spring constant of the linkage, $K_{Link}$.

**G. Ensemble Vinculin Molecular Tension**

To facilitate comparisons to experimental measurements with FRET-based vinculin MTS (VinTS), we measured in simulations the ensemble vinculin molecular tension, $\langle F_{Vcl} \rangle_{Links}$, defined as the average of tension across the vinculin molecules in all linkages:

$$\langle F_{Vcl} \rangle_{Links} = \frac{1}{N_{Link}} \sum_{i=1}^{N_{Link}} F_{Link,i}^{Vcl} \quad (29)$$

where the force across the vinculin molecule in the jth linkage is equal to the force across the Vcl:Factin bond, $F_{Link,j}^{Vcl} = F_{Link,j}^{Vcl:Factin}$ , which is also the same as the force across the Vcl:Talin or ($F_{Link,j}^{Vcl:Factin} = F_{Link,j}^{Vcl:Tal}$) or Vcl: α-Catenin bond ($F_{Link,j}^{Vcl:Factin} = F_{Link,j}^{Vcl:aCat}$) in that linkage as described earlier. As vinculin is able to bind actin and bear loads in reinforced linkages but not in non-reinforced linkages, the ensemble vinculin molecular tension is approximately equal to: $\langle F_{Vcl} \rangle_{Links} \approx \rho_{Vcl} \cdot \langle F_{Vcl} \rangle_{Reinforced\ Links} + (1 - \rho_{Vcl}) \cdot 0$.

**H. Simulation Algorithm**

Stochastic simulations of the FA and AJ clutch models were carried out using the Gillespie Stochastic Simulation Algorithm as previously described for clutch models[13]. It was verified that results with this simulation algorithm matched those obtained using a discrete time-step algorithm with a small time-step (0.0005 s). The simulations were started with all linkage bonds in the unbound state and initial connections were allowed to form subject to the binding rate constants listed in Table 1. Simulations were run for a duration of 1000s. Outputs (such as total force and engaged linkage fraction) were computed as time-averages over a time window after reaching steady-state. To compute mean engagement lifetime and fraction of disengagement events due to unbinding at certain interfaces, all linkage state transitions after reaching steady state were recorded and analyzed. Single linkage simulations were conducted at constant loading rates using a discrete time-step algorithm with a time-step of 0.0001 sec. Similar analyses to those described for the FA and AJ clutch models were conducted on these simulations.

**I. Single Component Friction Clutch Configuration for Validation of Model Implementation**

Implementation of the friction clutch models was validated by comparing simulations to analytical expressions of the friction-velocity relationship for simple linkages with ideal or slip bonds derived by Sens[5]. To conduct these simulations, linkages were set to bind/unbind at only one interface according to an ideal, $k_{off}(F) = k_{off}$, or Bell slip bond, $k_{off}(F) = k_{off,0}e^{F/F_b}$. Other parameters for the validation simulations were set to match the base parameters for the FA and AJ friction clutch models ($N_{link}$=50, $K_{link}$= 5 pN/nm, $k_{on}$=2/s) and the same range of velocities was also used ($v$ =[0.1,100] nm/s). Simulations were conducted in the limit of an infinitely rigid external spring ($K_{Ext} \gg N_{link}K_{link}$), where the following analytical expressions apply. The force-velocity relationship for the case of an ideal bond, $k_{off}(F) = k_{off}$, is given in Equation 1 of Sens[5]:

$$\langle F \rangle = \left[ \frac{N_{link}K_{link}k_{on}}{(k_{on} + k_{off})k_{off}} \right] v \quad (30)$$

The force-velocity relation for the case of a Bell slip bond, $k_{off}(F) = k_{off,0}e^{F/F_b}$, is given in Equation 6 of Sens[5]:

$$\langle F\rangle = \frac{N_{link}F_b\left(\frac{k_{on}}{k_{off,0}}\right)e^{\frac{v_\beta}{v}}\int_0^\infty f e^{-e^{\frac{f v_\beta}{v}}}df}{\frac{v}{v_\beta}+\left(\frac{k_{on}}{k_{off,0}}\right)e^{\frac{v_\beta}{v}}\int_{\frac{v_\beta}{v}}^\infty \frac{e^{-f}}{f}df}, \qquad v_\beta = \frac{k_{off,0}F_b}{K_{link}} \tag{31}$$

Lastly, as integrin- and cadherin-based adhesions are known to contain multiple proteins with catch bonds, we also considered a two-pathway catch-slip bond, $k_{off}(F) = k_{off,0}\left(0.9e^{-2F/F_b} + 0.1e^{F/F_b}\right)$, in the single component friction clutch as a tool to build intuition.

**J. Determination of External Spring Constant for given Effective Young's Modulus**

In clutch models, an external spring with a defined spring constant is used to represent the stiffness of the external material to which the molecular linkages are connected. For the FA model, this corresponds to the substrate stiffness. For the AJ model, this corresponds to the stiffness of the adjacent cell in the monolayer, or in the context of experiments measuring traction forces on cadherin-coated elastic substrates, this corresponds to the substrate stiffness. To relate the external spring constant, $K_{ext}$ [pN/nm], to an equivalent Young's modulus, $E_{eff}$ [kPa], we used a relationship from previous motor-clutch models, $E_{eff} = \frac{9K_{Ext}}{4\pi a}$, where $a$ is the radius of an equivalent circular adhesion[14,38]. For the AJ model, the adhesion radius was estimated using the circular adhesion areas corresponding to the base number of linkages ($N_{Link} = 50$) at the reported density of cadherin molecules in cadherin clusters[20] (2666/μm$^2$), corresponding to adhesion radius of 77 nm. This yielded a conversion factor $E_{eff}/K_{ext}$ of 9.27 kPa/(pN/nm). For the AJ model, we used this conversion factor to approximate the external spring constant from experimental estimates of the elastic modulus of cell monolayers[22,23] (~20-33 kPa or ~2.16-3.56 pN/nm). For the FA model, the external spring constant was set arbitrarily high to represent experimental conditions from this paper (stiffness of glass), and therefore no conversion factor was needed for the FA model.

**K. Plots of Mean Bond Lifetime versus Force**

To overlay unbinding rate constant models on mean bond lifetime versus force plots (Fig S1c,f), mean lifetimes were obtained from the force-dependent rate constant models as follows. For single bound state models, where there is a single unbinding transition, the lifetime at force F was taken to be the inverse of the rate constant at force F. For two bound state catch bond models, where there are unbinding transitions from weak and strong states, as well as interconversion between them, the mean lifetime was obtained by assuming an initial distribution of bound states based on equilibrium under no force, as previously done for this type of bond[12]. Then, the mean lifetime was solved for using a continuous time Markov chain to calculate the mean exit times from each bound state and then weighting them according to the starting distribution for the two bound states.

#### III. RESULTS

**A. Friction Clutch with Single-Component Linkage: Technical validation of model implementation and comparison of individual linkage dynamics to total friction.**

To validate implementation of the friction clutch model and to gain intuition about molecular determinants of friction, we used the previously developed friction clutch model that contained simple linkages with a single bond (Fig S2a and Section II-I). The single bond was modeled as an ideal bond, Bell model slip bond, or two pathway catch-slip bond (Fig S2b-c).

To validate our implementation of the friction clutch model, we simulated clutches comprised of linkages with ideal or slip bonds and compared them to previously determined analytical expressions relating average total friction force to speed (from Sens[5] and given in Equations 30 and 31). For the ideal bond, the friction force increased linearly with speed, and the simulation results for two different values of the unbinding rate constant both showed good agreement with the previously obtained analytical relationship from Sens (Fig S2d). For the slip bond, the friction force first increased then decreased with speed, and the simulation results for three combinations of unbinding rate parameters all showed good agreement with the analytical relationship from Sens (Fig. S2g). Taken together, these simulations validated our implementation of the friction clutch model.

We next sought to gain intuition about molecular determinants of friction. To do so, we compared individual linkage dynamics in the clutch, using mean engagement lifetime, and an effective friction coefficient (F/V), a standard parameter describing the frictional resistance between sliding surfaces (cell-ECM or cell-cell) in models of CCM[4]. For the ideal bond, the engagement lifetime (Fig S2e) and effective friction coefficient (Fig S2f) were both independent of speed. For the slip bond, both the engagement lifetime (Fig S2h) and effective friction coefficient (Fig S2i) were speed-dependent, decreasing monotonically with speed. As integrin- and cadherin-based adhesions are known to contain multiple proteins with catch bonds, we also investigated a catch-slip bond. For the catch-slip bond, both the engagement lifetime (Fig S1k) and effective friction coefficient (Fig S2l) were again speed-dependent, but this time increasing up to an intermediate speed then decreasing. Therefore, for each bond type, the qualitative shape of the effective friction coefficient-speed curve relates to the dynamics of individual linkages, meaning the underlying force-sensitive dynamics are indicative of the larger-scale mechanics of the friction clutch.

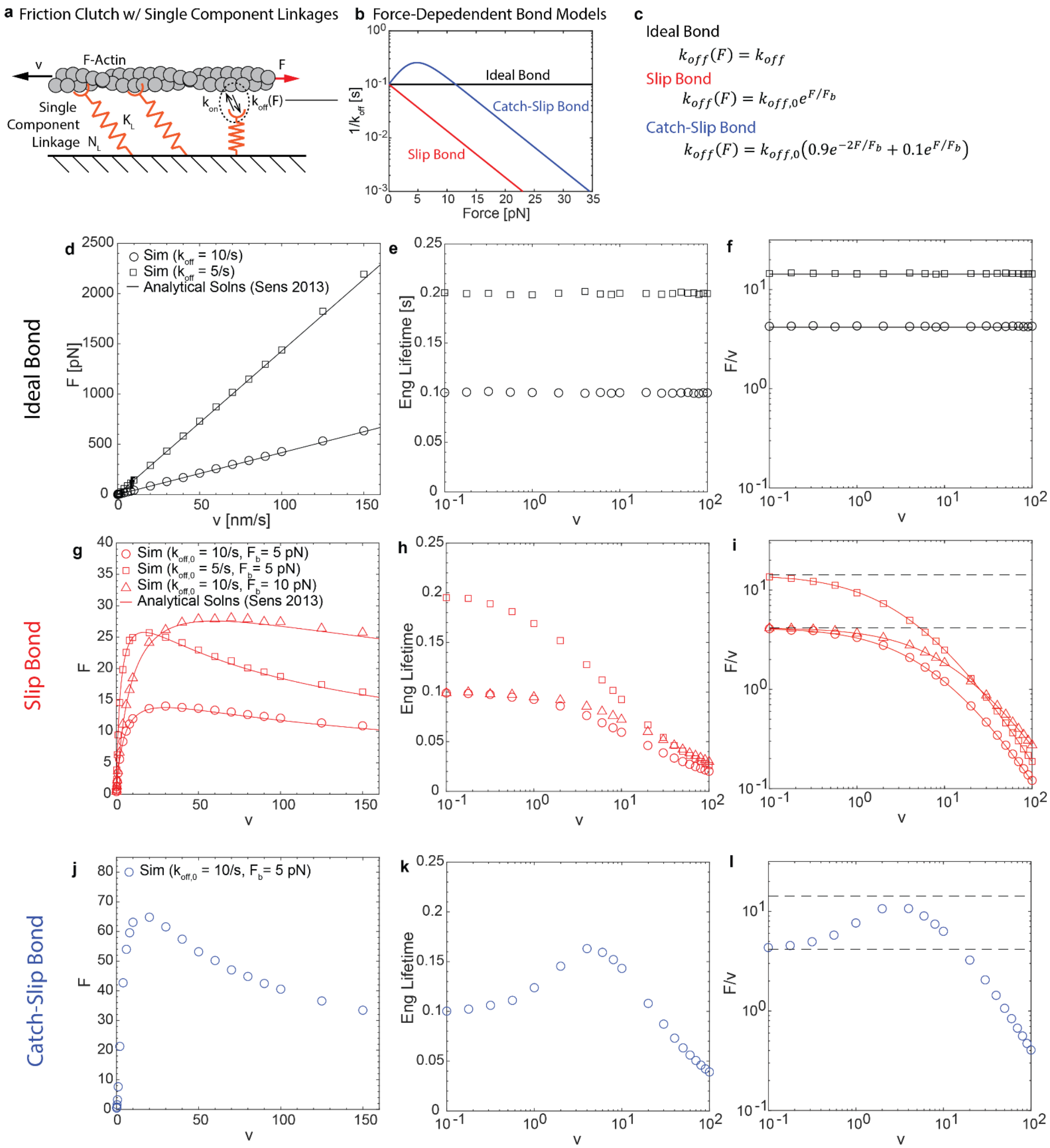


**Fig. S2 Validation of Friction Clutch Model with Single Component Linkages**. (a) Schematic of friction clutch model with single component linkages containing a single binding/unbinding interface. (b) Plot of mean bond lifetime versus force for three generic force-dependent bond models used for the unbinding rate constant for single component linkages. (c) Unbinding rate constant functions for the three generic force-dependent bond models used for single component linkages. (d-f) Plots of friction force, mean linkage engagement lifetime, and effective friction coefficient (F/V) for friction clutch containing single component linkages with ideal bonds, for two different unbinding rate constants). (g-i) Same for friction clutch containing single component linkages with a slip bond, for three different unbinding rate constant parameter combinations. (j-l) Same for friction clutch containing single component linkages with a catch-slip bond. For (d-l), data points are time averages from simulations and overlaid lines are analytical solutions (Equation 30 for ideal bond case and Equation 31 for slip bond case) from Sens 2013.

**B. Force sensitivity of multi-component integrin- and cadherin-based linkages.**

The molecular linkages that mediate mechanical connections in the FA/AJ between the actin cytoskeleton and the ECM/adjacent cell are composed of multiple proteins and therefore multiple dynamic interfaces. As the single bond linkages used in the previous friction clutch model (Section III-A) do not capture the complex connectivity and potential regulation of these biological linkages, we developed multi-component linkages to use in the frictional clutch models. These multi-component linkages consisted of integrin:talin:F-actin in FAs (Fig S1b) or E-cadherin:β-catenin:α-catenin:F-actin in AJs (Fig S1e), which could be reinforced through the incorporation of vinculin, and whose ability to bear force depend on the force-dependent lifetimes of bonds in the integrin-based (Fig S1c, Section II-B/D, and Table 2) or cadherin-based (Fig S1f, Section II-C/E, and Table 3) linkages.

To transmit forces, molecular linkages must be bound at both sides (actin and external interfaces in our model). We note that the lifetimes of the Intg:FN and Talin:F-actin bonds (Fig S1c), as well as those of the E-cad:E-cad and α-catenin:F-actin bonds (Fig S1f), are within an order of magnitude of each other over a wide range of physiological molecular loads. This suggests that the force-sensitive dynamics of both linkages are not driven by a single bond but rather are due to the combined behavior of multiple dynamic interfaces. Also, there are force regimes in both linkages over which the primary bond to the actin interface (Talin:F-actin or α-catenin:F-actin) has a shorter lifetime than bonds to the external interface, suggesting that reinforcement of the actin interface with a parallel bond, such as the Vcl:F-actin bond, could modulate the combined dynamics of the linkage. Therefore, the force-sensitivity and regulation of integrin- and cadherin-based linkages will depend on the combined behavior of multiple dynamic interfaces, not just a single bond in the linkage.

As parameters for force-sensitive bond dynamics were obtained from single molecule experiments characterizing the interfaces separately, we first assessed their suitability for use in combination to model multi-component linkages at the FA and AJ. To do so, we subjected single linkages to fixed loading rates (Fig S3a) based on estimates of protein loading rates inside cells (using typical protein stiffnesses ~1-10 pN/nm and actin flow speeds[16], ~2-600 nm/s, or cell migration speeds in monolayers[15], ~1-30 um/hr or ~0.28-8.33 nm/s). We quantified the linkage engagement lifetime, which is defined as the mean time linkages remain engaged and support force transmission. In the integrin-based linkage, the engagement lifetime was strongly biphasic, first increased then decreasing with loading rate, regardless of vinculin reinforcement (Fig S3b-c). This is consistent with the combined linkage having a strong catch-slip behavior. Vinculin reinforcement did not alter the functional form of the lifetime-load rate curve, but substantially increased the engagement lifetime across a wide range of low and intermediate loading rates (up to ~56 pN/s). This is consistent with the established role of vinculin as a mechanical reinforcement element at the FA. In the cadherin-based linkage, the engagement lifetime remained constant or increased slightly with loading rate, before decreasing strongly at higher loading rates (Fig S3d-e). This is consistent with the combined linkage having an ideal-slip or weak catch-slip behavior. Vinculin reinforcement did not alter the functional form of the lifetime-load rate curve, but substantially increased the engagement lifetime across a wide range of low and intermediate loading rates (up to ~100 pN/s). This is consistent with the established role of vinculin as a mechanical reinforcement element at the AJ. We also note that, as expected, the behavior of the multi-component linkages is not readily predictable from the force-sensitive dynamics of single bonds in the linkages and are not dominated by a single interface.

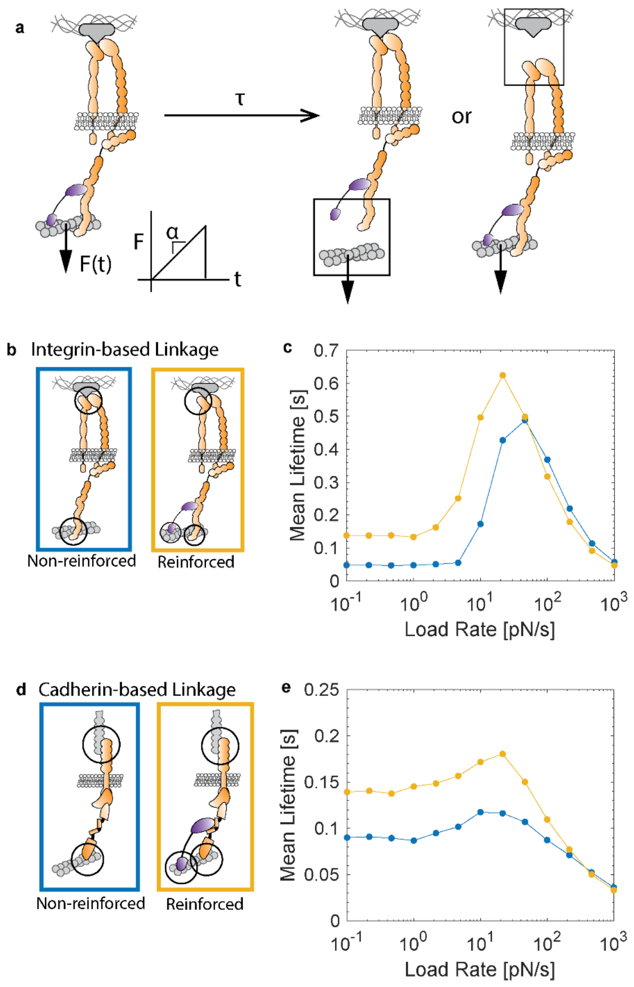


**Fig. S3 Characterization of Multi-component Integrin- and Cadherin Linkages**. (a) Schematic of ramp load (constant load rate) applied to a single integrin-based linkage leading to its disengagement once all bonds at either the external or actin interface are broken. (b-c) Plot of mean engagement lifetime versus load rate for integrin-based linkage without and with vinculin reinforcement. (d-e) Plot of mean engagement lifetime versus load rate for cadherin-based linkage without and with vinculin reinforcement.

**C. FA Friction Clutch and Effect of Vinculin Reinforcement.**

We next sought to assess regulators of cell-ECM friction mediated by the FA. To do so, we incorporated multi-component integrin linkages into the friction clutch model and assessed the effect of increasing the fraction of linkages with loadable vinculin (Fig S4). Time-traces for the number of engaged linkages and total force exhibit fluctuations (Fig S4c-f), consistent with the behavior of previous frictional clutch models[5]. In the absence of vinculin reinforcement ($\rho_{Vcl} = 0$), the mean linkage engagement lifetime (Fig S4g) and mean linkage engagement fraction (Fig S4h) had strongly biphasic relationships with speed, first increasing then decreasing. This was consistent with the strong catch-slip behavior of single integrin-based linkages (Section III-B). Furthermore, the effective friction coefficient (F/V) (Fig S4j) exhibited a similar biphasic relationship with speed. Therefore, as observed for friction clutch models with a single bond (Section III-A), the force-sensitive dynamics of the multi-component integrin-based linkage was predictive of the shape of the friction coefficient-speed relationship.

We next assessed the effect of vinculin mechanical reinforcement on the FA friction clutch by increasing the fraction of linkages with loadable vinculin ($\rho_{Vcl}$). Increasing the fraction of loadable vinculin substantially increased the mean linkage engagement lifetime (Fig S4g) and mean linkage engagement fraction (Fig S4h) over the range of speeds corresponding the monolayer velocities observed in this work and elsewhere[15], ~1-30 um/hr or ~0.28-8.33 nm/s. This was consistent with vinculin's effect on the dynamics of single integrin-based linkages being more pronounced at low and intermediate loading rates (Section III-B). As a result, vinculin reinforcement also significantly increased the friction force (Fig S4i) and effective friction coefficient (Fig S4j) across this range speeds in our system. At higher speeds, the friction coefficient strongly decreased with speed regardless of vinculin reinforcement, and vinculin reinforcement did not alter the functional forms of the engagement lifetime-speed or friction coefficient-speed relationships. Time-traces contain a significant number of Vcl:F-actin bonds in the strong bound state (Fig S4e-f), indicating engagement of the vinculin's catch bond in the FA model.

To further assess the ability of the vinculin regulatory switch to tune friction, we assessed the effect of finer variations in the fraction of loadable vinculin at an intermediate speed (10 um/hr) in the CCM range (Fig S4k). The friction coefficient varied approximately linearly with the fraction of loadable vinculin and had a maximum of effect of ~2-fold increase in the effective friction coefficient. To facilitate comparisons to experimental measurements with FRET-based vinculin MTS (VinTS), we also assessed ensemble vinculin molecular tension, $\langle F_{Vcl} \rangle_{Links}$. Ensemble vinculin molecular tension increased approximately linearly with the amount of loadable vinculin, meaning it varied in a similar fashion to the friction coefficient.

Overall, analysis of the FA friction clutch model demonstrates the relationship between dynamics of multi-component integrin linkages and subcellular friction at the FA and suggests that vinculin reinforcement increases cell-substrate friction in a tunable manner across a range of speeds corresponding to collective cell migration.

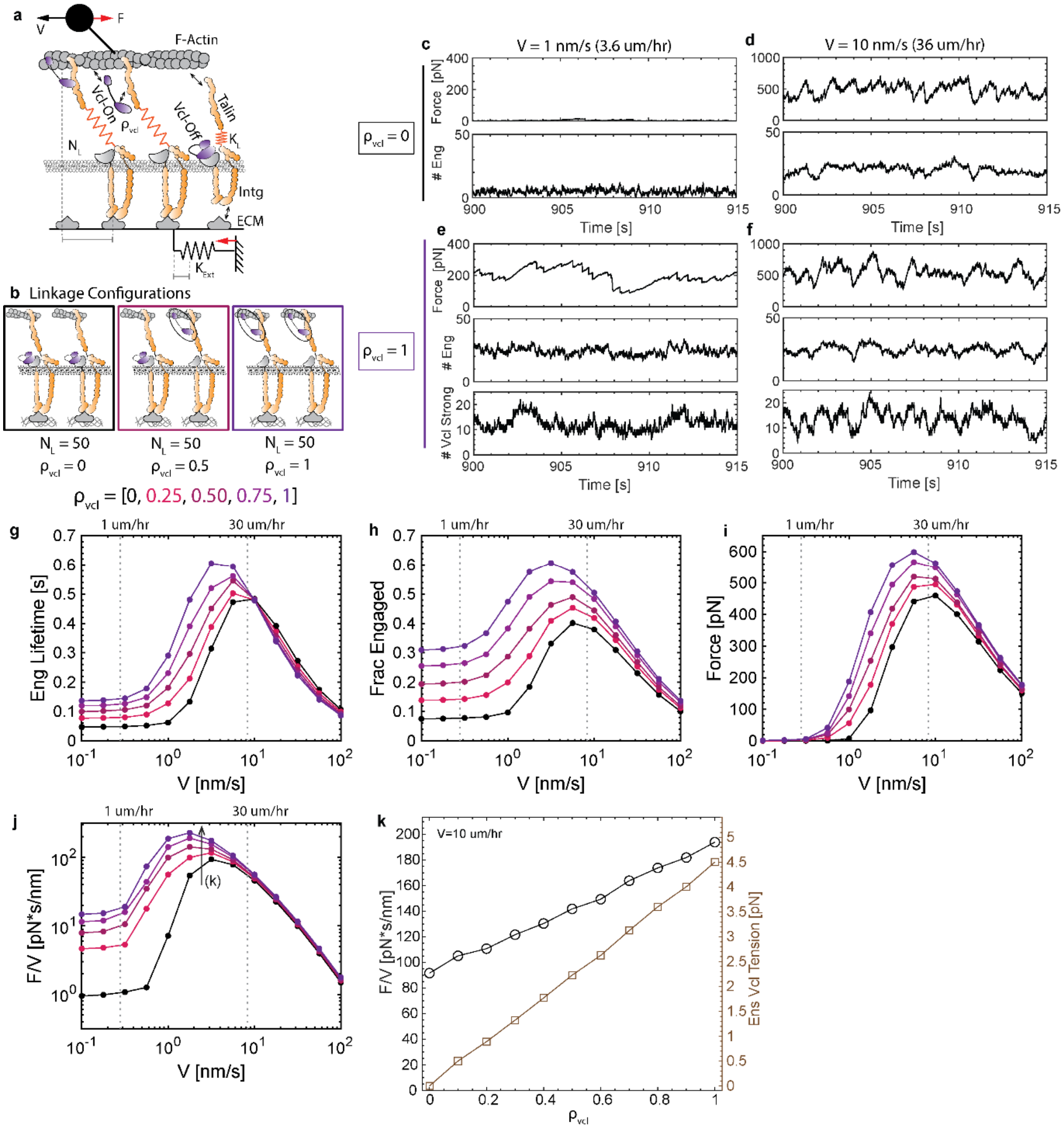


**Fig. S4 FA Friction Clutch Model**. (a) Schematic of FA friction clutch model. (b) Schematic of linkage configurations for different values of $\rho_{Vcl}$, the fraction of linkages with loadable vinculin. (c-d) Time-traces of total friction force and number of engaged linkages for simulations with $\rho_{Vcl} = 0$ and speed ($v$) 1 nm/s or 10 nm/s. (e-f) Time-traces of total friction force, number of engaged linkages, and number of linkages with the Vcl:F-actin bond in the strong bound state for simulations with $\rho_{Vcl} = 1$ and speed ($v$) 1 nm/s or 10 nm/s. (g-j) Plots of mean linkage engagement lifetime, mean fraction of linkages engaged, mean total friction force, and mean effective friction coefficient (F/v) for 5 values of

$\rho_{Vcl}$. (k) Plot of effective friction coefficient (F/V) (left y axis) and ensemble vinculin molecular tension (right y axis) versus $\rho_{Vcl}$ for speed ($v$) 10 um/hr.

**D. AJ Friction Clutch and Effect of Vinculin Reinforcement**

We next sought to assess regulators of cell-cell friction mediated by the AJ. To do so, we incorporated multi-component integrin linkages into the friction clutch model and assessed the effect of increasing the fraction of linkages with loadable vinculin (Fig S5). Time-traces for the number of engaged linkages and total force exhibit fluctuations (Fig S5c-f), consistent with the behavior of previous frictional clutch models[5]. In the absence of vinculin reinforcement ($\rho_{Vcl} = 0$), the mean linkage engagement lifetime (Fig S5g) and mean linkage engagement fraction (Fig S5h) had weak biphasic relationships with speed, increasing slightly or remaining constant at lower speeds and then decreasing at higher speeds. This was consistent with the weak catch-slip or ideal-slip behavior of single cadherin-based linages (Section III-B). Furthermore, the effective friction coefficient (F/V) (Fig S5j) exhibited a similar relationship with speed. Therefore, as observed for friction clutch models with a single bond (Section III-A), the force-sensitive dynamics of the multi-component cadherin-based linkage was predictive of the shape of the friction coefficient-speed relationship.

We next assessed the effect of vinculin mechanical reinforcement on the AJ friction clutch by increasing the fraction of linkages with loadable vinculin ($\rho_{Vcl}$). Increasing the fraction of loadable vinculin substantially increased the mean linkage engagement lifetime (Fig S5g) and mean linkage engagement fraction (Fig S5h) over the range of relative cell-cell speeds corresponding to monolayer velocities observed in this work and elsewhere[15], ~1-30 um/hr or ~0.28-8.33 nm/s. This was consistent with vinculin's effect on the dynamics of single cadherin-based linkages being more pronounced at low and intermediate loading rates (Section III-B). As a result, vinculin reinforcement also significantly increased the friction force (Fig S5i) and effective friction coefficient (Fig S5j) across this range speeds in our system. At higher speeds, the friction coefficient strongly decreased with speed regardless of vinculin reinforcement, and vinculin reinforcement did not alter the functional forms of the engagement lifetime-speed or friction coefficient-speed relationships. Time-traces contain significant numbers of α-catenin:F-actin and Vcl:F-actin bonds in the strong bound state (Fig S5e-f), indicating engagement of the α-catenin catch bond to a lesser extent and the vinculin catch bond to a larger extent in AJ model.

To further assess the ability of the vinculin regulatory switch to tune friction, we assessed the effect of finer variations in the fraction of loadable vinculin at an intermediate speed (10 um/hr) in the CCM range (Fig S5k). The friction coefficient varied approximately linearly with the fraction of loadable vinculin and had a maximum of effect of ~4-fold increase in the effective friction coefficient. To facilitate comparisons to experimental measurements with FRET-based vinculin MTS (VinTS), we also assessed ensemble vinculin molecular tension, $\langle F_{Vcl} \rangle_{Links}$. Ensemble vinculin molecular tension increased approximately linearly with the amount of loadable vinculin, meaning it varied in a similar fashion to the friction coefficient.

Together, analysis of the AJ friction clutch model demonstrates the relationship between dynamics of multi-component cadherin linkages and subcellular friction at the AJ and suggests that vinculin reinforcement increases cell-cell friction in a tunable manner across a range of speeds corresponding to collective cell migration. Compared to FA model, ensemble vinculin tension at the AJ was lower, but the effect of vinculin reinforcement on the friction coefficient was overall higher in AJs (~4-fold increase) than in FAs (~2-fold increase).

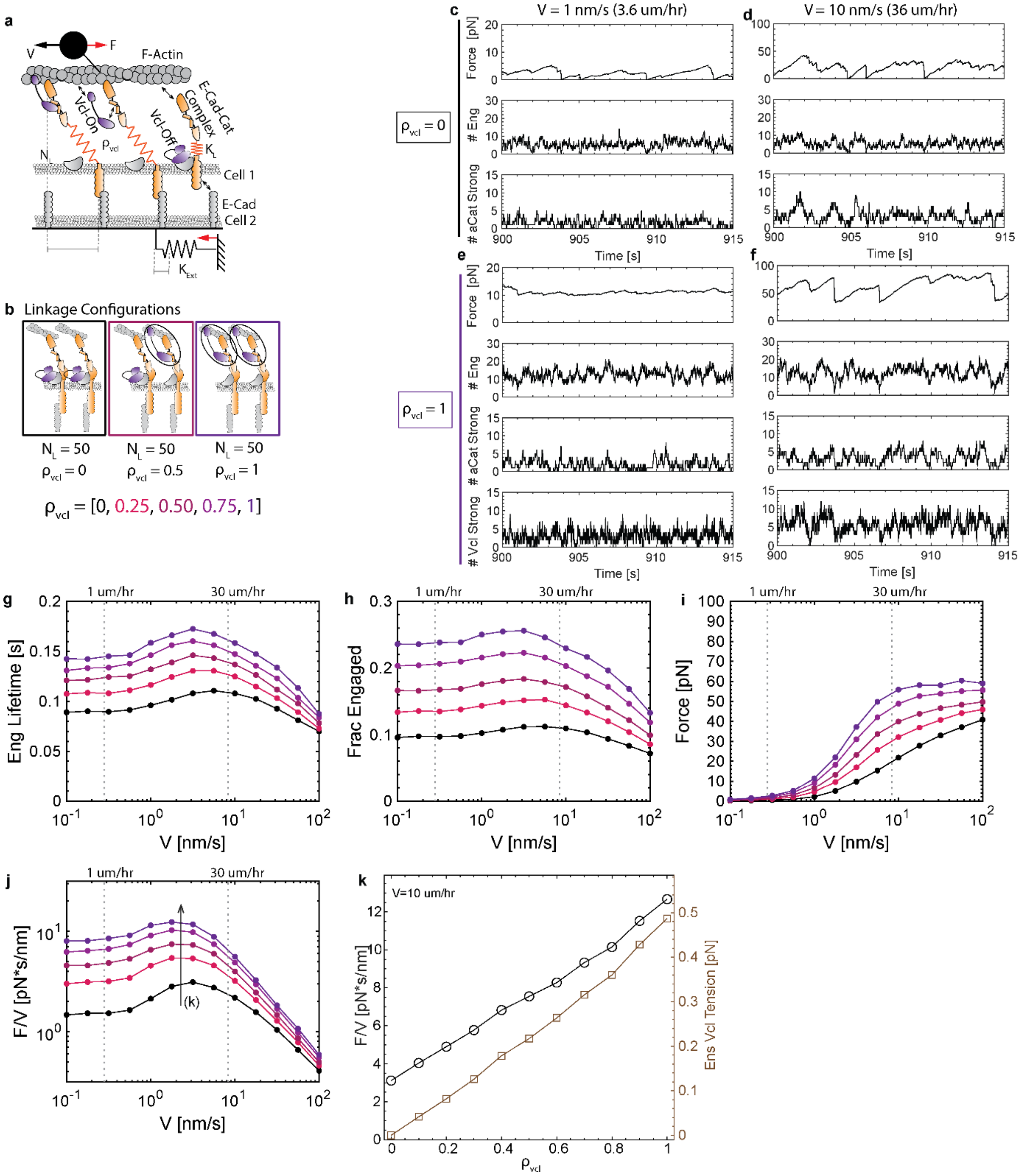

**Fig. S5 AJ Friction Clutch Model**. (a) Schematic of AJ friction clutch model. (b) Schematic of linkage configurations for different values of $\rho_{Vcl}$, the fraction of linkages with loadable vinculin. (c-d) Time-traces of total friction force, number of engaged linkages, and number of linkages with the α-catenin:F-actin bond in the strong bound state for simulations with $\rho_{Vcl} = 0$ and speed ($v$) 1 nm/s or 10 nm/s. (e-f) Time-traces of total friction force, number of engaged linkages, and number of linkages with the α-catenin:F-actin and Vcl:F-actin bond in the strong bound state for simulations with $\rho_{Vcl} = 1$ and speed ($v$) 1 nm/s or 10 nm/s. (g-j) Plots of mean linkage engagement lifetime, mean fraction of linkages engaged, mean total friction force, and mean effective friction coefficient (F/v) for 5 values of

$\rho_{Vcl}$. (k) Plot of effective friction coefficient (F/V) (left y axis) and ensemble vinculin molecular tension (right y axis) versus $\rho_{Vcl}$ for speed ($v$) 10 um/hr.

**E. Robustness of vinculin-reinforcement as a mechanism to tune friction at FA and AJ.**

We next sought to assess the robustness of the vinculin-based reinforcement mechanism across a wide parameter space in the FA and AJ friction clutch models. To do so, we varied parameters separately across a range of two orders of magnitude centered on the base value and obtained the relative change in total force transmission between the model with no vinculin reinforcement ($\rho_{vcl} = 0$) and full vinculin reinforcement ($\rho_{vcl} = 1$) (Fig S6). While the magnitude of vinculin's effect increased or decreased in response to changes in some of the parameters, across the 40 parameter combinations tested we found that the vinculin reinforcement resulted in at least a 20% increase in friction force for 36/40 parameter combinations at the FA (Fig S6a) and 40/40 parameter combinations at the AJ(Fig S6b), with ~100% (2-fold) or greater increases for many parameter combinations in the FA model and ~300% (4-fold) or greater increases for many parameter combinations in the AJ model. Overall, this indicates the general robustness of vinculin-based reinforcement as a mechanism for turning friction at the FA and AJ.

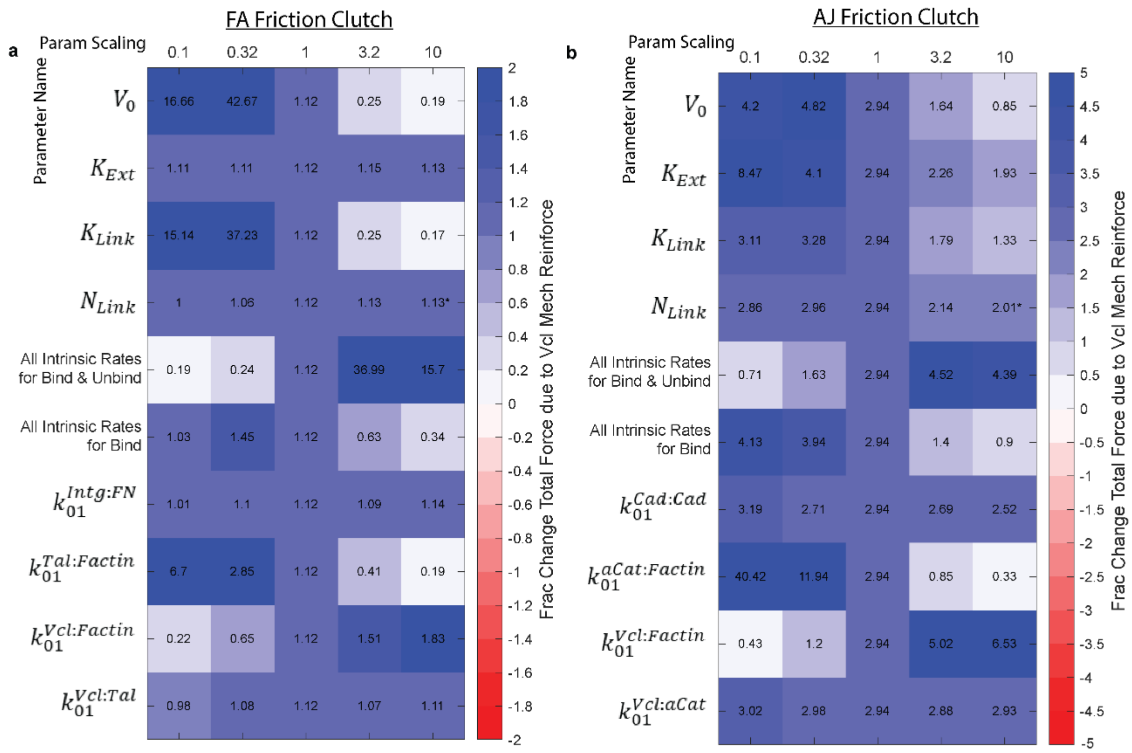


(*) Highest parameter value is 5x instead of 10x for # Linkages.

**Fig. S6 Robustness of Vinculin-based Reinforcement Mechanism.** Parameters in the (a) FA or (b) AJ clutch models were varied separately across a range of two orders of magnitude centered on the base value (0.1x, 0.32x, 1x, 3.2x, or 10x the base value reported in Table 1) to assess the robustness of the effect of loaded vinculin on friction. For each parameter value, the text and color indicate the relative change in mean total friction force between no vinculin reinforcement ($\rho_{vcl} = 0$) and full vinculin reinforcement ($\rho_{vcl} = 1$), $\left(F_{[\rho_{vcl}=1]} - F_{[\rho_{vcl}=0]}\right)/F_{[\rho_{vcl}=0]}$. (A relative change of 1 means that vinculin reinforcement results in a 100% increase in friction.) For linkage number only, the highest parameter value was set to 5x instead of 10x the base value.

#### IV. MODEL ASSUMPTIONS, LIMITATIONS, AND POSSIBLE EXTENSIONS

The FA and AJ friction clutch models share the basic assumptions previously detailed for molecular clutch models[13], except that no assumption on the force-velocity relationship for myosin is required, as a constant velocity based on the motion of the cell with respect to the external surface is used. In the rest of this section, we cover assumptions that are specific to the modeling of multi-bond linkages and vinculin mechanical reinforcement and address potential limitations on the interpretation of related experiments.

The following assumptions are made regarding the mechanical connectivity of vinculin in integrin- and cadherin-based linkages. First, we consider only 1 vinculin binding site (VBS) per linkage. However, Talin contains multiple VBS, and vinculin is also known to bind proteins other than Talin that associate with integrins, such as paxillin, or proteins other than α-Catenin that associate with E-cadherin, such as β-Catenin. These could result in additional vinculin-mediated connections to the actin interface in parallel to the existing actin bonds, resulting in further stabilization of the linkages and potentially increased force transmission. As such, the assumption of one VBS per linkage likely underestimates the effect of loadable vinculin on adhesion-based friction. Second, we assumed that the addition of vinculin to linkages did not alter the effective spring constant of the linkage. This is consistent with large portions of the integrin-/cadherin-based linkages located between the external surface and the VBS (mainly the integrin and cadherin molecules themselves) being softer than that of the back end of Talin or α-Catenin and therefore driving the effective spring constant of the linkage. If, in contrast, the region between the VBS and the actin bond in the linkage was actually softer than the rest of the linkage, reinforcement of this regime with a parallel vinculin connection could increase the effective stiffness of the linkage as well as alter the loading rate of the bonds. This would provide another means by which adapter proteins could alter friction at adhesions, which could be explored further in future models. Third, when both vinculin and Talin or α-Catenin are bound at the actin interface, we assumed that loads are shared equally between the two parallel bonds, as it is not known how the forces inside the integrin- or cadherin-based linkages are distributed. Deviations from this assumption would affect the interplay between the force-sensitive unbinding kinetics of actin bonds in parallel, which could be explored in future models.

Additionally, the following assumptions are made regarding the mechanochemical properties and regulation of vinculin. We modeled vinculin in mechanical linkages existing in one of two states, able or unable to bear loads, and treated the relative fraction of vinculin in each of these states to be tunable (via the parameter $\rho_{Vcl}$). The number of states vinculin exists in at FAs and AJs is likely higher and the regulation of the distribution of vinculin in these states is likely more complex. As the biological mechanisms are just being elucidated, this is an important point for future work. However, as the two state formulation represents the extremes of vinculin mechanical function, our approach likely captures key aspects of the system.

Furthermore, we consider the effect of vinculin due to the addition of a direct load-bearing linkage to F-actin. This is supported by the fact that vinculin molecular tension regulates FA dynamics[39], and that vinculin's effects on traction forces and adhesion strength require the physical coupling between the head and tail domain of full-length vinculin[40]. However, we note that vinculin could additionally affect force transmission at FAs and AJs through indirect effects, including conformational stabilization of the linkage[21,41] as well as the mechanosensitive recruitment of other adhesion or actin-binding proteins. For instance, recent work has demonstrated that vinculin plays a role in the inside-out regulation of cadherin conformation[42]. This suggests that vinculin reinforcement could also alter friction at the AJ through regulation of the cadherin trans-dimer state, and determining relationships between vinculin state or molecular loading and this mechanism is an important point for future work. Future versions of the model could be updated to include vinculin-mediated regulation of the state of other bonds in the linkage as well as the recruitment of additional proteins to adhesion structures.

#### V. Conclusions

Collective cell migration (CCM) requires the transmission of forces between cells and to their environment. Physical models of CCM based on mechanical forces exerted between cells and at the cell-ECM interface have advanced our understanding of this process, including recent work showing that cell-cell and cell-ECM friction are major determinants of CCM dynamics[2-4]. However, these models lack insight into the molecular players that regulate key biophysical parameters. Therefore, to probe the relationship between force-activated binding dynamics and adhesion-based friction, we developed stochastic models of friction at AJs/FAs based on multi-component cadherin-/integrin-linkages with force-sensitive bond parameters from single molecule experiments. Motivated by experimental observations of the effect of vinculin mechanical reinforcement on CCM dynamics, we investigated the effect of vinculin loading on molecular friction at the FA and AJ. We found that vinculin reinforcement enhances the lifetime of cadherin-/integrin-mediated linkages under load and thereby increases the effective friction coefficient at FAs and AJs across a range of speeds corresponding to cell velocities during CCM. At higher speeds, the effective friction coefficient sharply decreases regardless of vinculin reinforcement, consistent with a distinct weak adhesion regime previously reported[2]. The effect of loaded vinculin on friction was larger at AJs (~4-fold increase in friction) than at FAs (~2-fold increase in friction). Increases in friction were associated with increases in ensemble vinculin molecular tension. Vinculin reinforcement was robust at tuning adhesion-based friction across a wide parameter space. In the context of macroscopic physical models of CCM, the effect of vinculin loading on adhesion-based friction is consistent with the observed effects of loaded vinculin on CCM dynamics in our experimental system. Overall, this work provides a framework to bridge molecular scale dynamics and regulation to macroscopic physical parameters, which could inform the future development of physical models of CCM and facilitate connections between biological and mechanical descriptions of CCM.
