## Supplementary Note 2 for "Coupling During Collective Cell Migration is Controlled by a Vinculin Mechanochemical Switch"

### Supplementary Note 2: Supporting Tables for Statistical Tests on VinTS and VinCS Data

In this supplemental note, we provide supporting tables for the statistical tests performed on experiments with VinTS or VinCS that were presented separately over more than one figure. Table S1 and Figure S1 include MDCK II VinTS and VinTS-I997A data from Fig 1 and Extended Data Fig 4 in the paper. Table S2 and Figure S2 include MDCK II and MDCK Parental VinCS data from Fig 1, Extended Data Fig 2, and Extended Data Fig 3 in the paper. Table S3 and Figure S3 include MDCK Parental VinTS, VinTS-I997A, and VinTS-Y822F data from Fig Extended Data Fig 3, Extended Data Fig 4, Extended Data Fig 5, and Extended Data Fig 6. Table S4 includes MDCK Parental VinTS S1033 mutant data from Fig 2 and Extended Data Fig 7. Table S5 includes MDCK Parental VinCS S1033 mutant data from Fig 2 and Extended Data Fig 7 and MDCK Parental VinCS data in Fig Extended Data Fig 5. See Methods section of the paper for statistical methods.

**Table S1. P-values from Steel-Dwass Test for MDCK II VinTS and VinTS-I997A Mean Eff in Fig 1 and Extended Data Fig 4**

Levine's test for unequal variance was significant, so Welch's ANOVA was conducted. P-value of Welch's ANOVA was significant, so post-hoc tests were conducted using Steel-Dwass all pairs multiple comparison. Levels are labeled as [Cell Type]_[Construct]_[Condition]_[Structure].

| Level | - Level | p-Value |
|---|---|---|
| MDCKII_VinTSI997A_Live_ApicalCyto | MDCKII_VinTSI997A_Live_AJ | 0.9831 |
| MDCKII_VinTS_Live_ApicalCyto | MDCKII_VinTS_Live_AJ | 1 |
| MDCKII_VinTSI997A_Live_FA | MDCKII_VinTSI997A_Live_AJ | 0.9961 |
| MDCKII_VinTSI997A_Live_FA | MDCKII_VinTSI997A_Live_ApicalCyto | 0.8415 |
| MDCKII_VinTS_Live_ApicalCyto | MDCKII_VinTSI997A_Live_AJ | <.0001 |
| MDCKII_VinTS_Live_AJ | MDCKII_VinTSI997A_Live_AJ | <.0001 |
| MDCKII_VinTS_Live_ApicalCyto | MDCKII_VinTSI997A_Live_ApicalCyto | <.0001 |
| MDCKII_VinTS_Live_AJ | MDCKII_VinTSI997A_Live_ApicalCyto | <.0001 |
| MDCKII_VinTS_Live_ApicalCyto | MDCKII_VinTSI997A_Live_FA | <.0001 |
| MDCKII_VinTS_Live_AJ | MDCKII_VinTSI997A_Live_FA | <.0001 |
| MDCKII_VinTS_Live_FA | MDCKII_VinTS_Live_ApicalCyto | <.0001 |
| MDCKII_VinTS_Live_FA | MDCKII_VinTS_Live_AJ | <.0001 |
| MDCKII_VinTS_Live_FA | MDCKII_VinTSI997A_Live_AJ | <.0001 |
| MDCKII_VinTS_Live_FA | MDCKII_VinTSI997A_Live_ApicalCyto | <.0001 |
| MDCKII_VinTS_Live_FA | MDCKII_VinTSI997A_Live_FA | <.0001 |

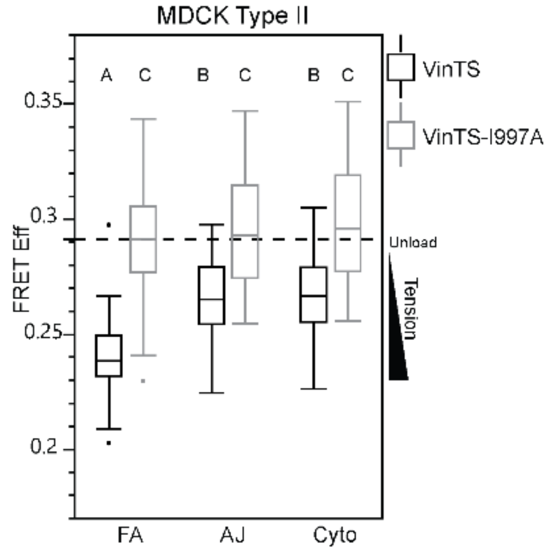


**Figure S1.** Combined box plot of MDCK II VinTS and VinTS-I997A data from Fig 1 and Extended Data Fig 4 corresponding to Table S1. Differences between groups were detected using the Steel-Dwass test. Levels not connected by the same letter are significantly different

**Table S2. P-values from Steel-Dwass Test for VinCS Mean Eff in Figures 1, Extended Data Fig 2, and Extended Data Fig 3**

Levine's test for unequal variance was significant, so Welch's ANOVA was conducted. P-value of Welch's ANOVA was significant, so post-hoc tests were conducted using Steel-Dwass all pairs multiple comparison. Levels are labeled as [Cell Type]_[Construct]_[Condition]_[Structure].

| Level | - Level | p-Value |
|---|---|---|
| pMDCK_VinCS_Live_pL | pMDCK_VinCS_Live_FA | <.0001 |
| pMDCK_VinCS_Live_pL | MDCKII_VinCS_Live_FA | <.0001 |
| pMDCK_VinCS_Live_pL | pMDCK_VinCS_Live_AJ | <.0001 |
| pMDCK_VinCS_Live_pL | MDCKII_VinCS_Live_AJ | <.0001 |
| pMDCK_VinCS_Live_pL | pMDCK_VinCS_Live_ApicalCyto | <.0001 |
| pMDCK_VinCS_Live_pL | MDCKII_VinCS_Live_ApicalCyto | <.0001 |
| pMDCK_VinCS_Live_ApicalCyto | pMDCK_VinCS_Live_FA | <.0001 |
| pMDCK_VinCS_Live_AJ | pMDCK_VinCS_Live_FA | <.0001 |
| MDCKII_VinCS_Live_ApicalCyto | MDCKII_VinCS_Live_FA | <.0001 |
| pMDCK_VinCS_Live_ApicalCyto | MDCKII_VinCS_Live_FA | <.0001 |
| MDCKII_VinCS_Live_AJ | MDCKII_VinCS_Live_FA | <.0001 |
| pMDCK_VinCS_Live_ApicalCyto | pMDCK_VinCS_Live_AJ | <.0001 |
| pMDCK_VinCS_Live_AJ | MDCKII_VinCS_Live_FA | <.0001 |
| MDCKII_VinCS_Live_ApicalCyto | MDCKII_VinCS_Live_AJ | 0.0716 |
| pMDCK_VinCS_Live_ApicalCyto | MDCKII_VinCS_Live_AJ | 0.9951 |
| pMDCK_VinCS_Live_ApicalCyto | MDCKII_VinCS_Live_ApicalCyto | 0.5277 |
| pMDCK_VinCS_Live_AJ | MDCKII_VinCS_Live_AJ | 0.0008 |
| pMDCK_VinCS_Live_AJ | MDCKII_VinCS_Live_ApicalCyto | <.0001 |
| pMDCK_VinCS_Live_FA | MDCKII_VinCS_Live_FA | <.0001 |
| pMDCK_VinCS_Live_FA | MDCKII_VinCS_Live_AJ | <.0001 |
| pMDCK_VinCS_Live_FA | MDCKII_VinCS_Live_ApicalCyto | <.0001 |

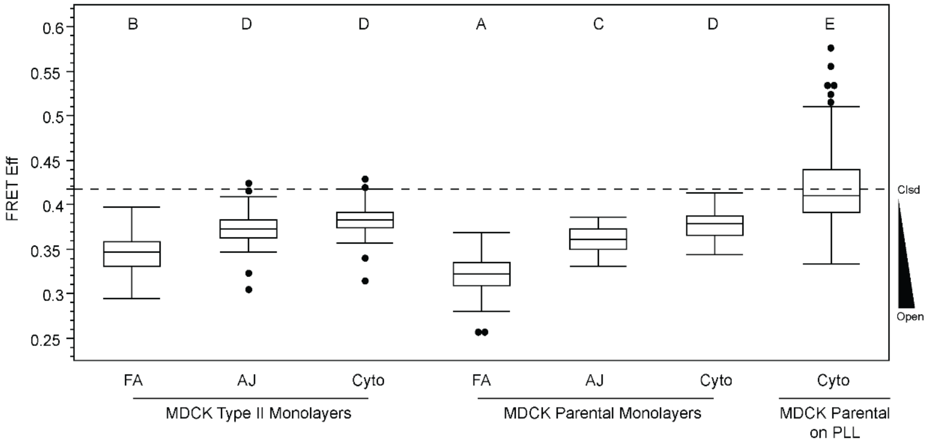


**Figure S2.** Combined box plot of MDCK II and MDCK Parental VinCS data from Fig 1, Extended Data Fig 2, and Extended Data Fig 3 corresponding to Table S2. Differences between groups were detected using the Steel-Dwass test. Levels not connected by the same letter are significantly different**.**

**Table S3. P-values from Steel-Dwass Test for Parental MDCK VinTS, VinTS-I997A, and VinTS-Y822F Mean Eff in Fig Extended Data Fig 3, Extended Data Fig 4, Extended Data Fig 5, Extended Data Fig 6**

Levine's test for unequal variance was significant, so Welch's ANOVA was conducted. P-value of Welch's ANOVA was significant, so post-hoc tests were conducted using Steel-Dwass all pairs multiple comparison. Levels are labeled as [Cell Type]_[Construct]_[Condition]_[Structure].

| Level | - Level | p-Value |
|---|---|---|
| pMDCK_VinTSY822F_Fix_ApicalCyto | pMDCK_VinTS_Fix_FA | <.0001 |
| pMDCK_VinTSY822F_Fix_ApicalCyto | pMDCK_VinTS_Fix_AJ | 0.0009 |
| pMDCK_VinTSY822F_Fix_AJ | pMDCK_VinTS_Fix_FA | 0.0039 |
| pMDCK_VinTS_Live_ApicalCyto | pMDCK_VinTS_Fix_FA | 0.0178 |
| pMDCK_VinTSY822F_Fix_ApicalCyto | pMDCK_VinTS_Live_FA | <.0001 |
| pMDCK_VinTSI997A_Live_ApicalCyto | pMDCK_VinTSI997A_Fix_FA | 0.0016 |
| pMDCK_VinTS_Fix_ApicalCyto | pMDCK_VinTS_Fix_AJ | 0.3947 |
| pMDCK_VinTSY822F_Fix_AJ | pMDCK_VinTS_Fix_AJ | 0.6251 |
| pMDCK_VinTSY822F_Fix_AJ | pMDCK_VinTS_Live_FA | 0.0517 |
| pMDCK_VinTSY822F_Fix_ApicalCyto | pMDCK_VinTS_Fix_ApicalCyto | 0.6149 |
| pMDCK_VinTS_Live_AJ | pMDCK_VinTS_Fix_FA | 0.6909 |
| pMDCK_VinTSI997A_Fix_ApicalCyto | pMDCK_VinTSI997A_Fix_AJ | 0.2442 |
| pMDCK_VinTSI997A_Live_AJ | pMDCK_VinTSI997A_Fix_FA | 0.0779 |
| pMDCK_VinTSY822F_Fix_ApicalCyto | pMDCK_VinTS_Live_AJ | 0.1259 |
| pMDCK_VinTS_Live_ApicalCyto | pMDCK_VinTS_Fix_AJ | 0.9164 |
| pMDCK_VinTSY822F_Fix_ApicalCyto | pMDCK_VinTSY822F_Fix_AJ | 0.5791 |
| pMDCK_VinTSI997A_Live_FA | pMDCK_VinTSI997A_Fix_FA | 0.703 |
| pMDCK_VinTSY822F_Fix_ApicalCyto | pMDCK_VinTS_Live_ApicalCyto | 0.823 |
| pMDCK_VinTSI997A_Live_ApicalCyto | pMDCK_VinTSI997A_Fix_AJ | 0.9945 |
| pMDCK_VinTSY822F_Fix_AJ | pMDCK_VinTS_Live_AJ | 0.9898 |
| pMDCK_VinTS_Live_ApicalCyto | pMDCK_VinTS_Live_AJ | 0.9924 |
| pMDCK_VinTSI997A_Live_ApicalCyto | pMDCK_VinTSI997A_Live_AJ | 0.9986 |
| pMDCK_VinTS_Live_AJ | pMDCK_VinTS_Fix_AJ | 1 |
| pMDCK_VinTS_Live_FA | pMDCK_VinTS_Fix_FA | 1 |
| pMDCK_VinTSY822F_Fix_AJ | pMDCK_VinTS_Fix_ApicalCyto | 1 |
| pMDCK_VinTSI997A_Live_AJ | pMDCK_VinTSI997A_Fix_AJ | 1 |
| pMDCK_VinTSY822F_Fix_AJ | pMDCK_VinTS_Live_ApicalCyto | 1 |
| pMDCK_VinTS_Live_ApicalCyto | pMDCK_VinTS_Fix_ApicalCyto | 1 |
| pMDCK_VinTSY822F_Fix_FA | pMDCK_VinTS_Live_FA | 1 |
| pMDCK_VinTSY822F_Fix_FA | pMDCK_VinTS_Fix_FA | 1 |
| pMDCK_VinTSI997A_Live_ApicalCyto | pMDCK_VinTSI997A_Fix_ApicalCyto | 1 |
| pMDCK_VinTSI997A_Live_FA | pMDCK_VinTSI997A_Live_AJ | 0.9961 |
| pMDCK_VinTSI997A_Live_FA | pMDCK_VinTSI997A_Fix_AJ | 0.9361 |
| pMDCK_VinTSI997A_Live_FA | pMDCK_VinTSI997A_Live_ApicalCyto | 0.5054 |
| pMDCK_VinTSY822F_Fix_ApicalCyto | pMDCK_VinTSI997A_Fix_FA | 0.913 |

| | | |
|---|---|---|
| pMDCK_VinTS_Live_FA | pMDCK_VinTS_Live_AJ | 0.8027 |
| pMDCK_VinTSI997A_Live_AJ | pMDCK_VinTSI997A_Fix_ApicalCyto | 0.8574 |
| pMDCK_VinTSY822F_Fix_FA | pMDCK_VinTS_Live_AJ | 0.5832 |
| pMDCK_VinTS_Live_AJ | pMDCK_VinTS_Fix_ApicalCyto | 0.9653 |
| pMDCK_VinTS_Live_FA | pMDCK_VinTS_Fix_AJ | 0.8181 |
| pMDCK_VinTSY822F_Fix_ApicalCyto | pMDCK_VinTSI997A_Live_FA | 0.0741 |
| pMDCK_VinTSY822F_Fix_FA | pMDCK_VinTS_Live_ApicalCyto | 0.0231 |
| pMDCK_VinTS_Live_FA | pMDCK_VinTS_Live_ApicalCyto | 0.0764 |
| pMDCK_VinTS_Live_ApicalCyto | pMDCK_VinTSI997A_Fix_FA | 0.028 |
| pMDCK_VinTS_Live_ApicalCyto | pMDCK_VinTSI997A_Live_FA | 0.002 |
| pMDCK_VinTSY822F_Fix_AJ | pMDCK_VinTSI997A_Live_FA | 0.0003 |
| pMDCK_VinTSY822F_Fix_FA | pMDCK_VinTSY822F_Fix_AJ | 0.0008 |
| pMDCK_VinTSY822F_Fix_ApicalCyto | pMDCK_VinTSI997A_Live_AJ | 0.0025 |
| pMDCK_VinTS_Live_ApicalCyto | pMDCK_VinTSI997A_Live_AJ | 0.0002 |
| pMDCK_VinTSY822F_Fix_AJ | pMDCK_VinTSI997A_Live_AJ | <.0001 |
| pMDCK_VinTSI997A_Live_FA | pMDCK_VinTSI997A_Fix_ApicalCyto | 0.0084 |
| pMDCK_VinTSY822F_Fix_FA | pMDCK_VinTS_Fix_AJ | 0.291 |
| pMDCK_VinTSY822F_Fix_AJ | pMDCK_VinTSI997A_Fix_FA | 0.001 |
| pMDCK_VinTS_Live_AJ | pMDCK_VinTSI997A_Live_FA | <.0001 |
| pMDCK_VinTS_Live_AJ | pMDCK_VinTSI997A_Live_AJ | <.0001 |
| pMDCK_VinTSY822F_Fix_AJ | pMDCK_VinTSI997A_Live_ApicalCyto | <.0001 |
| pMDCK_VinTSY822F_Fix_ApicalCyto | pMDCK_VinTSI997A_Live_ApicalCyto | 0.0001 |
| pMDCK_VinTS_Live_ApicalCyto | pMDCK_VinTSI997A_Live_ApicalCyto | <.0001 |
| pMDCK_VinTS_Fix_FA | pMDCK_VinTS_Fix_AJ | 0.2437 |
| pMDCK_VinTS_Live_AJ | pMDCK_VinTSI997A_Fix_FA | 0.0002 |
| pMDCK_VinTS_Live_AJ | pMDCK_VinTSI997A_Live_ApicalCyto | <.0001 |
| pMDCK_VinTSY822F_Fix_FA | pMDCK_VinTSI997A_Live_AJ | <.0001 |
| pMDCK_VinTSI997A_Fix_FA | pMDCK_VinTSI997A_Fix_AJ | 0.0005 |
| pMDCK_VinTSY822F_Fix_FA | pMDCK_VinTSI997A_Live_ApicalCyto | <.0001 |
| pMDCK_VinTSY822F_Fix_FA | pMDCK_VinTSY822F_Fix_ApicalCyto | <.0001 |
| pMDCK_VinTSY822F_Fix_FA | pMDCK_VinTSI997A_Live_FA | <.0001 |
| pMDCK_VinTS_Live_FA | pMDCK_VinTS_Fix_ApicalCyto | 0.0096 |
| pMDCK_VinTS_Live_FA | pMDCK_VinTSI997A_Live_FA | <.0001 |
| pMDCK_VinTS_Live_FA | pMDCK_VinTSI997A_Live_AJ | <.0001 |
| pMDCK_VinTSY822F_Fix_ApicalCyto | pMDCK_VinTSI997A_Fix_AJ | <.0001 |
| pMDCK_VinTS_Fix_ApicalCyto | pMDCK_VinTSI997A_Fix_FA | 0.0012 |
| pMDCK_VinTSY822F_Fix_FA | pMDCK_VinTS_Fix_ApicalCyto | 0.0011 |
| pMDCK_VinTS_Live_FA | pMDCK_VinTSI997A_Live_ApicalCyto | <.0001 |
| pMDCK_VinTSY822F_Fix_FA | pMDCK_VinTSI997A_Fix_FA | <.0001 |
| pMDCK_VinTS_Live_ApicalCyto | pMDCK_VinTSI997A_Fix_AJ | <.0001 |
| pMDCK_VinTS_Live_FA | pMDCK_VinTSI997A_Fix_FA | <.0001 |

| | | |
|---|---|---|
| pMDCK_VinTSI997A_Fix_FA | pMDCK_VinTSI997A_Fix_ApicalCyto | <.0001 |
| pMDCK_VinTS_Fix_ApicalCyto | pMDCK_VinTSI997A_Live_FA | 0.0002 |
| pMDCK_VinTSY822F_Fix_AJ | pMDCK_VinTSI997A_Fix_AJ | <.0001 |
| pMDCK_VinTS_Live_AJ | pMDCK_VinTSI997A_Fix_AJ | <.0001 |
| pMDCK_VinTSY822F_Fix_ApicalCyto | pMDCK_VinTSI997A_Fix_ApicalCyto | <.0001 |
| pMDCK_VinTS_Live_ApicalCyto | pMDCK_VinTSI997A_Fix_ApicalCyto | <.0001 |
| pMDCK_VinTS_Fix_FA | pMDCK_VinTS_Fix_ApicalCyto | <.0001 |
| pMDCK_VinTS_Live_AJ | pMDCK_VinTSI997A_Fix_ApicalCyto | <.0001 |
| pMDCK_VinTSY822F_Fix_AJ | pMDCK_VinTSI997A_Fix_ApicalCyto | <.0001 |
| pMDCK_VinTS_Fix_ApicalCyto | pMDCK_VinTSI997A_Live_AJ | <.0001 |
| pMDCK_VinTSY822F_Fix_FA | pMDCK_VinTSI997A_Fix_AJ | <.0001 |
| pMDCK_VinTSY822F_Fix_FA | pMDCK_VinTSI997A_Fix_ApicalCyto | <.0001 |
| pMDCK_VinTS_Fix_AJ | pMDCK_VinTSI997A_Live_FA | <.0001 |
| pMDCK_VinTS_Fix_AJ | pMDCK_VinTSI997A_Fix_FA | <.0001 |
| pMDCK_VinTS_Fix_ApicalCyto | pMDCK_VinTSI997A_Live_ApicalCyto | <.0001 |
| pMDCK_VinTS_Live_FA | pMDCK_VinTSI997A_Fix_AJ | <.0001 |
| pMDCK_VinTS_Fix_AJ | pMDCK_VinTSI997A_Live_AJ | <.0001 |
| pMDCK_VinTS_Live_FA | pMDCK_VinTSI997A_Fix_ApicalCyto | <.0001 |
| pMDCK_VinTS_Fix_FA | pMDCK_VinTSI997A_Live_FA | <.0001 |
| pMDCK_VinTS_Fix_AJ | pMDCK_VinTSI997A_Live_ApicalCyto | <.0001 |
| pMDCK_VinTS_Fix_FA | pMDCK_VinTSI997A_Fix_FA | <.0001 |
| pMDCK_VinTS_Fix_FA | pMDCK_VinTSI997A_Live_AJ | <.0001 |
| pMDCK_VinTS_Fix_ApicalCyto | pMDCK_VinTSI997A_Fix_AJ | <.0001 |
| pMDCK_VinTS_Fix_FA | pMDCK_VinTSI997A_Live_ApicalCyto | <.0001 |
| pMDCK_VinTS_Fix_ApicalCyto | pMDCK_VinTSI997A_Fix_ApicalCyto | <.0001 |
| pMDCK_VinTS_Fix_AJ | pMDCK_VinTSI997A_Fix_AJ | <.0001 |
| pMDCK_VinTS_Fix_FA | pMDCK_VinTSI997A_Fix_AJ | <.0001 |
| pMDCK_VinTS_Fix_AJ | pMDCK_VinTSI997A_Fix_ApicalCyto | <.0001 |
| pMDCK_VinTS_Fix_FA | pMDCK_VinTSI997A_Fix_ApicalCyto | <.0001 |

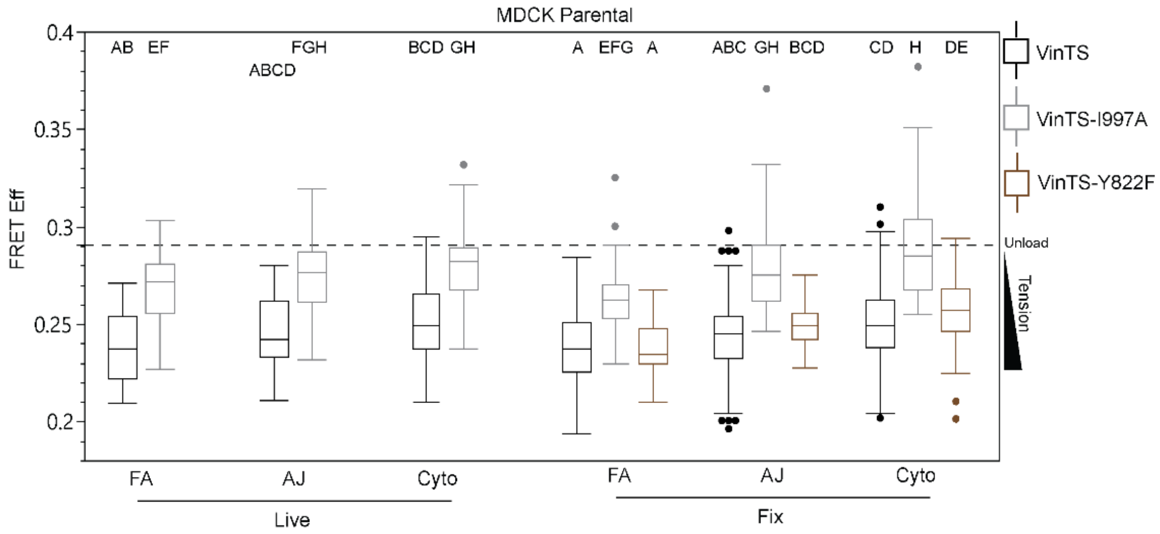


**Figure S3.** Combined box plot of MDCK Parental VinTS, VinTS-I997A, and VinTS-Y822F data from Fig Extended Data Fig 3, Extended Data Fig 4, Extended Data Fig 5, and Extended Data Fig 6 corresponding to Table S3. Differences between groups were detected using the Steel-Dwass test. Levels not connected by the same letter are significantly different.

**Table S4. P-values from Steel-Dwass Test for MDCK Parental VinTS S1033 Mutant FRET Eff in Fig 2 and Extended Data Fig 7**

Levine's test for unequal variance was significant, so Welch's ANOVA was conducted. P-value of Welch's ANOVA was significant, so post-hoc tests were conducted using Steel-Dwass all pairs multiple comparison. Levels are labeled as [Cell Type]_[Construct]_[Condition]_[Structure].

| Level | - Level | p-Value |
|---|---|---|
| pMDCK_VinTSS1033D_Fix_ApicalCyto | pMDCK_VinTS_Fix_FA | <.0001 |
| pMDCK_VinTSS1033D_Fix_AJ | pMDCK_VinTS_Fix_FA | <.0001 |
| pMDCK_VinTSS1033D_Fix_FA | pMDCK_VinTS_Fix_FA | <.0001 |
| pMDCK_VinTSS1033D_Fix_ApicalCyto | pMDCK_VinTS_Fix_AJ | <.0001 |
| pMDCK_VinTSS1033D_Fix_AJ | pMDCK_VinTS_Fix_AJ | <.0001 |
| pMDCK_VinTSS1033D_Fix_ApicalCyto | pMDCK_VinTS_Fix_ApicalCyto | <.0001 |
| pMDCK_VinTSS1033D_Fix_AJ | pMDCK_VinTS_Fix_ApicalCyto | <.0001 |
| pMDCK_VinTSS1033D_Fix_ApicalCyto | pMDCK_VinTSS1033A_Fix_AJ | <.0001 |
| pMDCK_VinTSS1033D_Fix_AJ | pMDCK_VinTSS1033A_Fix_AJ | <.0001 |
| pMDCK_VinTSS1033D_Fix_ApicalCyto | pMDCK_VinTSS1033A_Fix_ApicalCyto | <.0001 |
| pMDCK_VinTSS1033D_Fix_AJ | pMDCK_VinTSS1033A_Fix_ApicalCyto | <.0001 |
| pMDCK_VinTSS1033D_Fix_ApicalCyto | pMDCK_VinTSS1033A_Fix_FA | <.0001 |
| pMDCK_VinTSS1033D_Fix_AJ | pMDCK_VinTSS1033A_Fix_FA | <.0001 |
| pMDCK_VinTSS1033D_Fix_FA | pMDCK_VinTS_Fix_AJ | <.0001 |
| pMDCK_VinTSS1033D_Fix_FA | pMDCK_VinTSS1033A_Fix_AJ | <.0001 |
| pMDCK_VinTSS1033D_Fix_FA | pMDCK_VinTSS1033A_Fix_FA | <.0001 |
| pMDCK_VinTSS1033A_Fix_ApicalCyto | pMDCK_VinTS_Fix_FA | <.0001 |
| pMDCK_VinTSS1033D_Fix_FA | pMDCK_VinTS_Fix_ApicalCyto | <.0001 |
| pMDCK_VinTSS1033D_Fix_FA | pMDCK_VinTSS1033A_Fix_ApicalCyto | <.0001 |
| pMDCK_VinTSS1033A_Fix_AJ | pMDCK_VinTS_Fix_FA | 0.0029 |
| pMDCK_VinTSS1033A_Fix_ApicalCyto | pMDCK_VinTSS1033A_Fix_AJ | 0.4952 |
| pMDCK_VinTS_Fix_ApicalCyto | pMDCK_VinTS_Fix_AJ | 0.8893 |
| pMDCK_VinTSS1033D_Fix_ApicalCyto | pMDCK_VinTSS1033D_Fix_AJ | 0.8634 |
| pMDCK_VinTSS1033A_Fix_FA | pMDCK_VinTS_Fix_FA | 0.9974 |
| pMDCK_VinTSS1033A_Fix_ApicalCyto | pMDCK_VinTS_Fix_AJ | 0.9992 |
| pMDCK_VinTSS1033A_Fix_ApicalCyto | pMDCK_VinTS_Fix_ApicalCyto | 0.9994 |
| pMDCK_VinTSS1033A_Fix_AJ | pMDCK_VinTS_Fix_AJ | 0.7585 |
| pMDCK_VinTSS1033A_Fix_FA | pMDCK_VinTSS1033A_Fix_AJ | 0.2722 |
| pMDCK_VinTSS1033A_Fix_AJ | pMDCK_VinTS_Fix_ApicalCyto | 0.0879 |
| pMDCK_VinTSS1033A_Fix_FA | pMDCK_VinTS_Fix_AJ | 0.0062 |
| pMDCK_VinTSS1033A_Fix_FA | pMDCK_VinTSS1033A_Fix_ApicalCyto | 0.004 |
| pMDCK_VinTSS1033A_Fix_FA | pMDCK_VinTS_Fix_ApicalCyto | 0.0002 |
| pMDCK_VinTSS1033D_Fix_FA | pMDCK_VinTSS1033D_Fix_AJ | <.0001 |
| pMDCK_VinTSS1033D_Fix_FA | pMDCK_VinTSS1033D_Fix_ApicalCyto | <.0001 |
| pMDCK_VinTS_Fix_FA | pMDCK_VinTS_Fix_AJ | <.0001 |
| pMDCK_VinTS_Fix_FA | pMDCK_VinTS_Fix_ApicalCyto | <.0001 |

**Table S5. P-values from Steel-Dwass Test for MDCK Parental VinCS S1033 Mutant Norm FRET Eff Data in Fig 2 and Extended Data Fig 7 and MDCK Parental VinCS Live Norm FRET Eff Data in Fig Extended Data Fig 5**

Levine's test for unequal variance was significant, so Welch's ANOVA was conducted. P-value of Welch's ANOVA was significant, so post-hoc tests were conducted using Steel-Dwass all pairs multiple comparison. Levels are labeled as [Cell Type]_[Construct]_[Condition]_[Structure].

| Level | - Level | p-Value |
|---|---|---|
| pMDCK_VinCSS1033D_Fix_ApicalCyto | pMDCK_VinCS_Fix_FA | <.0001 |
| pMDCK_VinCSS1033D_Fix_AJ | pMDCK_VinCS_Fix_FA | <.0001 |
| pMDCK_VinCSS1033D_Fix_FA | pMDCK_VinCS_Fix_FA | <.0001 |
| pMDCK_VinCSS1033A_Fix_ApicalCyto | pMDCK_VinCS_Fix_FA | <.0001 |
| pMDCK_VinCSS1033A_Fix_AJ | pMDCK_VinCS_Fix_FA | <.0001 |
| pMDCK_VinCS_Live_ApicalCyto | pMDCK_VinCS_Fix_FA | <.0001 |
| pMDCK_VinCS_Live_AJ | pMDCK_VinCS_Fix_FA | <.0001 |
| pMDCK_VinCSS1033D_Fix_ApicalCyto | pMDCK_VinCS_Fix_AJ | <.0001 |
| pMDCK_VinCSS1033D_Fix_AJ | pMDCK_VinCS_Live_FA | <.0001 |
| pMDCK_VinCSS1033D_Fix_ApicalCyto | pMDCK_VinCS_Live_FA | <.0001 |
| pMDCK_VinCSS1033D_Fix_AJ | pMDCK_VinCS_Fix_AJ | <.0001 |
| pMDCK_VinCSS1033D_Fix_AJ | pMDCK_VinCSS1033A_Fix_FA | <.0001 |
| pMDCK_VinCSS1033D_Fix_ApicalCyto | pMDCK_VinCSS1033A_Fix_FA | <.0001 |
| pMDCK_VinCSS1033D_Fix_ApicalCyto | pMDCK_VinCS_Fix_ApicalCyto | <.0001 |
| pMDCK_VinCSS1033D_Fix_FA | pMDCK_VinCS_Live_FA | <.0001 |
| pMDCK_VinCSS1033D_Fix_AJ | pMDCK_VinCS_Fix_ApicalCyto | <.0001 |
| pMDCK_VinCSS1033A_Fix_ApicalCyto | pMDCK_VinCS_Live_FA | <.0001 |
| pMDCK_VinCSS1033D_Fix_FA | pMDCK_VinCSS1033A_Fix_FA | <.0001 |
| pMDCK_VinCSS1033A_Fix_AJ | pMDCK_VinCS_Live_FA | <.0001 |
| pMDCK_VinCSS1033A_Fix_ApicalCyto | pMDCK_VinCS_Fix_AJ | 0.0019 |
| pMDCK_VinCSS1033D_Fix_ApicalCyto | pMDCK_VinCS_Live_AJ | <.0001 |
| pMDCK_VinCSS1033D_Fix_ApicalCyto | pMDCK_VinCSS1033A_Fix_AJ | 0.0001 |
| pMDCK_VinCSS1033D_Fix_AJ | pMDCK_VinCS_Live_AJ | <.0001 |
| pMDCK_VinCSS1033D_Fix_AJ | pMDCK_VinCSS1033A_Fix_AJ | 0.0004 |
| pMDCK_VinCSS1033D_Fix_ApicalCyto | pMDCK_VinCS_Live_ApicalCyto | 0.0029 |
| pMDCK_VinCSS1033D_Fix_ApicalCyto | pMDCK_VinCSS1033A_Fix_ApicalCyto | 0.0037 |
| pMDCK_VinCS_Live_ApicalCyto | pMDCK_VinCS_Fix_AJ | 0.056 |
| pMDCK_VinCSS1033D_Fix_AJ | pMDCK_VinCSS1033A_Fix_ApicalCyto | 0.0154 |
| pMDCK_VinCSS1033D_Fix_AJ | pMDCK_VinCS_Live_ApicalCyto | 0.0132 |
| pMDCK_VinCS_Fix_ApicalCyto | pMDCK_VinCS_Fix_AJ | 0.2385 |
| pMDCK_VinCSS1033A_Fix_AJ | pMDCK_VinCS_Fix_AJ | 0.2643 |
| pMDCK_VinCSS1033A_Fix_ApicalCyto | pMDCK_VinCS_Fix_ApicalCyto | 0.2975 |
| pMDCK_VinCSS1033A_Fix_ApicalCyto | pMDCK_VinCS_Live_AJ | 0.1499 |
| pMDCK_VinCSS1033D_Fix_FA | pMDCK_VinCS_Fix_AJ | 0.8036 |
| pMDCK_VinCS_Live_ApicalCyto | pMDCK_VinCS_Fix_ApicalCyto | 0.8678 |
| pMDCK_VinCS_Live_ApicalCyto | pMDCK_VinCS_Live_AJ | 0.5196 |

| | | |
|---|---|---|
| pMDCK_VinCSS1033A_Fix_ApicalCyto | pMDCK_VinCSS1033A_Fix_AJ | 0.8365 |
| pMDCK_VinCSS1033A_Fix_AJ | pMDCK_VinCS_Live_AJ | 0.9721 |
| pMDCK_VinCSS1033D_Fix_ApicalCyto | pMDCK_VinCSS1033D_Fix_AJ | 0.9994 |
| pMDCK_VinCS_Live_AJ | pMDCK_VinCS_Fix_AJ | 0.9999 |
| pMDCK_VinCSS1033A_Fix_AJ | pMDCK_VinCS_Fix_ApicalCyto | 1 |
| pMDCK_VinCSS1033A_Fix_ApicalCyto | pMDCK_VinCS_Live_ApicalCyto | 1 |
| pMDCK_VinCSS1033D_Fix_FA | pMDCK_VinCS_Live_AJ | 1 |
| pMDCK_VinCS_Live_FA | pMDCK_VinCS_Fix_FA | 1 |
| pMDCK_VinCSS1033A_Fix_FA | pMDCK_VinCS_Live_FA | 0.9999 |
| pMDCK_VinCSS1033A_Fix_AJ | pMDCK_VinCS_Live_ApicalCyto | 0.9992 |
| pMDCK_VinCSS1033D_Fix_FA | pMDCK_VinCS_Fix_ApicalCyto | 0.9956 |
| pMDCK_VinCSS1033A_Fix_FA | pMDCK_VinCS_Fix_FA | 0.9935 |
| pMDCK_VinCS_Live_AJ | pMDCK_VinCS_Fix_ApicalCyto | 0.9836 |
| pMDCK_VinCSS1033D_Fix_FA | pMDCK_VinCSS1033A_Fix_AJ | 0.819 |
| pMDCK_VinCSS1033D_Fix_FA | pMDCK_VinCS_Live_ApicalCyto | 0.3375 |
| pMDCK_VinCSS1033D_Fix_FA | pMDCK_VinCSS1033A_Fix_ApicalCyto | 0.0145 |
| pMDCK_VinCSS1033A_Fix_FA | pMDCK_VinCS_Live_AJ | <.0001 |
| pMDCK_VinCS_Live_FA | pMDCK_VinCS_Live_AJ | <.0001 |
| pMDCK_VinCSS1033A_Fix_FA | pMDCK_VinCS_Live_ApicalCyto | <.0001 |
| pMDCK_VinCSS1033D_Fix_FA | pMDCK_VinCSS1033D_Fix_AJ | <.0001 |
| pMDCK_VinCSS1033D_Fix_FA | pMDCK_VinCSS1033D_Fix_ApicalCyto | <.0001 |
| pMDCK_VinCSS1033A_Fix_FA | pMDCK_VinCSS1033A_Fix_AJ | <.0001 |
| pMDCK_VinCSS1033A_Fix_FA | pMDCK_VinCSS1033A_Fix_ApicalCyto | <.0001 |
| pMDCK_VinCS_Live_FA | pMDCK_VinCS_Live_ApicalCyto | <.0001 |
| pMDCK_VinCSS1033A_Fix_FA | pMDCK_VinCS_Fix_AJ | <.0001 |
| pMDCK_VinCS_Live_FA | pMDCK_VinCS_Fix_AJ | <.0001 |
| pMDCK_VinCSS1033A_Fix_FA | pMDCK_VinCS_Fix_ApicalCyto | <.0001 |
| pMDCK_VinCS_Live_FA | pMDCK_VinCS_Fix_ApicalCyto | <.0001 |
| pMDCK_VinCS_Fix_FA | pMDCK_VinCS_Fix_AJ | <.0001 |
| pMDCK_VinCS_Fix_FA | pMDCK_VinCS_Fix_ApicalCyto | <.0001 |
